# Therapeutic genetic variation revealed in diverse Hsp104 homologs

**DOI:** 10.1101/2020.03.25.008748

**Authors:** Zachary M. March, Katelyn Sweeney, Hanna Kim, Xiaohui Yan, Laura M. Castellano, Meredith E. Jackrel, Edward Chuang, Edward Gomes, Karolina Michalska, Robert Jedrzejczak, Andrzej Joachimiak, Kim A. Caldwell, Guy A. Caldwell, Ophir Shalem, James Shorter

## Abstract

The AAA+ protein disaggregase, Hsp104, increases fitness under stress by reversing stress-induced protein aggregation. We have engineered potentiated Hsp104 variants to antagonize proteotoxic misfolding linked to human neurodegenerative diseases. However, these Hsp104 variants can exhibit off-target toxicity, which may limit their therapeutic utility. Hsp104 is conserved among all nonmetazoan eukaryotes, which raises the possibility that natural variants might exist with enhanced, selective activity against neurodegenerative disease substrates. To assess this possibility, we screened a cross-kingdom collection of Hsp104 homologs in several yeast proteotoxicity models. We uncovered therapeutic genetic variation among several Hsp104 homologs that specifically antagonize TDP-43 or *α*-synuclein condensate formation and toxicity in yeast, human cells, and *C. elegans*. Surprisingly, this variation manifested as increased passive chaperone activity, distinct from disaggregase activity, which neutralizes proteotoxicity of specific substrates. Thus, by exploring natural tuning of passive chaperone activity we elucidated enhanced, substrate-specific agents to counter proteotoxicity underlying neurodegenerative disease.

## INTRODUCTION

Alternative protein folding underpins fatal neurodegenerative diseases [1]. Diseases such as Parkinson’s disease (PD) and amyotrophic lateral sclerosis (ALS) have distinct clinical manifestations but are united by the prominent pathological accumulation of misfolded protein conformers [2]. The proteins implicated in each disease can adopt a range of misfolded conformations [3]. Thus, in PD, alpha-synuclein (αSyn) accumulates in toxic soluble oligomers and amyloid fibers that are the major component of cytoplasmic Lewy bodies in degenerating dopaminergic neurons [4–8]. Likewise, in ALS, the normally nuclear RNA-binding proteins, TDP-43 and FUS, accumulate in toxic oligomeric structures and cytoplasmic inclusions [9–13].

Protein disaggregation represents an innovative and appealing therapeutic strategy for the treatment of protein-misfolding disorders in that it simultaneously reverses: (a) loss-of-function phenotypes associated with sequestration of functional soluble protein into misfolded oligomers and insoluble aggregates; and (b) any toxic gain-of-function phenotypes associated with the misfolded conformers themselves [14–16]. The AAA+ (ATPases Associated with diverse Activities) protein Hsp104 from *Saccharomyces cerevisiae* (ScHsp104) rapidly disassembles a diverse range of misfolded protein conformers, including amorphous aggregates, preamyloid oligomers, and amyloid fibers [17–23]. Hsp104 assembles into asymmetric ring-shaped hexamers that undergo conformational changes upon ATP binding and hydrolysis, which drive substrate translocation across the central channel to power protein disaggregation [17, 22–27]. Each protomer is comprised of an N-terminal domain (NTD), nucleotide-binding domain 1 (NBD1), a middle domain (MD), NBD2, and a short C-terminal domain [22, 23]. Hsp104 can disassemble preamyloid oligomer and amyloid conformers of several proteins associated with neurodegenerative disease, including αSyn, polyglutamine, amyloid-beta, and tau [17, 21]. Moreover, these protein-remodeling events result in neuroprotection. For example, Hsp104 suppresses protein misfolding-induced neurodegeneration in rat and *D. melanogaster* models of polyglutamine-expansion disorders [28, 29], and a rat model Parkinson’s disease [21]. Hsp104 is the only factor known to eliminate αSyn fibers and oligomers *in vitro*, and prevent αSyn-mediated dopaminergic neurodegeneration in rats [21]. However, these activities have limits and typically high concentrations of Hsp104 are required for modest protein remodeling [17, 21, 30].

Previously, we circumvented limitations on Hsp104 disaggregase activity by developing a suite of potentiated Hsp104 variants, differing from wild-type Hsp104 by one or more missense mutations in the autoregulatory middle domain [31]. These potentiated Hsp104 variants antagonize proteotoxic misfolding of several disease-linked proteins, including TDP-43, FUS, TAF15, FUS-CHOP, EWS-FLI and αSyn [30, 32–36]. However, in some circumstances, these Hsp104 variants can also exhibit off-target toxicity [30, 32, 33]. Thus, uncovering other therapeutic Hsp104s with diminished propensity for off-target effects is a key objective [37].

Hsp104 is conserved among all non-metazoan eukaryotes [38], yet natural Hsp104 sequence space remains largely unexplored. This lack of exploration raises the possibility that natural Hsp104 sequences may exist with divergent enhanced and selective activity against neurodegenerative disease substrates. Indeed, we reported that an Hsp104 homolog from the thermophilic fungus *Calcarisporiella thermophila* antagonizes toxicity of TDP-43, αSyn, and polyglutamine in yeast without apparent toxic off-target effects [27]. These findings support our hypothesis that natural Hsp104 homologs may have therapeutically beneficial properties.

Here, we survey a cross-kingdom collection of Hsp104 homologs from diverse lineages for their ability to suppress proteotoxicity from several proteins implicated in human neurodegenerative disease. We discovered several homologs that are capable of selectively suppressing TDP-43 or αSyn toxicity. Mechanistic studies and mutational analysis suggest that these activities are due to genetic variation that impacts passive chaperone activity of Hsp104 homologs for select substrates. Thus, we establish that manipulating passive chaperone activity of Hsp104 represents a novel route to enhanced, substrate-specific agents to counter protein misfolding that causes neurodegenerative disease.

## RESULTS

### Diverse Hsp104 homologs selectively suppress TDP-43 toxicity and aggregation in yeast

In yeast, galactose-inducible expression of several proteins associated with neurodegenerative diseases, including *α*Syn [39], TDP-43 [40], and FUS [13] is toxic. However, coexpression of potentiated Hsp104 variants or the natural homolog *Calcarisporiella thermophila* Hsp104 (CtHsp104), but not ScHsp104, reduces this toxicity [27, 30, 33, 37]. For a deeper exploration of natural Hsp104 sequence space, we screened a collection of 15 additional Hsp104 homologs from diverse eukaryotes spanning ∼2 billion years of evolution and encompassing fungi (*Thielavia terrestris, Thermomyces lanuginosus, Dictyostelium discoideum, Chaetomium thermophilum, Lachancea thermotolerans, Myceliophthora thermophila, Scytalidium thermophilum,* and *Thermoascus aurantiacus*), plants (*Arabidopsis thaliana* and *Populus euphratica*), protozoa (*Monosiga brevicollis, Salpingoeca rosetta,* and *Plasmodium falciparum*), and chromista (*Chlamydomonas reinhardtii* and *Galdieria sulphuraria*) (**Figure 1A**; see **Table S1** for homolog sequences, **Figures S1** for an alignment of all homologs). All Hsp104 homologs, except the homolog from *Plasmodium falciparum (Pf)*, were non-toxic to yeast at 30 °C or 37°C (**Figure S2**). PfHsp104 was even more toxic than the potentiated Hsp104 variant, Hsp104^A503V^, at 37°C (**Figure S2**). This toxicity might reflect the very different role played by PfHsp104 in its host where it is not simply a soluble protein disaggregase [41–43]. Rather, PfHsp104 is a key component of a membrane-embedded translocon, which transports malarial proteins across a parasite-encasing vacuolar membrane into erythrocytes [41–43].

**Figure 1.**
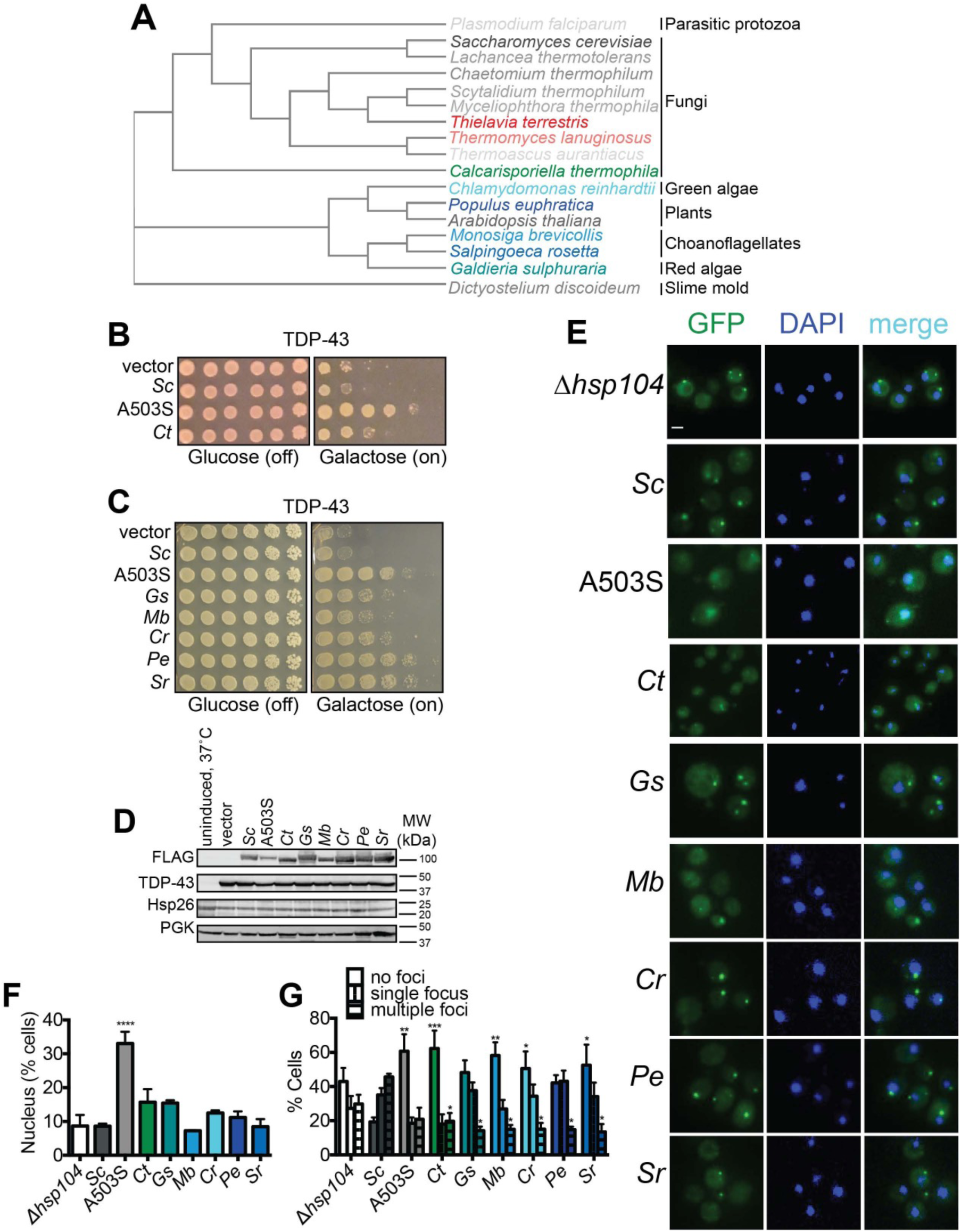
Diverse Hsp104 homologs suppress TDP-43 toxicity in yeast. **(A)** Phylogenetic tree constructed using EMBL-EBI Simple Phylogeny tool from a multiple sequence alignment of the indicated Hsp104 homologs generated in Clustal Omega (see also Supplemental Information for alignments) showing evolutionary relationships between Hsp104 homologs studied in this paper. *C. thermophila* is in green, TDP-43-specific homologs are colored in shades of blue, *α*Syn-specific homologs are colored in red, and non-rescuing homologs are colored in shades of gray. **(B)** *Δhsp104* yeast transformed with plasmids encoding galactose-inducible TDP-43 and the indicated galactose-inducible Hsp104 (either wild-type Hsp104 from *Saccharomyces cerevisiae*, the potentiated variant A503S, or the Hsp104 homolog from *Calcarisporiella thermophila* (Ct)) were serially diluted 5-fold and spotted onto glucose (expression off) or galactose (expression on). **(C)** *Δhsp104* yeast transformed with plasmids encoding galactose-inducible TDP-43 and the indicated galactose-inducible Hsp104 (either wild-type Hsp104 from *Saccharomyces cerevisiae*, the potentiated variant A503S, or homologs from *Galdieria sulphuraria* (Gs), *Monosiga brevicollis* (Mb)*, Chlamydomonas reinhardtii* (Cr)*, Populus euphratica* (Pe), and *Salpingoeca rosetta* (Sr)) were serially diluted 5-fold and spotted onto glucose (expression off) or galactose (expression on). **(D)** Western blots confirm consistent expression of FLAG-tagged Hsp104s and proteotoxic protein substrates in yeast, and that neither Hsp104 expression nor TDP-43 expression induces upregulation of the endogenous heat shock protein Hsp26. The first lane are isogenic yeast that have not been grown in galactose to induce Hsp104 and TDP-43 expression but instead have been pretreated at 37°C for 30 min to upregulate endogenous heat shock proteins. 3-phosphoglycerate kinase (PGK) is used as a loading control. Molecular weight markers are indicated (right). **(E)** Representative images of yeast co-expressing TDP-43-GFPS11 (and separately GFPS1-10 to promote GFP reassembly) and the indicated Hsp104 homologs. Cells were stained with DAPI to visualize nuclei (blue). Scale bar, 2.5µm. **(F)** Quantification of cells where TDP-43 displays nuclear localization. Values represent means ± SEM (n=3 trials with >200 cells counted per trial). One-way ANOVA with Dunnett’s multiple comparisons test was used to compare *Δhsp104* to all other conditions. ****, p<0.0001. **(G)** Quantification of cells with no, single, or multiple TDP-43 foci. Values represent means ± SEM (n=3). Two-way ANOVA with Tukey’s multiple comparisons test was used to compare the proportion of cells with either no or multiple TDP-43 foci between strains expressing different Hsp104 homologs and a control strain expressing ScHsp104. *, p<0.05; **, p<0.01; ***, p<0.001.

We screened the Hsp104 homologs for suppression of TDP-43, FUS, and *α*Syn proteotoxicity (**Figures 1B-C, 2A-B, and S3**). The toxic Hsp104 homolog, PfHsp104, was unable to reduce TDP-43, FUS, and *α*Syn proteotoxicity (**Figure S3C**). By contrast, in addition to CtHsp104, which suppresses TDP-43 and *α*Syn (**Figures 1B and 2A**), five of these new homologs (from protozoa: *Monosiga brevicollis* (*Mb*) and *Salpingoeca rosetta* (*Sr*), from chromista: *Chlamydomonas reinhardtii* (*Cr*) and *Galdieria sulphuraria* (*Gs*), and the plant *Populus euphratica* (*Pe*)), suppress TDP-43 toxicity (**Figure 1C**; see **Figure S4** for an alignment comparing TDP-43-rescuing homologs to ScHsp104). Interestingly, the Hsp104 homologs that suppress TDP-43 toxicity do not suppress *α*Syn toxicity, nor do they suppress toxicity of FUS (**Figure S3A**). Thus, we describe the first natural or engineered Hsp104 variants that diminish TDP-43 toxicity in a substrate-specific manner.

**Figure 2.**
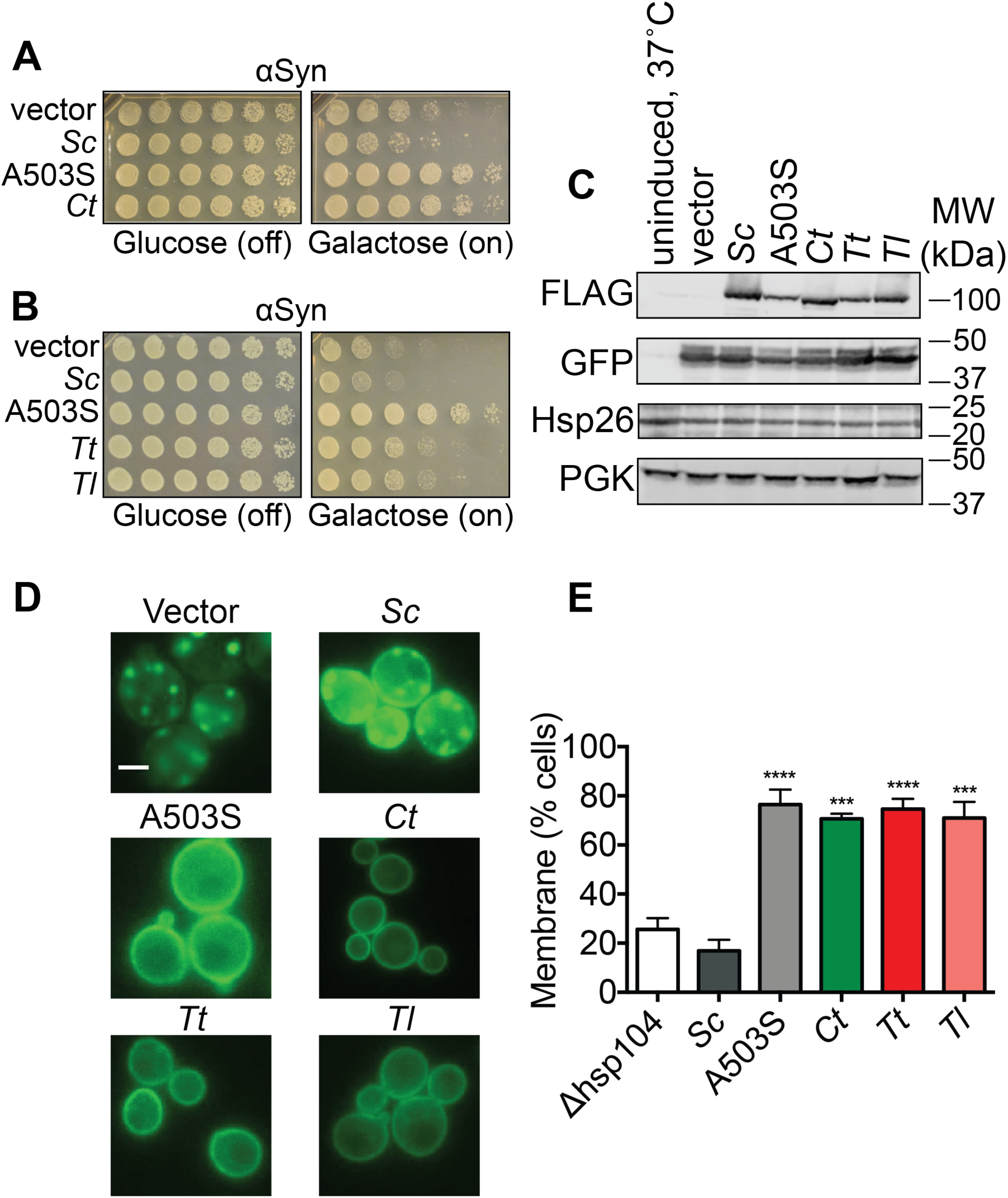
Hsp104 homologs from thermophilic fungi suppress αSyn toxicity in yeast. **(A)** *Δhsp104* yeast transformed with plasmids encoding galactose-inducible *α*Syn and the indicated galactose-inducible Hsp104 (either wild-type Hsp104 from *Saccharomyces cerevisiae*, the potentiated variant A503S, or the Hsp104 homolog from *Calcarisporiella thermophila* (Ct)) were serially diluted 5-fold and spotted onto glucose (expression off) or galactose (expression on). **(B)** *Δhsp104* yeast transformed with plasmids encoding galactose-inducible *α*Syn and the indicated galactose-inducible Hsp104 (either wild-type Hsp104 from *Saccharomyces cerevisiae*, the potentiated variant A503S, or homologs from *Calcarisporiella thermophila* (Ct), *Thielavia terrestris* (Tt), and *Thermomyces lanuginosus* (Tl)) were serially diluted 5-fold and spotted onto glucose (expression off) or galactose (expression on). **(C)** Western blots confirm consistent expression of FLAG-tagged Hsp104s and proteotoxic protein substrates in yeast, and that neither Hsp104 expression nor αSyn expression induces upregulation of the endogenous heat shock protein Hsp26. The first lane are isogenic yeast that have not been grown in galactose to induce Hsp104 and αSyn expression but instead have been pretreated at 37 °C for 30 min to upregulate heat shock proteins. 3-phosphoglycerate kinase (PGK) is used as a loading control. Molecular weight markers are indicated (right). **(D-E)** Fluorescence microscopy of cells coexpressing GFP-tagged αSyn and vector, ScHsp104^WT^, ScHsp104^A503S^, CtHsp104, TtHsp104, or TlHsp104. Scale bar, 2.5 µm. αSyn localization was quantified as the number of cells with fluorescence at the membrane or cytoplasmic inclusions. Values are means ± SEM (n=3 trials with >200 cells counted per trial). One-way ANOVA with Dunnett’s multiple comparisons test was used to compare *Δhsp104* to all other conditions. ***, p<0.001; ****, p<0.0001.

Hsp104 homologs were consistently expressed, and had minimal effect on TDP-43 levels (**Figure 1D**), indicating that suppression of toxicity was not due to reduced TDP-43 expression. Moreover, levels of the small heat-shock protein, Hsp26, were similar in all strains and lower than in control strains that had been pretreated at 37 °C to mount a heat shock response (HSR) [44]. Thus, expression of heterologous Hsp104s does not indirectly suppress TDP-43 toxicity by inducing a yeast HSR (**Figure 1D**).

Next, we examined how these Hsp104 homologs affect TDP-43 localization in our yeast model. In human cells, TDP-43 normally shuttles between the nucleus and cytoplasm, but in ALS TDP-43 becomes mislocalized to cytoplasmic inclusions [45]. Expression of GFP-tagged TDP-43 (TDP43-GFP) in yeast recapitulates this phenotype (**Figure 1E-F**) [10, 40]. Consistent with previous observations, expression of ScHsp104 does not affect TDP43-GFP cytoplasmic localization (**Figure 1F**) but slightly exacerbates formation of TDP43-GFP foci (**Figure 1G**). By contrast, the potentiated variant, Hsp104^A503S^, restores nuclear TDP-43 localization in ∼40% of cells [30, 32] (**Figure 1F**) and suppresses TDP-43 foci formation (**Figure 1F-G**). Cells expressing Hsp104 homologs that suppress TDP-43 toxicity (e.g. *Ct*, *Gs*, *Mb*, *Cr*, *Pe*, and *Sr*), show at most a modest increase in nuclear TDP43-GFP compared to control strains lacking Hsp104 (*Δhsp104*) or expressing ScHsp104 (**Figure 1F**). However, formation of cytoplasmic TDP43-GFP foci is suppressed by all homologs (**Figure 1G**). Thus, TDP-43 toxicity can be mitigated in yeast by limiting TDP-43 focus formation without restoring nuclear localization. Indeed, PeHsp104 and SrHsp104 reduced TDP-43 toxicity as effectively as Hsp104^A503S^, but without restoring TDP-43 to the nucleus (**Figure 1C, F**).

### Distinct Hsp104 homologs selectively suppress *α*Syn toxicity and inclusion formation in yeast

In addition to the five Hsp104 homologs that suppress TDP-43 toxicity, we discovered two new Hsp104 homologs (from *Thielavia terrestris* (*Tt*) and *Thermomyces lanuginosus* (*Tl*)) that suppress *α*Syn toxicity (**Figure 2B**; see **Figure S5** for an alignment comparing *α*Syn-rescuing homologs to ScHsp104 and **Figure S6** for an alignment comparing TDP-43-rescuing homologs to *α*Syn-rescuing homologs). Similar to the Hsp104s that suppress TDP-43 toxicity, these new Hsp104 homologs were selective and suppressed *α*Syn toxicity but not TDP-43 or FUS toxicity (**Figure S3B**). Eight of the fifteen Hsp104 homologs tested do not suppress TDP-43, *α*Syn, or FUS toxicity (**Figure S3C**). TtHsp104 and TlHsp104 suppressed *α*Syn toxicity without affecting *α*Syn expression and without inducing an HSR as indicated by Hsp26 levels (**Figure 2C**).

We also examined how Hsp104 homologs that suppress *α*Syn toxicity affect *α*Syn localization in yeast. *α*Syn is a lipid-binding protein that mislocalizes to cytoplasmic Lewy bodies in Parkinson’s disease [46]. Overexpression of *α*Syn in yeast recapitulates some aspects of this phenotype [39]. Indeed, *α*Syn forms toxic cytoplasmic foci that are detergent-insoluble and react with the amyloid-diagnostic dye, Thioflavin-S, and cluster cytoplasmic vesicles akin to Lewy pathology observed in PD patients [32, 39, 47–50]. We observed that all Hsp104 homologs that suppress *α*Syn toxicity also restore *α*Syn localization to the plasma membrane in ∼75% of cells (**Figure 2D-E**). Cells with membrane-localized *α*Syn showed no cytoplasmic foci (**Figure 2D-E**). Taken together, our results demonstrate that Hsp104 homologs that suppress TDP-43 or *α*Syn toxicity also suppress the formation of TDP-43 or *α*Syn inclusions.

### Sequence characteristics of Hsp104 homologs

We next sought to examine sequence relatedness among Hsp104 homologs in an effort to understand what sets Hsp104 homologs with particular toxicity-suppression characteristics apart. Hsp104 homologs with particular suppression characteristics clustered together phylogenetically (**Figure 1A**) although there are exceptions. Thus, while *A. thaliana* and *P. euphratica* are closely related species, AtHsp104 does not suppress TDP-43 while PeHsp104 does. We wondered whether Hsp104 homologs had particular sequence signatures that would predict their toxicity-suppression capacities. Multiple sequence alignments did not reveal any clear patterns to differentiate, for instance, TDP-43-suppressing Hsp104s from ScHsp104 (**Figure S4**) or from *α*Syn-suppressing ones (**Figure S6A**). We next calculated pairwise sequence identities between each possible pair of Hsp104 homologs (**Table S3**), and compared the average percent identity of pairs with similar and dissimilar toxicity suppression profiles (**Figure S6B**). The mean pairwise identity between TtHsp104 and TlHsp104, which both suppress *α*Syn toxicity, was 76%, which was unsurprising given that these homologs are from two closely related species. The ten pairwise identities comparing Hsp104 homologs that both rescue TDP-43 had a mean of 56% (with range of 51-71%), while the ten pairwise identities comparing one Hsp104 homolog that suppresses TDP-43 to another that suppresses *α*Syn had a mean of 44% (with range of 25-49%). Thus, homologs that suppress TDP-43 or *α*Syn seem to be more similar to one another than between groups.

### ClpG_GI_ from *Pseudomonas aeruginosa* reduces *α*Syn toxicity

In addition to the eukaryotic Hsp104 homologs discussed above, we also studied two prokaryotic Hsp104 homologs: ClpB from *Escherichia coli* and ClpG_GI_ from the pathogenic bacteria *Pseudomonas aeruginosa*. Wild-type ClpB does not suppress TDP-43, FUS, or *α*Syn toxicity (**Figure S7A**). We wondered whether ClpB activity could be enhanced via missense mutations in the middle domain, in analogy with Hsp104, so we also selected two previously-described hyperactive ClpB variants, ClpB^K476D^ and ClpB^Y503D^ [51–53], to test for suppression of TDP-43, FUS, and *α*Syn toxicity. Neither ClpB^K476D^ nor ClpB^Y503D^ suppressed TDP-43, FUS, or *α*Syn toxicity (**Figure S7A**). Thus, despite being able to exert forces of more than 50 pN and translocate polypeptides at speeds of more than 500 residues per second [54], neither ClpB nor hyperactive ClpB variants are capable of suppressing TDP-43, FUS or *α*Syn proteotoxicity.

Next, we tested ClpG_GI_, which bears significant homology to Hsp104 but is distinguished from other ClpB/Hsp104 proteins by an extended N-terminal domain and a shorter middle domain roughly corresponding to loss of middle domain Motif 2 [55] (**Figure S1**). ClpG_GI_ has been reported to be a more effective disaggregase than ClpB from *E. coli* [56]. Furthermore, we previously established that deleting Motif 2 from ScHsp104 yields a potentiated disaggregase [33].

Thus, we wondered whether ClpG_GI_ may suppress TDP-43, FUS, or *α*Syn toxicity. Indeed, we found ClpG_GI_ potently suppresses *α*Syn toxicity and slightly suppresses TDP-43 and FUS toxicity (**Figure S7B**). ClpB, ClpB variants, and ClpG_GI_ all express robustly in yeast and do not affect TDP-43, FUS, or *α*Syn levels (**Figure S7C**). ClpB and ClpG_GI_ are also not toxic to yeast when expressed at 37 °C (**Figure S7D**). Thus, ClpG_GI_ is a prokaryotic disaggregase with therapeutic properties and may represent a natural example of Hsp104 potentiation via loss of Motif 2 from the middle domain [33].

### Hsp104 homologs prevent TDP-43 aggregation in human cells

We next examined whether Hsp104 homologs that suppress TDP-43 toxicity in yeast would have a beneficial effect in higher model systems. Thus, we transfected human HEK293T cells with an inducible plasmid encoding fluorescently-tagged TDP-43 lacking a functional nuclear-localization sequence (mClover3-TDP43*Δ*NLS) to enhance cytoplasmic accumulation and aggregation [57]. We co-transfected these cells with an empty vector or inducible plasmids encoding ScHsp104^WT^, the potentiated variant ScHsp104^A503S^, PeHsp104, SrHsp104, or a catalytically-inactive variant ScHsp104^DPLA:DWB^ deficient in both peptide translocation (due to Y257A and Y662A mutations in NBD1 and NBD2 substrate-binding pore loops) and ATP hydrolysis (due to E285Q and E687Q mutations in NBD1 and NBD2 Walker B motifs) [17]. PeHsp104 and SrHsp104 display the most potent and selective suppression of TDP-43 toxicity in yeast (**Figure 1B**). We monitored TDP-43 expression and aggregation by pulse-shape analysis of flow cytometry data (see methods) [58] over time and quantified the percentage of cells bearing aggregates upon coexpression of each Hsp104 (**Figure 3A**). Expression of Hsp104 variants in HEK293T cells was confirmed by Western blotting (**Figure 3B**). We also monitored mClover3-TDP-43*Δ*NLS levels by Western blot, and did not observe a strong Hsp104-dependent effect on TDP-43 expression (**Figure 3B**), suggesting that Hsp104 variants are not merely affecting accumulation of TDP-43 foci by reducing TDP-43 levels. At 24 h post-transfection, all catalytically active Hsp104 variants significantly decreased the proportion of cells with TDP-43 aggregates compared to cells expressing no Hsp104 or cells expressing the catalytically-inactive ScHsp104^DPLA:DWB^ [17] (**Figure 3C**, *left panel,* Day 1). We were surprised that ScHsp104^WT^ reduced the proportion of cells with TDP-43 aggregates at this time point, given that it does not reduce TDP-43 aggregation in yeast (**Figure 1B-C**). However, at 48 h post-transfection, we observed that cells expressing the catalytically-inactive variant ScHsp104^DPLA:DWB^ or ScHsp104^WT^ both had a significantly increased proportion of cells with TDP-43 aggregates compared to cells expressing no Hsp104, while cells expressing the potentiated variant ScHsp104^A503S^ or the TDP-43 specific variant PeHsp104 continued to show a significantly lower number of cells with TDP-43 aggregates (**Figure 3C**, *right panel,* Day 2). The TDP-43-specific variant SrHsp104, meanwhile, reduced TDP-43 aggregate burden at day 1 but not day 2, suggesting an intermediate effect (**Figure 3C**). Thus, while ScHsp104 and SrHsp104 show a short-lived suppression of TDP-43 foci formation, we show two Hsp104 variants, one engineered (A503S) and one natural (PeHsp104) that show an enduring reduction of TDP-43 foci in human cells.

**Figure 3.**
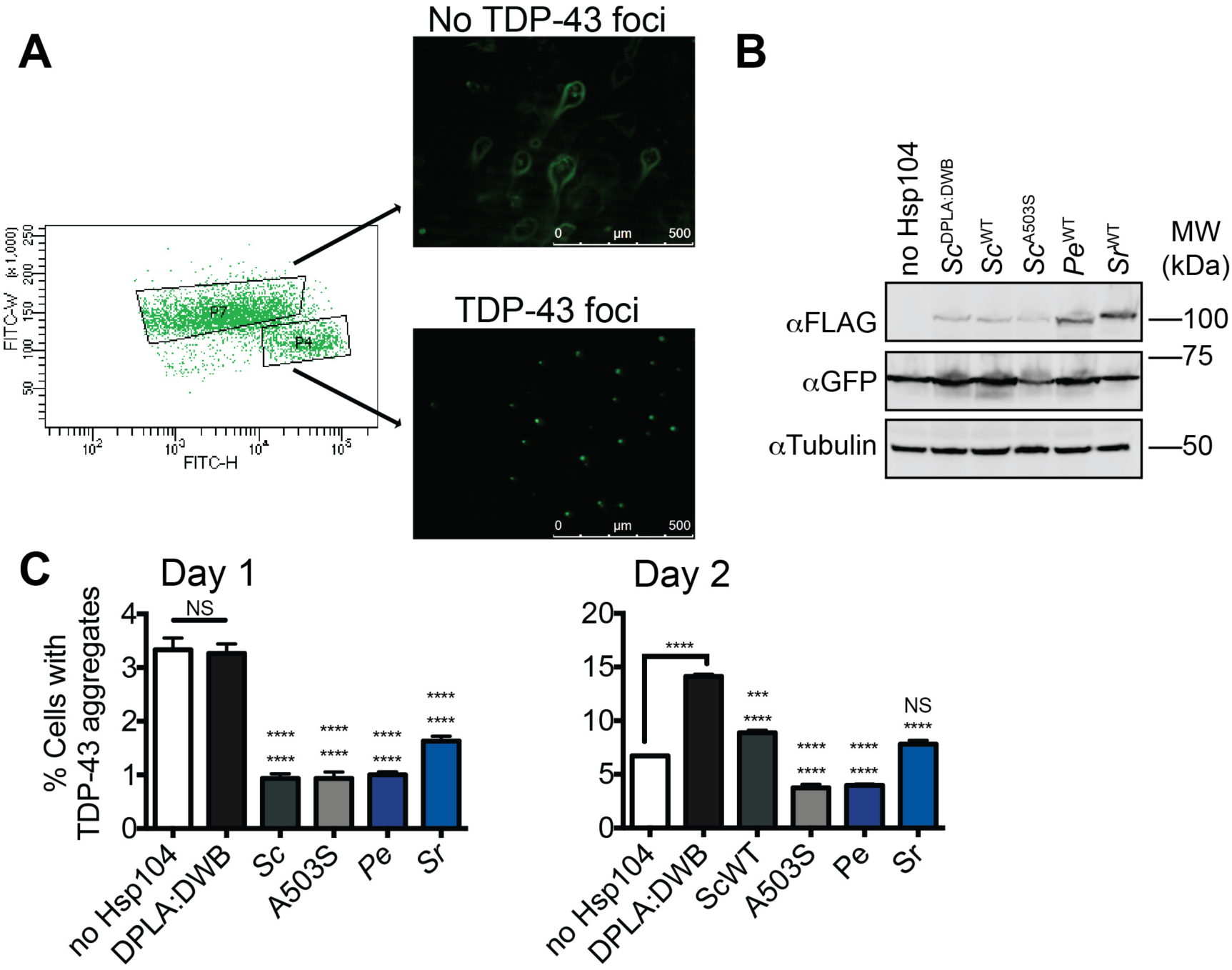
Hsp104 homologs reduce TDP-43 aggregation in HEK293T cells. **(A)** HEK293T cells were cotransfected with doxycycline-inducible constructs encoding mClover3-TDP-43*Δ*NLS. Protein expression was induced with 1 µg/ml doxycycline 6h post-transfection. At varying times, cells were sorted by FACS into populations lacking TDP-43 foci (P7) or cells with TDP-43 foci (P4). Representative fluorescent microscopy of sorted cells is shown at right. Scale bar, 500 µm. **(B)** At 24h post-transfection, cells were processed for Western blot to confirm Hsp104 expression and mClover3-TDP-43*Δ*NLS expression. **(C)** At 24h post-transfection, cells were analyzed by flow cytometry to quantify cells bearing TDP-43 aggregates (C, *left*). Cells were also analyzed by flow cytometry 48h post-transfection (C, *right*) Values are means ± SEM (n=3 independent transfections with 10,000 cells counted per trial). One-way ANOVA with Tukey’s multiple comparisons test was used to compare no Hsp104 (top set of asterisks) and DPLA:DWB (bottom set of asterisks) to all other conditions, and to each other. ***, p<0.001; ****, p<0.0001.

### αSyn-selective Hsp104 homologs prevent dopaminergic neurodegeneration in *C. elegans*

To test whether CtHsp104, TtHsp104, and TlHsp104, which suppress αSyn toxicity in yeast, would likewise suppress αSyn toxicity in animals, we turned to a *C. elegans* model of Parkinson’s disease in which the dopamine transporter (*dat-1*) promoter is used to direct expression of αSyn to dopaminergic (DA) neurons [59]. We generated transgenic worms expressing αSyn either alone or in combination with different Hsp104 variants in DA neurons, and confirmed αSyn and Hsp104 expression by qRT-PCR (**Figure 4A**). Only ∼20% of worms expressing αSyn alone have a full complement of DA neurons at day 7 post-hatching (**Figure 4B-C**). Wild-type Hsp104 from *S. cerevisiae* does not protect *C. elegans* DA neurons in this context [30]. Surprisingly, CtHsp104 also does not affect *C. elegans* DA neuron survival (**Figure 4B-C**), despite robust suppression of αSyn-mediated toxicity and inclusion formation in yeast (**Figure 2**) [27]. This lack of neuroprotection by CtHsp104 may be due to the fact that it is promiscuous. Indeed, coexpression of TtHsp104 or TlHsp104, which both selectively mitigate αSyn toxicity in yeast, both result in significant protection of DA neurons in *C. elegans* (∼40% worms with normal DA neurons in each case) after 7 days post-hatching (**Figure 4B-C**). This level of DA neuron protection is comparable to that conferred by the potentiated Hsp104 variants, Hsp104^A503S^ and Hsp104^DPLF:A503V^ [30]. Thus, our results demonstrate that natural, substrate-specific Hsp104 homologs can function in a wide variety of contexts, including in an intact animal nervous system.

**Figure 4.**
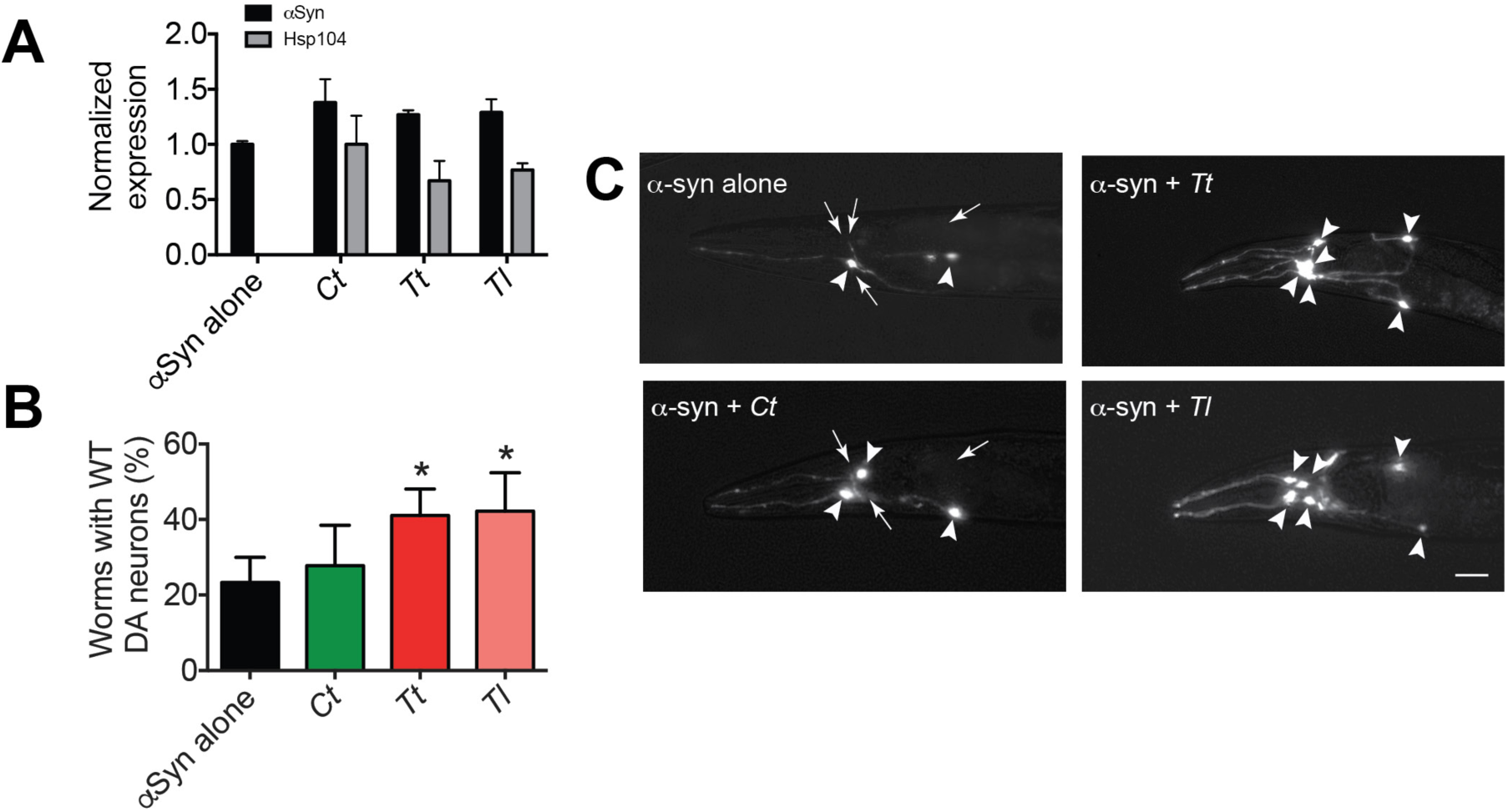
Hsp104 homologs protect against *α*Syn toxicity and dopaminergic neurodegeneration in *C. elegans*. **(A)** qRT-PCR for the expression of αSyn and various Hsp104 homologs in transgenic *C. elegans*. αSyn expression was normalized to transgenic worms overexpressing αSyn alone. Hsp104 expression was normalized to transgenic worms overexpressing both αSyn and CtHsp104. Values represent means ± SEM (N=100 worms per transgenic line, 3 independent transgenic lines examined for each genotype). The expression of either αSyn or Hsp104 homologs among all genotypes was not significantly different (one-way ANOVA, p>0.05). **(B)** αSyn and the indicated Hsp104 homolog were coexpressed in the dopaminergic (DA) neurons of *C. elegans*. Hermaphrodite nematodes have six anterior DA neurons, which were scored at day 7 posthatching. Worms are considered WT if they have all six anterior DA neurons intact (see methods for more details). TtHsp104 and TlHsp104 significantly protect over αSyn alone, while CtHsp104 had no protective effect. Values represent means ± SEM (n=30 worms per genotype per replicate, 3 independent replicates). One-way ANOVA with Dunnett’s multiple comparisons test was used to compare *α*Syn alone to all other conditions. *, p<0.05. **(C)** Photomicrographs of the anterior region of *C. elegans* coexpressing GFP with αSyn. Worms expressing αSyn alone (top left) exhibit an age-dependent loss of DA neurons. Worms expressing αSyn plus either *Ct* (bottom left) *Tt* (top right) or *Tl* (bottom right) exhibit greater neuronal integrity. Arrows indicate degenerating or missing neurons. Arrowheads indicate normal neurons. Scale bar, 10 µm.

### Differential suppression of proteotoxicity by Hsp104 homologs is not due to changes in disaggregase activity

Next, we sought to understand why some Hsp104 homologs suppress TDP-43 or *α*Syn toxicity while others do not. One possible explanation is that Hsp104 homologs differ in disaggregase activity, as has been the case with potentiated Hsp104 variants [30, 32–34, 37, 60]. Potentiated disaggregases typically display elevated ATPase and disaggregase activities. This elevated activity can sometimes manifest as a temperature-dependent toxicity phenotype [30, 33]. However, the Hsp104 homologs we assess here (except for PfHsp104) are non-toxic under conditions where some potentiated Hsp104 variants, such as Hsp104^A503V^, are toxic (**Figure S2**). This finding hints that these natural homologs are not potentiated in the same way as engineered variants. To explore this issue further, we directly assessed whether the toxicity-suppression behavior of Hsp104 homologs could be rationalized by differences in disaggregase activity.

First, we tested how well our Hsp104 homologs conferred thermotolerance to yeast. Hsp104 is an essential factor for induced thermotolerance in yeast [61], and Hsp104 homologs in bacteria and plants have similar functions in their respective hosts [62, 63]. The ability of Hsp104 to confer thermotolerance depends on its disaggregase activity, which solubilizes proteins trapped in heat-induced protein assemblies [18, 64–67]. Thus, thermotolerance is a convenient *in vivo* proxy for disaggregase activity among different Hsp104 homologs. Indeed, ∼75% of wild-type yeast survive a 20-min heat shock at 50°C whereas ∼1% of *Δhsp104* mutants survive the same shock (**Figure 5A-B**). Expressing FLAG-tagged *Sc*Hsp104 from a plasmid effectively complements the thermotolerance defect of *Δhsp104* yeast (**Figure 5A-B**). We generated transgenic yeast strains in which Hsp104 homologs are expressed under the control of the native *S. cerevisiae HSP104* promoter, and assessed the thermotolerance phenotypes of these strains. We observed a range of phenotypes. Specifically, 15 of 17 Hsp104s tested conferred some degree of thermotolerance above *Δhsp104* alone (**Figure 5A-B**). The two exceptions were ClpB from *Escherichia coli* and Hsp104 from *Populus euphratica* (**Figure 5A-B**). Some homologs, such as those from *Thielavia terrestris, Galdieria sulphuraria*, and *Dictyostelium discoideum* strongly complement thermotolerance while others were quite weak (**Figure 5A-B**). These differences are not due to differences in expression, as all homologs were consistently expressed, as confirmed by Western blot (**Figure 5C**).

**Figure 5.**
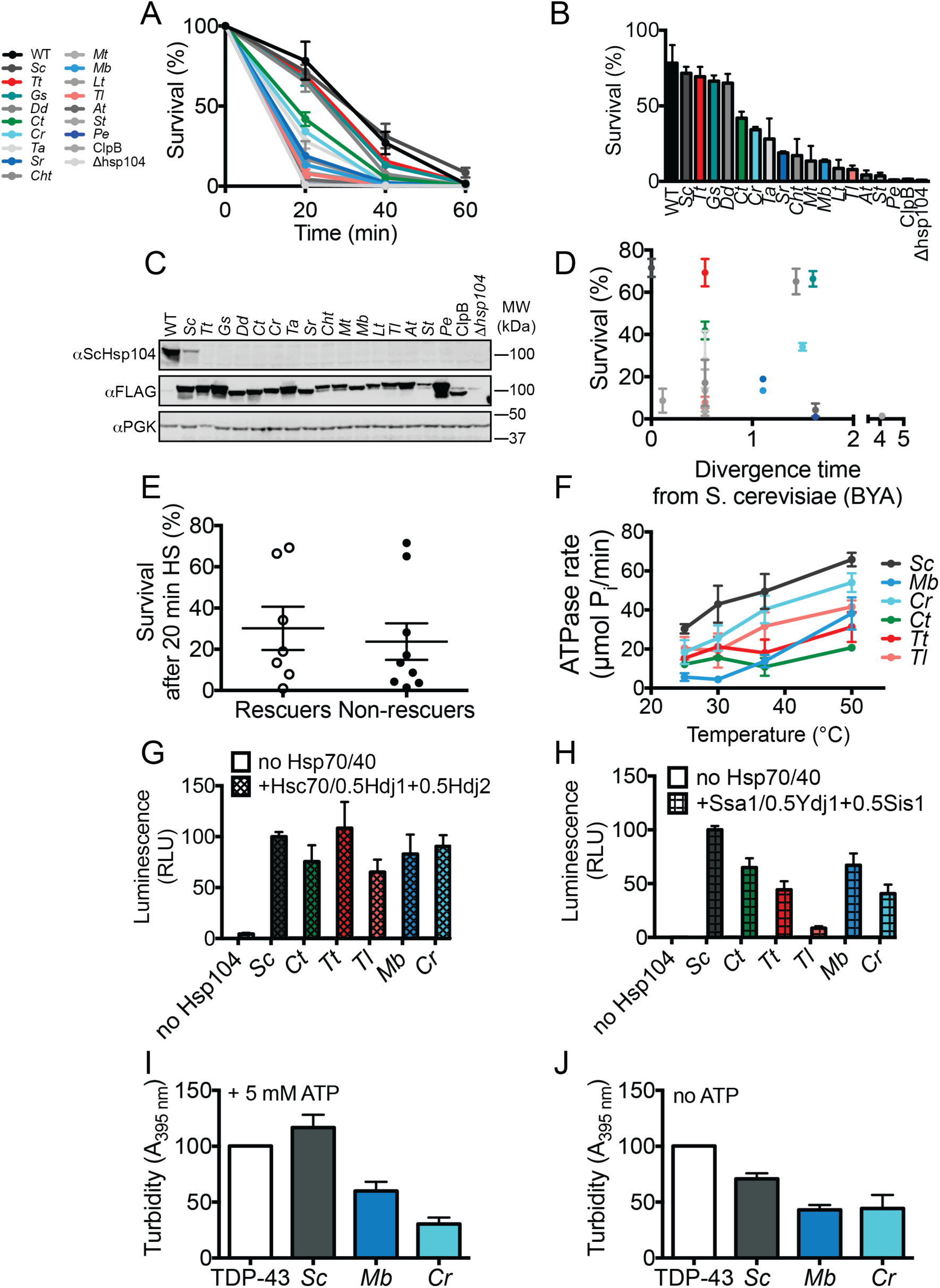
Hsp104 homologs function in induced thermotolerance and are ATPase-driven disaggregases, but differences in disaggregase activity do not explain suppression of TDP-43 or αSyn toxicity. **(A)** WT or *Δhsp104* yeast carrying a plasmid encoding the indicated Hsp104 homolog under the control of the native *HSP104* promoter were pre-treated at 37°C for 30 min, treated at 50 °C for 0-60 min, and plated. Surviving colonies were quantified after 2d recovery. Values represent means ±SEM (n=3 independent transformations). **(B)** Hsp104s ranked by thermotolerance performance after a 20 min heat shock at 50°C. **(C)** Expression of Hsp104 homologs from the *HSP104* promoter was confirmed by Western blotting. Molecular weight markers are indicated (right). **(D)** Survival after 20 min heat shock does not track with the evolutionary separation between a given species and *S. cerevisiae*. **(E)** Thermotolerance activity of Hsp104 homologs that suppress TDP-43 or αSyn does not noticeably differ from Hsp104 homologs that do not suppress TDP-43 or αSyn toxicity. **(F)** ATPase activity of the indicated Hsp104 homologs. Values represent means ± SEM (n=3). **(G)** Luciferase aggregates (50 nM) were incubated with the indicated Hsp104 (0.167 µM hexamer) with or without 0.167 µM Ssa1, 0.073 µM Ydj1, and 0.073 µM Sis1 for 90 min at 25°C. Values represent means ± SEM (n=3). **(H)** Luciferase aggregates were treated as in (B) but Ssa1, Ydj1, and Sis1 were replaced with Hsc70, Hdj1, and Hdj2. Values represent means ± SEM (n=3). **(I)** TDP-43 (3 µM) was incubated in the presence of the indicated Hsp104 (6 µM) and 5 mM ATP, and turbidity was measured at 3h relative to TDP-43 aggregation reactions containing no Hsp104. Values represent means +/-SEM (n=3). **(J)** As in (I), except ATP was omitted and turbidity was measured at 2h.

Thermotolerance phenotypes are not explained by the evolutionary divergence time of a particular species from *S. cerevisiae* (**Figure 5D**). Likewise, thermotolerance was not noticeably different between Hsp104 homologs that reduced TDP-43 or *α*Syn toxicity compared to those that did not (**Figure 5E**). Interestingly, ClpG_GI_ confers a strong thermotolerance phenotype to *Δhsp104* yeast (**Figure S7E**) while ClpB does not, although both are from prokaryotes. This difference may be due to the fact that ClpG_GI_ is a stand-alone disaggregase and does not depend on Hsp70 and Hsp40 for disaggregation [55, 56] while ClpB is incompatible with yeast Hsp70 and Hsp40 [68]. Nevertheless, our results cumulatively demonstrate that evolutionarily diverse Hsp104s confer thermotolerance to *Δhsp104* yeast, but differences in thermotolerance activity do not explain differences in suppression of TDP-43 or *α*Syn toxicity.

We next expressed and purified several Hsp104 homologs (MbHsp104, CrHsp104, TtHsp104, TlHsp104, ScHsp104, and CtHsp104) to define their biochemical properties. All Hsp104 homologs are active ATPases, and display increased ATPase activity with increasing temperature (**Figure 5F**). These findings are consistent with the fact that Hsp104 is a disaggregase induced by thermal stress. We also investigated the disaggregase activity of Hsp104 homologs by assessing their ability to disaggregate disordered luciferase aggregates *in vitro* [17, 18]. Each Hsp104 homolog was able to disassemble and reactivate aggregated luciferase (**Figure 5G-H**). Luciferase disaggregation required the presence of Hsp70 and Hsp40 chaperones, which could be from yeast (Ssa1, Sis1, and Ydj1 **Figure 5H**) or human (Hsc70, Hdj1, and Hdj2 **Figure 5G**). We did not observe differences between luciferase disaggregation activity of Hsp104 homologs that suppress TDP-43 toxicity versus those that suppress *α*Syn (**Figure 5G-H**).

Next, we tested the activity of several Hsp104 homologs (ScHsp104, CtHsp104, TtHsp104, AtHsp104, and MtHsp104) against an ordered amyloid substrate, semen-derived enhancer of viral infection (SEVI) [69]. As previously reported [27, 53], all Hsp104 homologs tested rapidly remodeled SEVI fibrils (**Figure S8A**). Electron microscopy revealed that Hsp104 homologs remodeled SEVI fibrils into small, amorphous structures (**Figure S8B**). Thus, eukaryotic Hsp104 homologs are generally able to remodel amyloid fibrils. However, there was not an obvious difference in amyloid-remodeling activity between homologs that also suppress *α*Syn toxicity (CtHsp104 and TtHsp104) and those that do not (ScHsp104, AtHsp104, and MtHsp104). Taken together, these findings suggest that differences in proteotoxicity suppression by Hsp104 homologs is not simply due to differences in general disaggregase activity of Hsp104 homologs.

### Hsp104 homologs inhibit protein aggregation in an ATP-independent manner

Since differences in general disaggregase activity do not explain differences in proteotoxicity suppression among Hsp104 homologs, we wondered whether Hsp104 homologs may act instead to inhibit protein aggregation. To test this possibility, we reconstituted TDP-43 aggregation *in vitro* to test how Hsp104 homologs affect TDP-43 aggregation. We performed reactions in the presence (**Figure 5I**) or absence (**Figure 5J**) of ATP. Hsp104 from *Monosiga brevicollis* and *Chlamydomonas reinhardtii*, which selectively suppress TDP-43 toxicity and foci formation in yeast, inhibit TDP-43 aggregation *in vitro,* whereas Hsp104 from *Saccharomyces cerevisiae* has no effect (**Figure 5I-J**). Hsp104-mediated inhibition of TDP-43 aggregation occurred in the presence or absence of ATP (**Figure 5I-J**). Hsp104 homologs did not inhibit TDP-43 aggregation merely by inhibiting cleavage of the MBP tag by TEV protease (**Figure S9**). Thus, unexpectedly, specific Hsp104 homologs appear to suppress TDP-43 toxicity by inhibiting its aggregation in an ATP-independent manner.

### ATPase-independent passive chaperone activity underlies suppression of TDP-43 and *α*Syn toxicity by Hsp104 homologs

Since we determined that Hsp104 homologs engage in passive chaperone activity to inhibit TDP-43 aggregation *in vitro*, we next tested whether or not this reflected the predominant mechanism by which Hsp104 homologs suppress TDP-43 or *α*Syn toxicity in yeast. Thus, we generated a series of mutants for each Hsp104 homolog intended to disrupt their disaggregase activity. Hsp104 disaggregase activity is driven by: (1) ATP binding and hydrolysis, which are mediated by Walker A and Walker B motifs, respectively, and which drive conformational changes within the hexamer to support substrate translocation [17, 25, 66], and (2) substrate translocation through the central pore of the hexamer, which is mediated by tyrosine-bearing pore loops [17, 24, 70, 71]. These motifs are highly conserved among all Hsp104 homologs (**Figures 6A and S1**). We mutated: (1) conserved lysine residues in the Walker A motifs to either alanine or threonine (Hsp104^DWA(KA)^ and Hsp104^DWA(KT)^) in NBD1 and NBD2 to impair ATP binding, (2) conserved glutamate residues in the Walker B motifs to either alanine or glutamine (e.g. Hsp104^DWB(EA)^ or Hsp104^DWB(EQ)^) in NBD1 and NBD2 to impair ATP hydrolysis, and (3) conserved tyrosines in the pore loops to alanine (e.g. Hsp104^DPLA^) in NBD1 and NBD2 to impair substrate threading through the central hexamer pore [17]. We also generated mutants lacking the NTD, which also plays a role in substrate binding and processing [35, 72]. We monitored expression of Hsp104 mutant proteins by Western blotting, and found all mutants were expressed similarly to wild-type protein, from either the galactose or native *HSP104* promoter (**Figure S10A-H**). Strains expressing Walker A, Walker B, or pore-loop mutant proteins are all severely impaired in thermotolerance compared to wild-type controls, while *Δ*N mutants are only mildly impaired in thermotolerance compared to wild-type proteins (**Figure S10I**). The only exception is PeHsp104, where the wild-type protein confers no thermotolerance benefit over *Δhsp104* cells, which precludes any conclusions being drawn about the impact of mutations on this protein (**Figure 5B, S10I**). These findings are similar to thermotolerance phenotypes of ScHsp104 mutants that have been previously reported [17, 66, 72]. Thus, we confirm that these mutations have a conserved effect on Hsp104 disaggregase activity.

**Figure 6.**
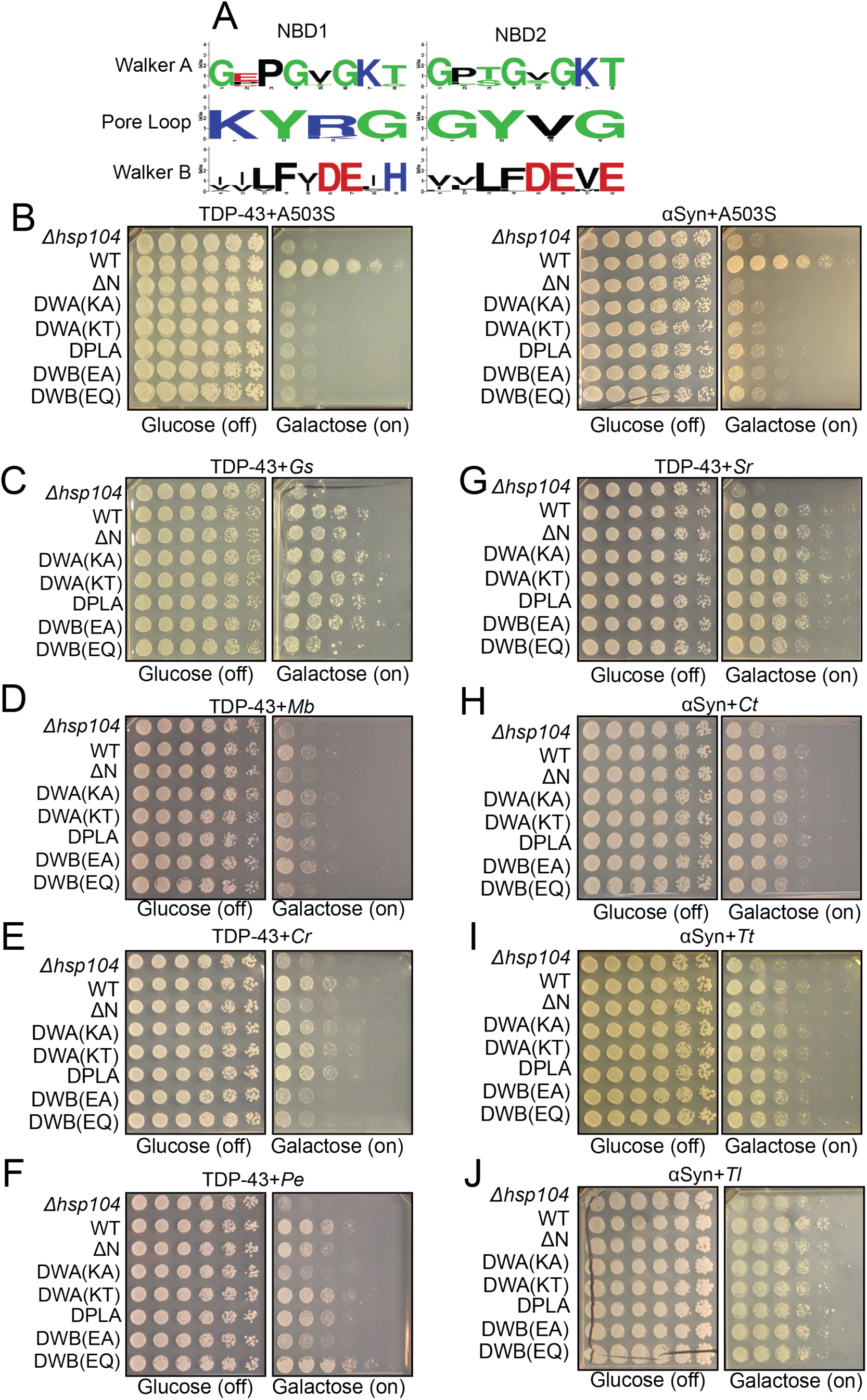
Genetic variation within Hsp104 homologs enables their disaggregase-independent suppression of TDP-43 or *α*Syn toxicity. **(A)** WebLogo sequence logos demonstrating high conservation of Walker A, tyrosine-bearing pore loops, and Walker B motifs in both NBD1 and NBD2 across all Hsp104 homologs. **(B)** Spotting assay demonstrating that suppression of TDP-43 (*left*) and *α*Syn (*right*) toxicity by ScHsp104^A503S^ (A503S) is inhibited by mutations in Walker A (DWA), pore loop (DPLA), and Walker B (DWB) motifs. **(C-G)** Spotting assays demonstrate that suppression of TDP-43 toxicity by GsHsp104 (C), MbHsp104, (D), CrHsp104 (E), PeHsp104 (F), and SrHsp104 (G) is resistant to multiple mutations that impair ATP-driven disaggregase activity. **(H-J)** Spotting assays demonstrate that suppression of *α*Syn toxicity by CtHsp104 (H), TtHsp104 (I), and TlHsp104 (J) is resistant to multiple mutations that impair ATP-driven disaggregase activity.

Next, we examined how these mutations affect the ability of different Hsp104 homologs to reduce TDP-43 or *α*Syn toxicity in yeast. TDP-43 or *α*Syn expression levels were unaffected by Hsp104 mutants, as assessed by Western blot (**Figure S10A-H**). All of the aforementioned mutations strongly impair the ability of a potentiated Hsp104 variant, ScHsp104^A503S^, to suppress TDP-43 and *α*Syn toxicity [60, 72] (**Figure 6B**). Thus, ScHsp104^A503S^ suppresses TDP-43 toxicity by a disaggregase-mediated mechanism that necessitates ATP binding and hydrolysis and substrate-engagement by conserved pore-loop tyrosines [60]. Remarkably, however, Hsp104 homolog-mediated suppression of TDP-43 or *α*Syn toxicity was largely unaffected by these mutations (**Figure 6C-J**). For example, mutation of conserved pore-loop tyrosines to alanine had no effect on suppression of TDP-43 or *α*Syn toxicity (**Figure 6C-J**). Thus, canonical substrate translocation is not required to mitigate TDP-43 or *α*Syn toxicity. Likewise, in most cases, mutation of Walker A or Walker B motifs did not affect suppression of TDP-43 or *α*Syn toxicity (**Figure 6C-J**). There were, however, some exceptions: CrHsp104 which was inhibited by mutations in the Walker B motifs (**Figure 6E**), and PeHsp104, which was modestly inhibited by K to A but not K to T substitutions in the Walker A motif and E to A but not E to Q substitutions in the Walker B motifs (**Figure 6F**). Thus, CrHsp104 and PeHsp104 likely require ATPase activity for optimal suppression of toxicity, but this requirement was not coupled to substrate translocation by conserved pore-loop tyrosines. Nevertheless, the remaining Hsp104 homologs tested here (GsHp104, MbHsp104. SrHsp104, CtHsp104, TtHsp104, and TlHsp104) suppressed TDP-43 or *α*Syn toxicity by a mechanism that does not require ATPase activity or conserved pore-loop tyrosines that engage substrate in the Hsp104 channel. Mechanistically, these findings suggest that toxicity suppression by the Hsp104 homologs is primarily due to ATPase-independent chaperoning of specific substrates, such as TDP-43 or *α*Syn, and not due to disaggregation or ATPase-dependent chaperone activity.

### Suppression of TDP-43 toxicity by MbHsp104 and CrHsp104, and suppression of *α*Syn toxicity by TtHsp104 is mediated by the NTD

Next, we sought to identify the requisite architecture of Hsp104 homologs that enables TDP-43 or *α*Syn proteotoxicity suppression. The engineered ScHsp104 variant, Hsp104^A503S^, requires the NTD to mitigate TDP-43 and *α*Syn toxicity (**Figure 6B**) [72]. TDP-43 or *α*Syn proteotoxicity suppression by several Hsp104 homologs (GsHsp104, PeHsp104, SrHsp104, CtHsp104, and TlHsp104) is unaffected by NTD deletion (**Figure 6C, F, G, H, J**). Interestingly, MbHsp104^ΔN^ and CrHsp104^ΔN^ are unable to suppress TDP-43 toxicity (**Figure 6D-E**), and TtHsp104^ΔN^ is unable to suppress *α*Syn toxicity (**Figure 6I**). Thus, proteotoxicity suppression by these Hsp104 homologs appears to be mediated by their NTDs.

We wondered whether the NTDs might drive the toxicity suppression phenotypes for MbHsp104, CrHsp104, and TtHsp104. To test this possibility, we made a series of chimeras in which we systematically replaced domains of ScHsp104 with the homologous domains from other Hsp104 homologs (see **Figure 7A** for illustration of domain boundaries and chimeras). First, we found that replacing the NTD of ScHsp104 with the NTD of either MbHsp104 or CrHsp104 did not enable suppression of TDP-43 toxicity by the resulting chimeras (**Figures 7B-C**). Similarly, replacing the ScHsp104 NTD with the NTD from TtHsp104 did not enable suppression of *α*Syn toxicity by the resulting chimera (**Figure 7D**). Thus, proteotoxicity suppression by MbHsp104, CrHsp104, and TtHsp104 is not simply transmitted through or encoded by the NTD. Rather, the NTD must work together with neighboring domains of the same Hsp104 homolog to enable toxicity suppression.

**Figure 7.**
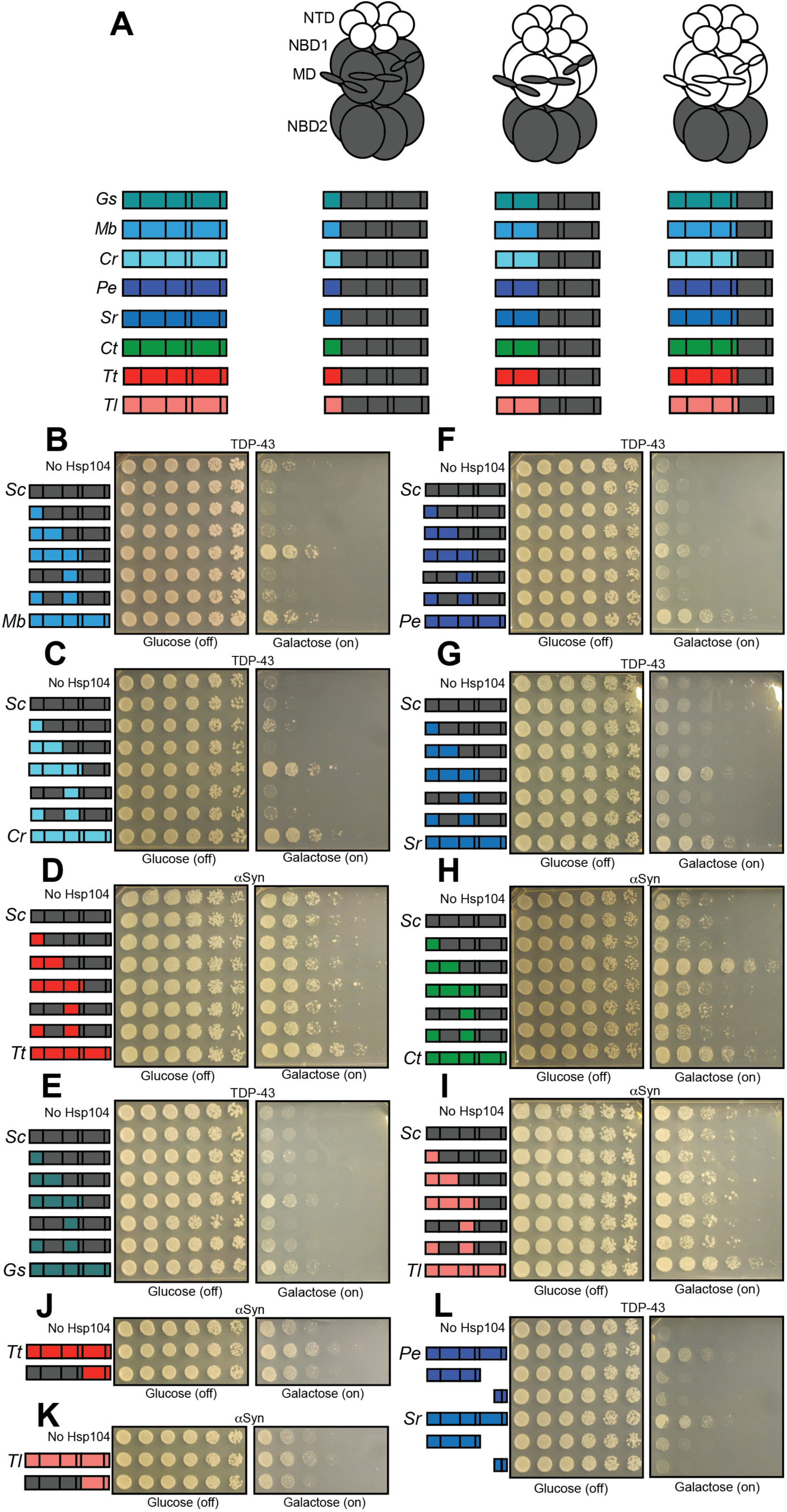
Interactions between the NTD, NBD1, and MD support proteotoxicity suppression by Hsp104 homologs. **(A)** Color codes and domain boundaries and labels of Hsp104 homologs. **(B)** Spotting assay of *Δhsp104* yeast coexpressing TDP-43 and the indicated chimeric Hsp104s between ScHsp104 and MbHsp104 illustrates that chimeras possessing the NTD, NBD1, and MD from MbHsp104 copy the TDP-43 toxicity suppression phenotype of MbHsp104. **(C)** Spotting assay of *Δhsp104* yeast coexpressing TDP-43 and the indicated chimeric Hsp104 between ScHsp104 and CrHsp104 illustrates that chimeras possessing the NTD, NBD1, and MD from CrHsp104 copy the TDP-43 toxicity suppression phenotype of CrHsp104. **(D)** Spotting assay of *Δhsp104* strains coexpressing *α*Syn and the indicated chimeric Hsp104 between ScHsp104 and TtHsp104 illustrates that no chimeras between ScHsp104 and TtHsp104 replicate the *α*Syn toxicity-suppressing phenotype of TtHsp104. **(E)** Spotting assay of *Δhsp104* yeast coexpressing TDP-43 and the indicated chimeric Hsp104 between ScHsp104 and GsHsp104 illustrates that chimeras possessing the NTD, NBD1, and MD from GsHsp104 copy the TDP-43 toxicity suppression phenotype of GsHsp104. **(F)** Spotting assay of *Δhsp104* yeast coexpressing TDP-43 and the indicated chimeric Hsp104 between ScHsp104 and PeHsp104 illustrates that chimeras possessing the NTD, NBD1, and MD from PeHsp104 copy the TDP-43 toxicity suppression phenotype of PeHsp104. **(G)** Spotting assay of *Δhsp104* yeast coexpressing TDP-43 and the indicated chimeric Hsp104 between ScHsp104 and SrHsp104 illustrates that chimeras possessing the NTD, NBD1, and MD from SrHsp104 copy the TDP-43 toxicity suppression phenotype of SrHsp104. **(H)** Spotting assay of *Δhsp104* strains coexpressing the indicated chimeric Hsp104 and *α*Syn illustrates that chimeras possessing the NTD and NBD1 from CtHsp104 copies the *α*Syn toxicity-suppressing phenotype of CtHsp104^WT^. **(I)** Spotting assay of *Δhsp104* strains coexpressing *α*Syn and the indicated chimeric Hsp104 between ScHsp104 and TlHsp104 illustrates that no chimeras between ScHsp104 and TlHsp104 replicate the *α*Syn toxicity-suppressing phenotype of TlHsp104. **(J-K)** Spotting assays of *Δhsp104* strains coexpressing the indicated chimeric Hsp104 and *α*Syn illustrates that NBD2 from TtHsp104 **(J)** or TlHsp104 **(K)** is not sufficient to copy the *α*Syn toxicity-suppression phenotype of TtHsp104 or TlHsp104. **(L)** Spotting assay of *Δhsp104* strains coexpressing TDP-43 and either full-length PeHsp104 or SrHsp104, or monomeric fragments derived from these homologs demonstrates that Hsp104-mediated toxicity suppression is an emergent property of hexameric Hsp104.

### Suppression of TDP-43 toxicity is enabled by NBD1 and MD residues in Hsp104 homologs

Based on our observation that suppression of TDP-43 and *α*Syn toxicity by MbHsp104, CrHsp104, and TtHsp104 is not conferred solely by the NTD (**Figure 7B-D**), and the observation that *Δ*N mutants of other homologs (GsHsp104, PeHsp104, SrHsp104, CtHsp104, and TlHsp104) retained their ability to reduce either TDP-43 or *α*Syn toxicity (**Figure 6C, F, G, H, J**), we reasoned that residues in other domains must also contribute to substrate selectivity. Within the ScHsp104 hexamer, the NTD interacts with NBD1 and the MD [35]. Thus, we made a series of additional chimeras by progressively replacing the NTD, NBD1 or MD of ScHsp104 for the homologous domain from another Hsp104 homolog (see **Figure 7A** for illustration). Generally, the chimeras expressed well in yeast (**Figure S11**). We observed that chimeras consisting of the NTD, NBD1, and MD from a TDP-43-selective homolog appended to the NBD2 and CTD from ScHsp104 phenocopied the TDP-43 suppression phenotype associated with the homolog itself (**Figure 7B, C, E, F, G**). These same chimeras do not reduce *α*Syn toxicity (**Figure S12A-E**). Thus, these chimeras encompass the sequence determinants that are crucial for passive chaperoning of TDP-43. Chimeras where cognate NTD:NBD1:MD networks were disrupted do not reduce TDP-43 toxicity (**Figure 7B, C, E, F, G**). We conclude that genetic variation in NBD1 and the MD of Hsp104 homologs also contributes to mitigation of TDP-43 proteotoxicity.

### Non-cognate NTD:NBD1 units can yield toxic chimeras at 37°C

We next assessed the fitness effects intrinsic to the chimeras. We expressed all chimeras in *Δhsp104* yeast at 37 °C in the absence of any toxic substrate protein to observe any intrinsic toxicity associated with the chimeras themselves. We observed that NTD replacement alone does not cause toxicity (**Figure S13A-H**, third row in all panels). By contrast, replacing both the NTD and NBD1 resulted in toxicity in several cases (e.g. for GsHsp104, CrHsp104, PeHsp104, SrHsp104, and CtHsp104; **Figure S13A, C-F**, *boxed*), although TtHsp104 and TlHsp104 (*α*Syn-specific variants) were exceptions **Figure S13G, H**). Interestingly, no toxicity was observed in chimeras where the NTD, NBD1, and MD were replaced together (**Figure S13A-H**, fifth row), nor in chimeras where the MD was replaced alone (**Figure S13A-H**, sixth row) or in combination with the NTD (**Figure S13A-H**, seventh row). These findings suggest that altered non-cognate interactions between the NTD:NBD1 unit and the MD can elicit off-target toxicity [35].

### Hsp104 chimeras display reduced thermotolerance

We then characterized the thermotolerance activity of all chimeras (**Figure S14**) to understand how perturbing cognate interdomain interactions affects the disaggregase activity of each chimera. We found that, while swapping the NTD alone generally has a minimal effect on thermotolerance activity (the exception was Sr^NTD^Sc^NBD1:MD:NBD2:CTD^ which displayed minimal thermotolerance) swaps of subsequent domains, either alone or in combination, severely impair thermotolerance function (**Figure S14**). Some of the weak thermotolerance phenotypes may be attributable to low expression levels of certain chimeras from *pHSP104*; for instance, several chimeras between MbHsp104 and ScHsp104 are poorly expressed from *pHSP104* (**Figure S11C**). But, expression levels of chimeras alone are insufficient to explain their thermotolerance phenotypes: even specific chimeras that reduce TDP-43 toxicity and express well from *pHSP104* (e.g. Cr^NTD:NBD2:MD^Sc^NBD2:CTD^) fail to confer thermotolerance. We conclude that this observation provides further evidence that the mechanism that enables Hsp104 homologs to suppress TDP-43 toxicity is distinct from Hsp104 disaggregase activity.

### Select chimeras mimic potentiated ScHsp104 variants to suppress *α*Syn toxicity

Interestingly, two chimeras consisting of cognate NTD:NBD1 pairs fused to MD:NBD2:CTD from ScHsp104 unexpectedly suppressed *α*Syn toxicity. First, a chimera consisting of the CtHsp104 NTD and NBD1 fused to the ScHsp104 MD, NBD2, and CTD suppressed *α*Syn toxicity even more strongly than CtHsp104^WT^ (**Figure 7H**). Similarly, a chimera consisting of the PeHsp104 NTD and NBD1 fused to the ScHsp104 MD, NBD2, and CTD reduced *α*Syn toxicity (**Figure S12D**). This finding was particularly unexpected because full-length PeHsp104 is specific to TDP-43, and this same chimera is inactive against TDP-43 (**Figure 7F**). Both of these chimeras were toxic to yeast when expressed alone at 37°C (**Figure S13D, F**, boxed). Thus, in these cases, disruption of interaction between the NTD:NBD1 unit and the MD appears to mimic potentiated ScHsp104 variants and enables suppression of *α*Syn toxicity.

### The NBD2:CTD unit of TtHsp104 and TlHsp104 contribute to suppression of ***α*Syn toxicity**

For the *α*Syn-specific Hsp104 homologs, TtHsp104 and TlHsp104, chimeras consisting of NTD:NBD1:MD from TtHsp104 or TlHsp104 fused to NBD2:CTD from ScHsp104 were unable to reduce *α*Syn toxicity (**Figure 7D, I**). To test whether suppression of *α*Syn toxicity is encoded by residues within NBD2:CTD, we tested chimeras in which NTD:NBD1:MD were from ScHsp104 and NBD2:CTD was from either TtHsp104 or TlHsp104. However, these chimeras were also incapable of suppressing *α*Syn toxicity (**Figure 7J, K**; see **Figure S11J, K** for accompanying Western blots). Thus, these results suggest that additional residues or contacts in the NBD2:CTD unit are necessary but not sufficient for TtHsp104 and TlHsp104 to suppress *α*Syn toxicity.

### Homolog-mediated suppression of TDP-43 toxicity requires the NBD2:CTD unit

Finally, we tested whether NBD2 was required for suppression of TDP-43 toxicity by Hsp104 homologs, or whether expressing Hsp104 fragments encompassing the NTD, NBD1, and MD (which contain sequence determinants that reduce TDP-43 toxicity) would recapitulate the suppression of toxicity seen with the full-length homolog. We therefore co-expressed PeHsp104^1-541^and SrHsp104^1-551^ with TDP-43 in yeast. Neither of these fragments reduce TDP-43 toxicity (**Figure 7L**; see **Figure S11L** for accompanying Western blot). Two additional fragments, PeHsp104^767-914^ and SrHsp104^781-892^, which correspond to the small domain of NBD2 and the CTD also did not reduce TDP-43 toxicity (**Figure 7L**). Since these Hsp104 fragments are likely monomeric [73, 74], we thus conclude that proteotoxicity suppression is an emergent property of hexameric Hsp104 homologs or chimeras.

## DISCUSSION

Here, we used yeast toxicity models to identify naturally occurring Hsp104 homologs, from diverse eukaryotic hosts, capable of buffering proteotoxicity of several proteins implicated in human neurodegenerative diseases, including TDP-43 and αSyn (**Figures 1 and 2**). Excitingly, Hsp104 homologs that suppress TDP-43 toxicity in yeast also suppress TDP-43 aggregation in human cells (**Figure 3**), and Hsp104 homologs that suppress *α*Syn toxicity in yeast also suppress *α*Syn toxicity-induced neurodegeneration in *C. elegans* (**Figure 4**). Thus, we suggest that, like previously-defined potentiated Hsp104 variants, these naturally-occurring Hsp104 variants may be able to mitigate proteotoxicity in a wide variety of circumstances, including in metazoan systems.

Several features of the Hsp104 homologs presented here contrast with potentiated Hsp104 variants. Hsp104 homologs are substrate-specific and typically only suppress either TDP-43 or *α*Syn toxicity (the exception is CtHsp104). Indeed, we did not isolate any Hsp104 homologs capable of suppressing FUS toxicity. In contrast, potentiated Hsp104 variants typically are able to suppress toxicity of multiple toxic substrates (**Figures 1, 2, and S3**) [30, 32–34, 37]. Natural Hsp104 homologs also displayed no intrinsic toxicity, even when expressed at elevated temperatures (**Figure S2**). Thus, extant Hsp104 homologs have likely been filtered through natural selection to avoid deleterious and destabilizing sequences. In contrast, potentiated Hsp104 variants have predominantly been engineered by destabilizing the NBD1:MD interface [30, 33-35, 37]. This difference is further reflected in fundamental mechanistic differences between how the Hsp104 homologs described here and potentiated Hsp104 variants operate to antagonize proteotoxic misfolding. Engineered Hsp104 variants are generally enhanced disaggregases [30, 32–34]. We hypothesize that these enhanced disaggregase and unfoldase activities may come at the cost of substrate specificity (e.g. by mistargeting natively-folded complexes for disassembly), analogous to trade-offs between speed and fidelity observed in other NTPase molecular machines, such as RNA polymerases [75]. However, the Hsp104 homologs presented here are not similarly enhanced disaggregases, and in fact do not require disaggregase activity to antagonize proteotoxic misfolding of TDP-43 or *α*Syn. Rather, they appear to act as ATP-independent chaperones to suppress toxicity of specific substrates (**Figures 5 and 6**).

The dual role of Hsp104 as a molecular chaperone capable of preventing substrate misfolding in addition to a disaggregase capable of reversing protein aggregation has been long-appreciated [19, 20]. Both of these activities require ATPase activity for maximum effect, although disaggregase activity is much more sensitive to reduced ATPase activity than chaperone activity [19, 20]. Here, we find that diverse Hsp104 homologs are able to suppress toxicity of select substrates in a manner that requires neither active ATPase domains nor substrate engagement by the canonical pore loops (**Figure 6**). Rather, we have uncovered genetic variation outside of these core AAA+ features that enables molecular recognition of specific substrates. Thus, in future work it will be important to tune this genetic variation to create designer disaggregases with highly specific molecular recognition. Indeed, we envision making modular disaggregases by combining potentiating mutations with molecular recognition motifs to generate Hsp104 variants with enhanced specificity *and* disaggregase activity.

What sequence determinants of Hsp104 homologs enable their toxicity suppression phenotypes? Hsp104 homologs that suppress TDP-43 or suppress *α*Syn are more similar to each other than between groups (average sequence identities of 56%, 76%, and 44%, respectively; **Table S3** and **Figure S6B**). To further address this question, we first tested the effect of deleting the NTD on the toxicity suppression phenotypes associated with Hsp104 homologs (**Figure 6**).

We observed that suppression of TDP-43 toxicity by MbHsp104 and CrHsp104, and suppression of *α*Syn toxicity by TtHsp104, depends on the presence of the NTD of these homologs (**Figure 6**). The ScHsp104 NTD enables many aspects of ScHsp104 function, including hexamer cooperativity, substrate binding, amyloid dissolution, and proteotoxicity suppression by potentiated Hsp104 variants [35, 72]. We suggest that the NTD-dependent rescue phenotypes we observed may reflect a role of NTD-substrate binding in suppressing proteotoxicity.

While the NTD is necessary for proteotoxicity suppression under some circumstances, it is not universally required. Three out of five TDP-43-specific homologs did not require the NTD to suppress TDP-43 toxicity, and TlHsp104 did not require the NTD to mitigate *α*Syn toxicity (**Figure 6**). Furthermore, simply replacing the ScHsp104 NTD with the NTD from MbHsp104 or CrHsp104 does not enable suppression of TDP-43 toxicity, nor does replacing the ScHsp104 NTD with the NTD from TtHsp104 impart suppression of *α*Syn toxicity (**Figure 7**). Thus, we reasoned that additional domains must be required. Indeed, for TDP-43, we found that replacing the NTD:NBD1:MD unit of the ScHsp104 hexamer with the homologous domains from any TDP-43-rescuing homolog enables suppression of TDP-43 toxicity by the resulting chimera (**Figure 7**). The NBD2 and CTD domains were required to facilitate suppression of toxicity, as fragments of PeHsp104 and SrHsp104 encompassing NTD:NBD1:MD but lacking NBD2 and the CTD failed to suppress toxicity. However, NBD2:CTD could be from ScHsp104. In contrast, NBD2:CTD could not be from ScHsp104 for suppression of *α*Syn toxicity by TtHsp104 and TlHsp104 (**Figure 7**). We propose two explanations for these divergent observations. The first is that, particularly for the homologs where NTD deletion does not diminish suppression of toxicity (e.g. GsHsp104, PeHsp104, and SrHsp104 for TDP-43 and TlHsp104 for *α*Syn), there are important sequence determinants throughout the protein that enable each homolog to produce a protective phenotype. The second possible explanation is that these other domains contribute to suppression of toxicity indirectly by stabilizing overall protein architecture of the chimeras. There are no obvious trends in the primary sequence of the homologs, however, that would clarify these two competing possibilities. Indeed, further work is needed to gain a complete understanding of how homologous Hsp104 sequences confer variable substrate-specific proteotoxicity suppression.

That nearly half of the Hsp104 homologs we tested (8/17) were able to antagonize toxicity of either TDP-43 or *α*Syn (or both, in the case of CtHsp104) was unanticipated. There are no clear TDP-43 or *α*Syn homologs in the host species from which we selected Hsp104 homologs to test. Our observation that, for a given Hsp104, there was not a predictable relationship between TDP-43 or *α*Syn toxicity suppression and thermotolerance function, which may more closely reflect the primary *in vivo* function of these Hsp104 homologs, suggests one interpretation involving cryptic genetic variation [76]. Variation that is neutral or nearly neutral with respect to thermotolerance function, but which enables suppression of TDP-43 or *α*Syn, may have accumulated in these lineages in the absence of selective pressure and was only revealed by our synthetic experimental paradigm. Alternatively, it is possible that the Hsp104 homologs we tested are adapted to host proteomes that have higher-than-average content of TDP-43-like or *α*Syn-like motifs, which is reflected in their ability to selectively antagonize misfolding of similar motifs in TDP-43 or *α*Syn. Nevertheless, the fact that Hsp104 homologs that are active against at least one of TDP-43 or *α*Syn were so widespread amongst diverse eukaryotic lineages, and the fact that we have now identified ∼100 potentiating mutations in Hsp104 that enable similar toxicity suppression phenotypes (albeit by different mechanisms) suggests that Hsp104, and possibly other AAA+ proteins, are uniquely poised to buffer deleterious protein misfolding and aggregation [15].

## ACKNOWLEDGEMENTS

We thank Ryan Cupo, JiaBei Lin, and Charlotte Fare for comments on the manuscript. We thank Susan Lindquist, Aaron Gitler, Simon Alberti, and Adrian Tsang for generous provision of reagents. This work was supported by a Muscular Dystrophy Association Research Award (J.S.), an ALS Association Award (J.S.), the Life Extension Foundation (J.S.), a Linda Montague Pechenik Research Award (J.S.), the Packard Center for ALS Research at Johns Hopkins University (J.S.), Target ALS (M.E.J. and J.S.), NSF Graduate Research Fellowship DGE-0822 (L.M.C.), a Blavatnik Family Fellowship (E.C.), NIH grants T32GM071399, F31NS101807 (to Z.M.M.), T32GM008076 (E.C.), and R01GM099836 (J.S.).

## AUTHOR CONTRIBUTIONS

ZMM conceived, designed, and performed experiments and analyzed data. KS designed and performed experiments in human cells. HK and ZY designed and performed experiments with *C. elegans*. LMC designed and performed SEVI remodeling experiments. MEJ designed and performed experiments with ClpB variants. EC, EG, KM, RJ, and AJ contributed reagents. KAC, GAC, OS, and JS conceived and designed experiments, analyzed data, and supported research. ZMM and JS wrote the paper with feedback from all authors.

## Methods

### Bioinformatic analyses

Multiple sequence alignments of Hsp104 homologs were generated with Clustal Omega [77]. Multiple sequence alignments were visualized in JalView [78]. The phylogenetic tree in **Figure 1** was generated using EMBL-EBI Simple Phylogeny tool [79]. Pairwise sequence identities in **Table S3** and **Figure S6B** were calculated using UniProt Align tool. Divergence times shown in **Figure 5** were from TimeTree [80].

### Yeast strains and media

All yeast were WT W303a (*MATa, can1-100, his3-11,15, leu2-3,112, trp1-1, ura3-1, ade2-1*) or the isogenic strain W303a*Δhsp104* [81]. Yeast were grown in rich medium (YPD) or in the appropriate synthetic selection media. Media was supplemented with 2% glucose, raffinose, or galactose.

### Plasmids

Yeast expression vectors encoding TDP-43 (pAG303GAL-TDP-43), FUS (pAG303GAL-FUS), and αSyn-GFP (pAG303GAL-αSyn-GFP and pAG304GAL-αSyn-GFP) were from Aaron Gitler [13, 40, 47]. pRS313HSE-ScHsp104^WT^-FLAG, pAG416GAL-ScHsp104^WT^-FLAG, pAG416GAL-ScHsp104^A503S^-FLAG, and pAG416GAL-CtHsp104-FLAG have been described previously [27]. cDNAs encoding some Hsp104 homologs were kind gifts from Adrian Tsang (Concordia University; TtHsp104, TlHsp104, ChtHsp104, LtHsp104, MtHsp104, StHsp104, and TaHsp104), Susan Lindquist (Whitehead Institute; AtHsp104), and Simon Alberti (MPI-CBG; DdHsp104), while yeast codon-optimized cDNAs encoding other Hsp104s were synthesized by Invitrogen (MbHsp104 and PfHsp104) or Genscript (GsHsp104, CrHsp104, SrHsp104, and PeHsp104). Hsp104 cDNAs codon-optimized for expression in mammalian cells were used for experiments shown in **Figure 4**. Amino acid sequences of all Hsp104 homologs are included in **Table S1**. Gateway BP reactions were used to shuttle Hsp104 genes into a Gateway entry vector, pDONR221-ccdB. The entry clones were then used to shuttle Hsp104 ORFs into suitable yeast (pAG416GAL-ccdB or pAG413GAL-ccdB or pRS313HSE-ccdB), *C. elegans* (pDAT-ccdB), or mammalian (pInducer20-ccdB) expression vectors via LR reactions. MbHsp104 and CrHsp104 were cloned into pNOTAG for bacterial expression through NdeI and SacI sites. Hsp104 mutants were obtained by Quikchange mutagenesis. Hsp104 chimeras were cloned by overlap-extension PCR. Sequences of all Hsp104 chimeras are included in **Table S2**.

Doxycycline-inducible TDP-43*Δ*NLS mammalian expression vector [57] was modified as follows. Human wild-type TDP-43 was amplified in two separate PCR reactions excluding the NLS and reassembled using Gibson cloning (NEB) into a Doxycycline-inducible expression vector containing an N-terminal mClover3 tag. PCR primers used to generate TRE-mClover3-linker-TDP-43deltaNLS: 5’-GGATCCGGAAGTGGCTCAAGCGGAATGTCTGAATATATTCGGGT-3’; 5’-TGCTGCTGCCACTGCCACTGCTGATGAAGCATCTGTCTCATCCATTGCTGCTGCGTTAT CTTTTGGATAGTTGACA-3’; 5’-ATCAGCAGTGGCAGTGGCAGCAGCAGCAGTCCAGAAAACATCCGA-3’; 5’-AAGTTTGTTGCGCCGGATCC CATTCCCCAGCCAGAAGACT-3’.

### Yeast transformation and spotting assays

Yeast were transformed according to standard protocols using polyethylene glycol and lithium acetate [82]. For spotting assays, yeast were grown to saturation overnight in synthetic raffinose dropout media at 30 °C. Cultures were normalized and serially diluted 5-fold and spotted in duplicate onto synthetic dropout media containing glucose or galactose. Plates were analyzed after growth for 2-3 days at 30 °C.

### Assessing toxicity of Hsp104 homologs and chimeras

W303a*Δhsp104* yeast were transformed with the indicated Hsp104 plasmid. Transformants were grown to saturation overnight in synthetic raffinose media. Cultures were normalized, spotted onto synthetic dropout media containing glucose or galactose, and incubated at 30 °C or 37 °C for 2-3 d before growth was documented.

### Western blotting

Yeast were grown in galactose-containing media to induce protein expression for 5h (for strains expressing Hsp104s alone or with TDP-43) or 8h (for strains expressing αSyn). Cultures were normalized to OD_600_=0.6, and 6 mL of cells were harvested. For heat shock controls, samples were incubated at 37 °C for 30 minutes prior to processing. Yeast lysates were extracted by incubation with 0.1M NaOH at room temperature for 5 min. Lysates were mixed with SDS sample buffer, boiled for 5 min, and subjected to Tris-HCl SDS-PAGE (4-20% gradient, Bio-Rad) followed by transfer to a PVDF membrane (Millipore). Membranes were blocked in Odyssey blocking buffer (LI-COR) for 1h at room temperature or overnight at 4 °C. Primary antibody incubations were performed at 4 °C overnight or at room temperature for 2h. After washing with PBST, membranes were incubated with fluorescently labeled secondary antibodies at room temperature for 1h, followed by washing with PBST. Proteins were detected using an Odyssey Fc Dual-Mode Imaging system. Primary antibodies used: mouse monoclonal anti-FLAG M2 (Sigma-Aldrich), rabbit polyclonal anti-TDP-43 (Proteintech), rabbit polyclonal anti-GFP (Sigma-Aldrich), mouse monoclonal 3-phosphoglycerate kinase (Novex), rabbit polyclonal anti-Hsp26 (Johannes Buchner, TU-Munich), fluorescently labeled anti-mouse and anti-rabbit secondary antibodies (Li-Cor).

### Fluorescence microscopy

To visualize changes in TDP-43 and *α*Syn localization in response to coexpression of various Hsp104s, yeast strains expressing fluorescently-tagged TDP-43 and *α*Syn were previously generated and described [30]. For microscopy, these strains were grown for 5 h (TDP-43) or 8 h (*α*Syn) in galactose-containing media at 30 °C. For TDP-43 microscopy, cells were harvested and fixed with ice-cold 70% ethanol, washed 3 times with ice-cold PBS, and stained with 4’,6-diamidino-2-phenylindole (DAPI) in Vectashield mounting medium (Vector Laboratories) to visualize nuclei. For *α*Syn, imaging was performed with live cells. Images were collected at 100x magnification using a Leica-DMIRBE microscope and processed using ImageJ software (NIH).

### HEK-293T cell culture and transfections

HEK-293T cells were maintained in Dulbecco’s modified Eagle’s medium, high glucose (Gibco) containing 10% fetal bovine serum (Life Sciences), 1% non-essential amino acids (Gibco), and 1% anti-anti (Gibco). Cells were plated in gelatin-coated 6-well plates at a density of 3×10^6 cells/plate 24hr before transfection. Cells were transfected with 2 µg total DNA and 7.35 µl polyethylenimine HCl MAX transfection reagent (Polysciences, Inc). Wells co-transfected with mClover3-TDP*Δ*NLS and HSP104 variants received 1 µg of each plasmid. Media was changed 6hrs post-transfection to media containing 1 µg/ml of doxycycline hyclate (Sigma-Aldrich) in order to induce transgene expression. Transfected cells were lifted every 24 h over 2 days, at which point cells were analyzed by FACS (FACSAria Fusion BD). Cells were gated to have a narrow range of FCS and SSC values and to be fluorescence positive. TDP-43 aggregation was quantified by comparing the height (FITC-H) to the width (FIRC-W) of the fluorescence channel using 488nm laser and FITC filters.

### Generation of transgenic *C. elegans* and neurodegeneration analysis

Nematodes were maintained through well-established methods [83]. Constructs were injected into animals to create transgenic line using previously described methods [84]. Strains UA381 (*baln11* [P*_dat-1_*::*α*-syn, P*_dat-1_*::GFP]; *baEx210* [P*_dat-1_*::*CtHsp104*, *rol-6*], UA382 (*baln11* [P*_dat-1_*::*α*-syn, P*_dat-1_*::GFP]; *baEx211* [P*_dat-1_*::*TtHsp104*, *rol-6*], UA383 (*baln11* [P*_dat-1_*::*α*-syn, P*_dat-1_*::GFP]; *baEx212* [P*_dat-1_*::*TIHsp104*, *rol-6*] were generated by injecting 50 ng/μl of corresponding plasmid construct into UA44 (*baln11* [P*_dat-1_*::*α*Syn, P*_dat-1_*::GFP]) with phenotypic marker (*rol-6*, 50 ng/μl, for roller expression). Three independent stable lines were created for each group. For dopaminergic neurodegeneration analyses, the transgenic animals were scored as described previously [85]. Briefly, on the day of analysis, the six anterior dopaminergic neurons [four CEP (cephalic) and two ADE (anterior deirid)] were examined in 30 animals randomly selected for each trial worms that express the roller marker in the body wall muscle cells. Neurons were analyzed for any degenerative phenotypes, such as a missing dendritic process, cell body loss, or a blebbing neuronal process. Each animal was scored as having normal or wild-type neurons when none of the degenerative phenotypes were present in any anterior dopaminergic neurons. Three independent transgenic worm lines were analyzed per genetic background and an average of total percentage of worms with normal neurons was reported in the study. One-way ANOVA, followed by a Dunnett’s multiple comparisons *post hoc* test, was performed for statistical analysis using GraphPad Prism Software.

### Quantitative real-time PCR

RNA isolation and RT-qPCR was performed on worms using previously published methods [86]. Briefly, total RNA was isolated from 100 young adult (day 4 post hatching) nematodes from corresponding transgenic lines using TRI reagent (Molecular Research Center). The genomic DNA contamination was removed with 1 µl of DNaseI (Promega) treatment for 60 minutes at 37°C, then with DNase Stop solution for 10 minutes at 65°C. 1µg of RNA was used for cDNA synthesis using the iScript Reverse Transcription Supermix for RT-qPCR (Bio-Rad) following the manufacturer’s protocol. RT-qPCR was performed using IQ-SYBR Green Supermix (Bio-Rad) with the Bio-Rad CFX96 Real-Time System. Each reaction contained 7.5 μl of the IQ SYBR Green Supermix, 200 nM of forward and reverse primers and 5 ng of cDNA, to a final volume of 15 μl. The cycling conditions were as follows: polymerase activation and DNA denaturation at 95°C for 3 min, followed by 35 cycles of 10 s at 95°C, 30 s at 60°C. After the final cycle, a melting curve analysis was performed using the default setting of CFX96 Real-Time System. A single melt peak for each targeted gene was observed and no non-specific amplification was detected in each reaction mixture by agarose gel electrophoresis. PCR efficiency was calculated from standard curves that were generated using serial dilution (E*_α-syn_*=98.8%, E*_TtHsp104_*=101.0%, E*_TlHsp104_*=97.4%, E*_tba-1_*=98.8%, E*_cdc-42_*=98.4%, E*_snb-1_*=95.3%). The expression levels of *α*-syn and Hsp104 variants were normalized to three reference genes (*snb-1, cdc-42*, and *tba-1*). No amplification was detected in NTC and NRT controls. The reference target stability was analyzed by GeNorm and passed for all reference genes listed above. All target genes were measured in triplicates for three independent transgenic lines for each sample in this study. The data analysis was performed by the Gene Expression Module of CFX Manager software.

The following primers were used for the assays;

*CtHsp104* Forward: GACGAAGCGTGTGCCAATAC

*CtHsp104* Reverse: CACTTCCTGGAGCCGCTG

*TtHsp104* Forward: CAACTACTTCCTGCCCGAG

*TtHsp104* Reverse: ATCTGGACGTTGCGGTCGT

*TlHsp104* Forward: AACCGTCTCACCAAGCGTG

*TlHsp104* Reverse: GCCTCTCCGAGATAGTCCT

*α-syn* Forward: ATGTAGGCTCCAAAACCAAGG

*α-syn* Reverse: ACTGCTCCTCCAACATTTGTC

*snb-1* Forward: CCGGATAAGACCATCTTGACG

*snb-1* Reverse: GACGACTTCATCAACCTGAGC

*cdc-42* Forward: CCGAGAAAAATGGGTGCCTG

*cdc-42* Reverse: TTCTCGAGCATTCCTGGATCAT

*tba-1* Forward: ATCTCTGCTGACAAGGCTTAC

*tba-1* Reverse: GTACAAGAGGCAAACAGCCAT

### Thermotolerance

W303a*Δhsp104* yeast were transformed with plasmids bearing either Hsp104 from *S. cerevisiae,* the indicated Hsp104 homolog, or Hsp104 mutant under the native *HSP104* promoter (except for TtHsp104 and derivative mutants, which were expressed from p*GAL*), or an empty vector control. Transformants were selected, grown to saturation in yeast minimal media (SD-His), and then diluted to OD_600_=0.2 in fresh SD-His. Yeast were allowed to double at 30 °C, after which cultures were normalized. Cells were then heat shocked at 50 °C for the indicated time and cooled for 2 min on ice. Cultures were then diluted 1000-fold, plated on SD-His, and plates were incubated at 30 °C for 2-3 days to observe viable colonies.

### Protein expression and purification

Untagged ScHsp104 was expressed from the pNOTAG-ScHsp104 vector [73] and purified as previously described [60, 87]. Briefly, pNOTAG-ScHsp104 was used to transform BL21(DE3)RIL *E. coli*. Transformed cells were grown in 2xYT broth supplemented with 25 µg/ml chloramphenicol and 100 µg/ml ampicillin at 37 °C until an OD_600_ of 0.4-0.6 was reached, at which point cells were cooled to 15 °C. Expression was induced by addition of 1 mM isopropyl 1-thio-β-D-galactopyranoside for 15-18 h. Cells were harvested by centrifugation (4,000×g, 4°C, 25 min), resuspended in lysis buffer (50 mM Tris-HCl, pH 8.0, 10 mM MgCl_2_, 2.5% glycerol, 2 mM β-mercaptoethanol, 5 µM pepstatin, c0mplete EDTA-free protease inhibitors (Roche). Cells were treated on ice with 20 mg lysozyme per 1L culture and lysed by sonication. Cell debris was removed by centrifugation at 16,000×g at 4°C for 20 min, and the supernatant was applied to Affi-Gel Blue resin (Bio-Rad). Resin was incubated with the lysates for 4 h at 4 °C with slow rotation. Resin was then washed 4 times with wash buffer (50 mM Tris-HCl, pH 8.0, 10 mM MgCl_2_, 100 mM KCl, 2.5% glycerol, 2 mM β-mercaptoethanol).

ScHsp104 was eluted with wash buffer supplemented with 1 M KCl. The protein was then exchanged into running buffer Q (20 mM Tris-HCl pH 8.0, 0.5 mM EDTA, 5 mM MgCl_2_, 50 mM NaCl), further purified by ResourceQ anion exchange chromatography, and eluted with a linear salt gradient (50 mM-1 M NaCl). Eluted protein was then exchanged into storage buffer (40 mM HEPES-KOH pH 7.4, 150 mM KCl, 20 mM MgCl_2_, 10% glycerol, 1 mM DTT), snap-frozen, and stored at −80 °C until use.

MbHsp104 and CrHsp104 were expressed as untagged proteins by subcloning the MbHsp104 and CrHsp104 ORFs into pNOTAG through NdeI and SacI sites. Protein expression and lysis were carried out as for ScHsp104. However, MbHsp104 and CrHsp104 bound poorly to Affi-Blue resin, even after extended incubation times (data not shown). Instead, MbHsp104 and CrHsp104 were precipitated from clarified bacterial lysates by addition of solid ammonium sulfate to 40% of saturation. Precipitates were collected by centrifugation (16,000×g at 4 °C, 20 min) and resuspended in buffer containing 50 mM Tris-HCl, pH 8.0, 10 mM MgCl_2_, 100 mM KCl, 2.5% glycerol, 2 mM β-mercaptoethanol. To resolubilize precipitates, ammonium sulfate was removed by dialysis against this buffer, with 3 buffer changes. The dialyzate was filtered and applied to a 5 mL HiTrapQ column and purified with running buffer Q (20 mM Tris pH 8.0, 0.5 mM EDTA, 5 mM MgCl_2_, 50 mM NaCl) and eluted with a linear gradient of buffer Q+ (20 mM Tris pH 8.0, 0.5 mM EDTA, 5mM MgCl_2_, 1 M NaCl) over 40 column volumes. Peak fractions were collected, pooled, and exchanged into size exclusion buffer (40 mM HEPES-KOH pH 7.4, 150 mM KCl, 10 mM MgCl_2_ 1 mM DTT) and further purified by size exclusion chromatography on a pre-equilibrated Superdex 200 column (GE healthcare). Protein in size exclusion buffer was concentrated to ∼10 mg/mL, supplemented with 10% glycerol, snap-frozen and stored at −80 °C until use.

CtHsp104, TtHsp104, and TlHsp104 were expressed from pMCSG68 vector as TEV protease-cleavable His_6_-tagged fusion proteins. Protein expression and lysis were carried out as for other Hsp104s, except in this case lysis buffer consisted of 40 mM HEPES-KOH pH 7.4, 500 mM KCl, 20 mM MgCl_2_, 2.5% glycerol, 20 mM imidazole, 2 mM β-mercaptoethanol supplemented with 5 µM pepstatin A and complete protease inhibitor tablets. Clarified lysate was loaded onto Ni-NTA resin. The resin was washed with 10 volumes of wash buffer (same formulation as lysis buffer except without protease inhibitors) and eluted in wash buffer supplemented with 350 mM imidazole. TEV protease was added to the eluted protein, and the sample was dialyzed against wash buffer containing no imidazole for 4 h at room temperature followed by ∼16h at 4 °C. After dialysis and cleavage, the protein was loaded onto a second Ni-NTA column to remove the His_6_ tag and uncleaved protein. Eluted protein was pooled, concentrated, and exchanged into high salt storage buffer (40 mM HEPES-KOH pH 7.4, 500 mM KCl, 20 mM MgCl_2_, 10% glycerol, and 1 mM DTT), snap-frozen in liquid nitrogen, and stored at −80 °C until further use.

Ssa1, Hsc70, Sis1, Ydj1, Hdj1, and Hdj2 (in pESUMO (Life Sensors)) were expressed as N-terminally His_6_-SUMO-tagged proteins in BL21(DE3)RIL cells. Transformed cells were grown at 37 °C in Luria broth supplemented with 25 µg/ml chloramphenicol and 100 µg/ml ampicillin to an OD_600_ ∼0.5. Cultures were cooled to 15 °C, and expression was induced with 1 mM IPTG for 16 h. Cells were harvested, resuspended in lysis buffer (50 mM HEPES pH 7.5, 750 mM KCl, 5 mM MgCl_2_, 10% glycerol, 20 mM imidazole, 2 mM β-mercaptoethanol, 5 µM pepstatin A, and c0mplete protease inhibitor (Roche)), and lysed by treatment with lysozyme and sonication. Lysates were clarified by centrifugation (16,000×g, 20 min, 4°C), and incubated with Ni-NTA resin for 90 min at 4°C. Resin was washed with 10 column volumes of wash buffer (50 mM HEPES pH 7.5, 750 mM KCl, 10 mM MgCl_2_, 10% glycerol, 20 mM imidazole, 1 mM ATP, 2 mM β-mercaptoethanol) and eluted with 2 column volumes of elution buffer (wash buffer+300 mM imidazole). To cleave the His_6_-SUMO tag, Ulp1 was added at a 1:100 molar ratio, and imidazole was removed by dialysis against wash buffer. After dialysis, protein was loaded onto a 5 mL HisTrap column (GE Healthcare) and eluted with a linear imidazole gradient (20-350 mM) over 40 column volumes. Fractions containing cleaved protein were pooled, concentrated, and purified further by Resource Q (Ssa1, Hsc70, Ydj1, and Hdj2) or Resource S (Sis1 and Hdj1) ion exchange chromatography.

TDP-43 was expressed with an N-terminal, TEV-cleavable maltose binding protein (MBP) tag in BL21(DE3)RIL cells. Transformed cells were grown at 37 °C in Luria broth supplemented with 25 µg/ml chloramphenicol and 100 µg/ml ampicillin to an OD_600_ ∼0.8. Cultures were cooled to 15 °C, and expression was induced with 1 mM IPTG for 16 h. Cells were harvested, resuspended in lysis buffer (50 mM HEPES-KOH pH 7.5, 500 mM NaCl, 10% glycerol, 2 mM EDTA, 2 mM DTT, and c0mplete protease inhibitor (Roche)), and lysed by treatment with lysozyme and sonication. Lysates were clarified by centrifugation (16,000×g, 20 min, 4 °C). Cleared lysate was added to a 50% slurry of amylose resin (New England Biolabs) in lysis buffer and incubated at 4 °C for 30 min to bind MBP-TDP-43. The resin was washed with lysis buffer, and eluted MBP-TEV with elution buffer (50 mM HEPES-KOH pH 7.5, 500 mM NaCl, 10% glycerol, 2 mM EDTA, 2 mM DTT, 10 mM maltose). Purified fractions were pooled, snap-frozen, and stored at −80 °C until use.

### ATPase activity

Hsp104 (0.25 µM monomer) was incubated with ATP (1 mM) for 5 min at the indicated temperatures in buffer (25 mM HEPES-KOH pH 7.4, 150 mM potassium acetate, 10 mM magnesium acetate, 10 mM DTT). ATPase activity was determined by inorganic phosphate release using a malachite green phosphate detection kit (Innova). Background hydrolysis was determined at time zero and subtracted.

### Luciferase reactivation

Aggregated luciferase (50 nM) was incubated with Hsp104 (0.167 µM hexamer) with ATP (5 mM) and an ATP regeneration system (ARS; 1 mM creatine phosphate, 0.25 µM creatine kinase) plus or minus 0.167 µM Hsp70 (Ssa1 or Hsc70) and 0.167 µM Hsp40 (Sis1, Ydj1, Hdj1, or Hdj2, as indicated). Luciferase activity was assessed by luminescence on a TECAN Safire II plate reader.

### Semen-Derived Enhancer of Virus Infection (SEVI) remodeling

SEVI remodeling was performed as previously described [53]. SEVI fibrils (20 µM monomer) were incubated with Hsp104 homologs (3 µM hexamer) in LRB buffer in the presence of ATP (5 mM) and an ATP regeneration system (0.1 mM ATP, 0.02 mg/mL creatine kinase, 10 mM creatine phosphate). Samples were incubated at 37 °C for the duration of the experiments. At various time points, aliquots were removed, added to a 96-well plate containing a solution of 25 µM ThT in LRB buffer. ThT fluorescence characteristics were measured on a Tecan Safire^2^ microplate reader with excitation and emission filters set to 440 nm and 482 nm, respectively. To assess fibril morphology by negative stain EM, reaction aliquots were spotted on Formvar carbon-coated grids (EM Sciences) and stained with 2% uranyl acetate. Samples were visualized using a JEOL-1010 electron microscope.

### TDP-43 aggregation experiments

TDP-43 aggregation was initiated by incubating 3 µM MBP-TEV-TDP-43 with TEV protease in assembly buffer (40 mM HEPES-KOH pH 7.5, 150 mM NaCl, 10 mM MgCl_2_, 1 mM DTT) without agitation. Aggregation was monitored by turbidity (A_395 nm_) in a TECAN Infinite M1000 plate reader. In some reactions, either ScHsp104, MbHsp104, or CrHsp104 were added (6 µM) with or without ATP (5 µM) and regeneration system (1 mM creatine phosphate and 0.25 µM creatine kinase). We verified that Hsp104s nor ATP affected cleavage of the MBP tag by Western blotting (see **Figure S2**).

## Supplemental Information

**Figure S1.**
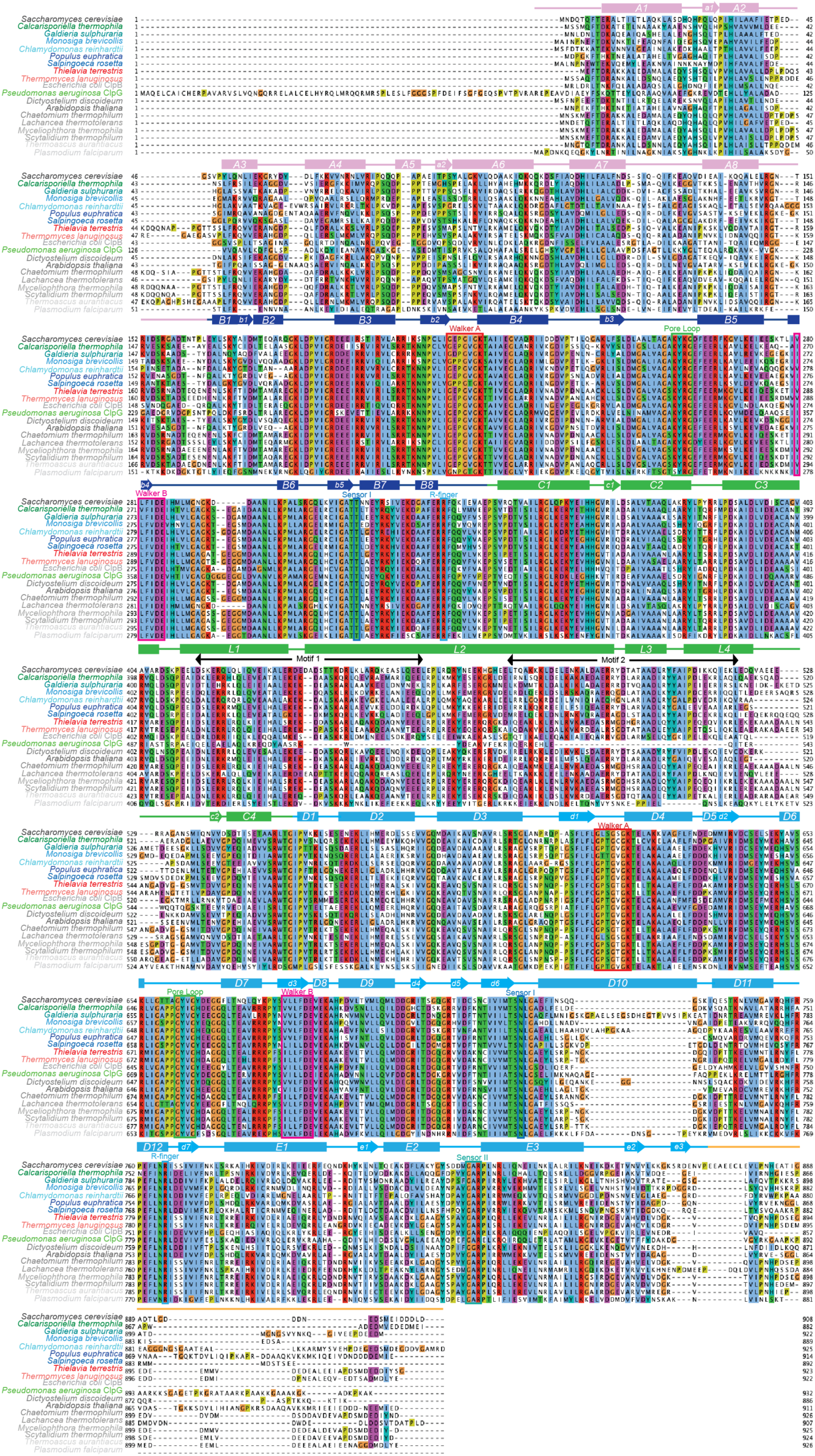
Alignment of all Hsp104 homologs investigated in this study (related to Figures 1 and 2). Amino acid sequences of Hsp104 homologs were aligned using Clustal Omega, and the multiple sequence alignment was visualized with JalView. Positions in the alignment are colored by Clustal X convention: conserved hydrophobic positions (A,I,L,M,F,W,V,C) are blue; conserved basic residues (K,R) are red; acidic residues (E,D) are magenta; polar residues (N,Q,S,T) are green; aromatic residues (H,Y) are cyan; conserved cysteines are pink; glycines are orange, prolines are yellow. Structural elements of Hsp104 are indicated. *C. thermophila* and ClpG are in green, TDP-43-specific homologs are colored in shades of blue, *α*Syn-specific homologs are colored in red, and non-rescuing homologs are colored in shades of gray.

**Figure S2.**
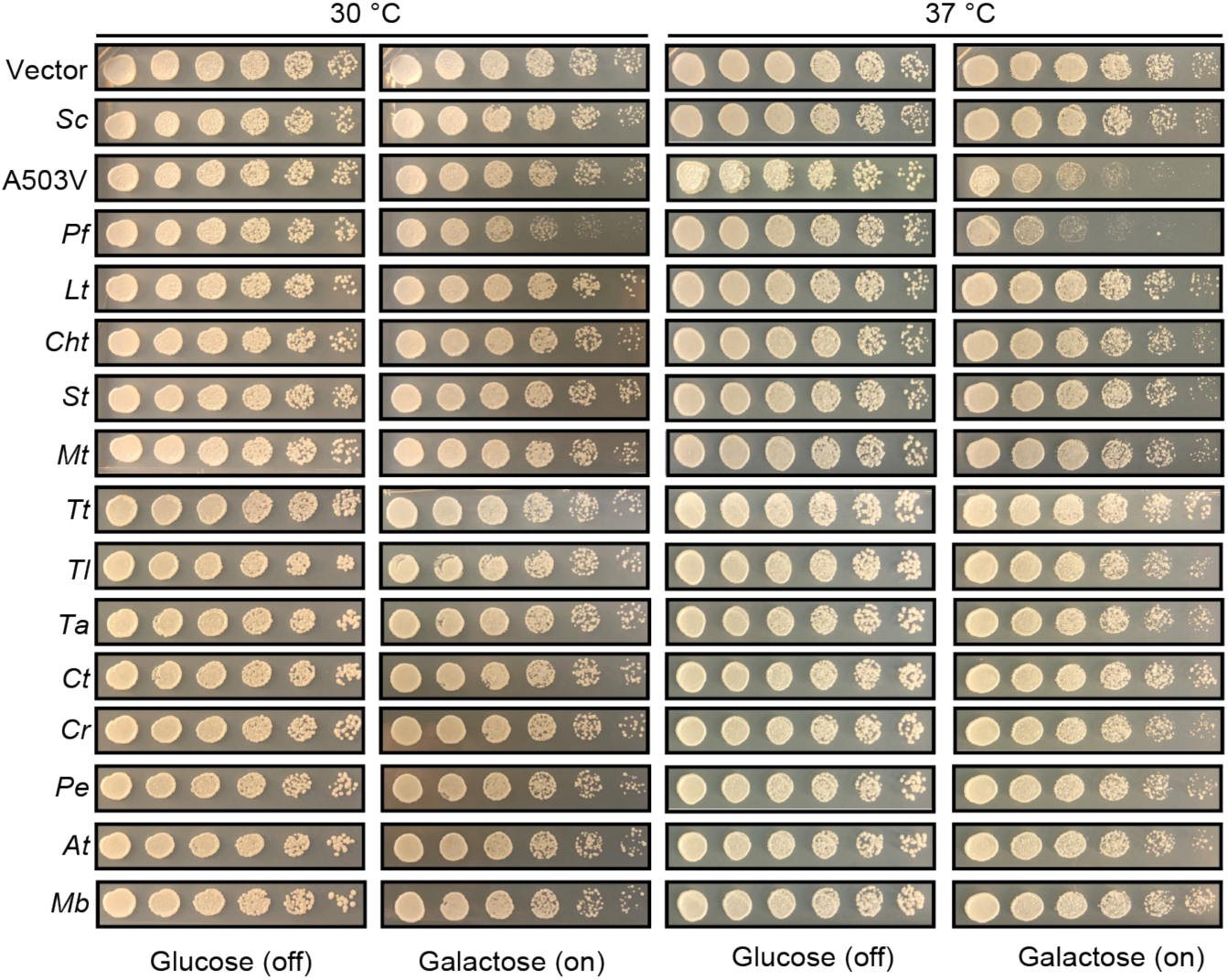
Hsp104 homologs do not typically induce temperature-dependent toxicity (related to Figures 1 and 2). Spotting assay showing that yeast strains expressing the indicated Hsp104 homolog do not show a temperature-dependent growth defect. None of the Hsp104 homologs were toxic at 30°C or 37°C with the exception of PfHsp104, which was toxic at both temperatures. PfHsp104 was even more toxic than the potentiated Hsp104 variant, Hsp104^A503V^, at 37°C. Unlike PfHsp104, Hsp104^A503V^ was not toxic at 30°C.

**Figure S3.**
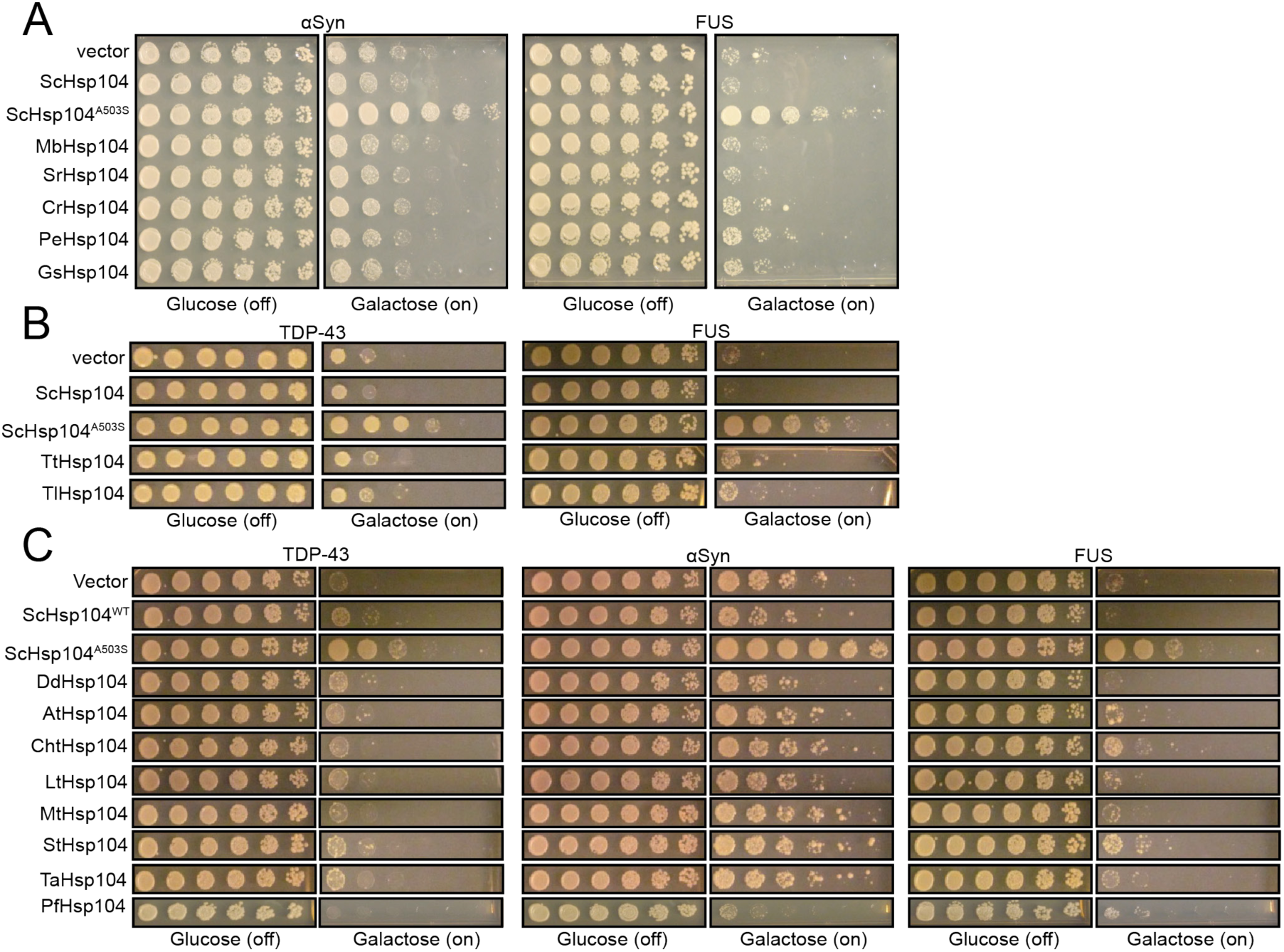
Suppression of TDP-43 or *α*Syn toxicity by select Hsp104 homologs is a substrate-specific effect (related to Figures 1 and 2) (A) Spotting assay showing that MbHsp104, SrHsp104, CrHsp104, PeHsp104, and GsHsp104, which all suppress TDP-43 toxicity, do not suppress *α*Syn nor FUS toxicity. ScHsp104^A503S^ is included as a positive control and empty vector and ScHsp104^WT^ are included as negative controls (B) Spotting assay showing that TtHsp104 and TlHsp104 do not suppress TDP-43 nor FUS toxicity (C) Spotting assay showing that other Hsp104s investigated in this study do not suppress toxicity of TDP-43 (left), *α*Syn (middle), or FUS (right).

**Figure S4.**
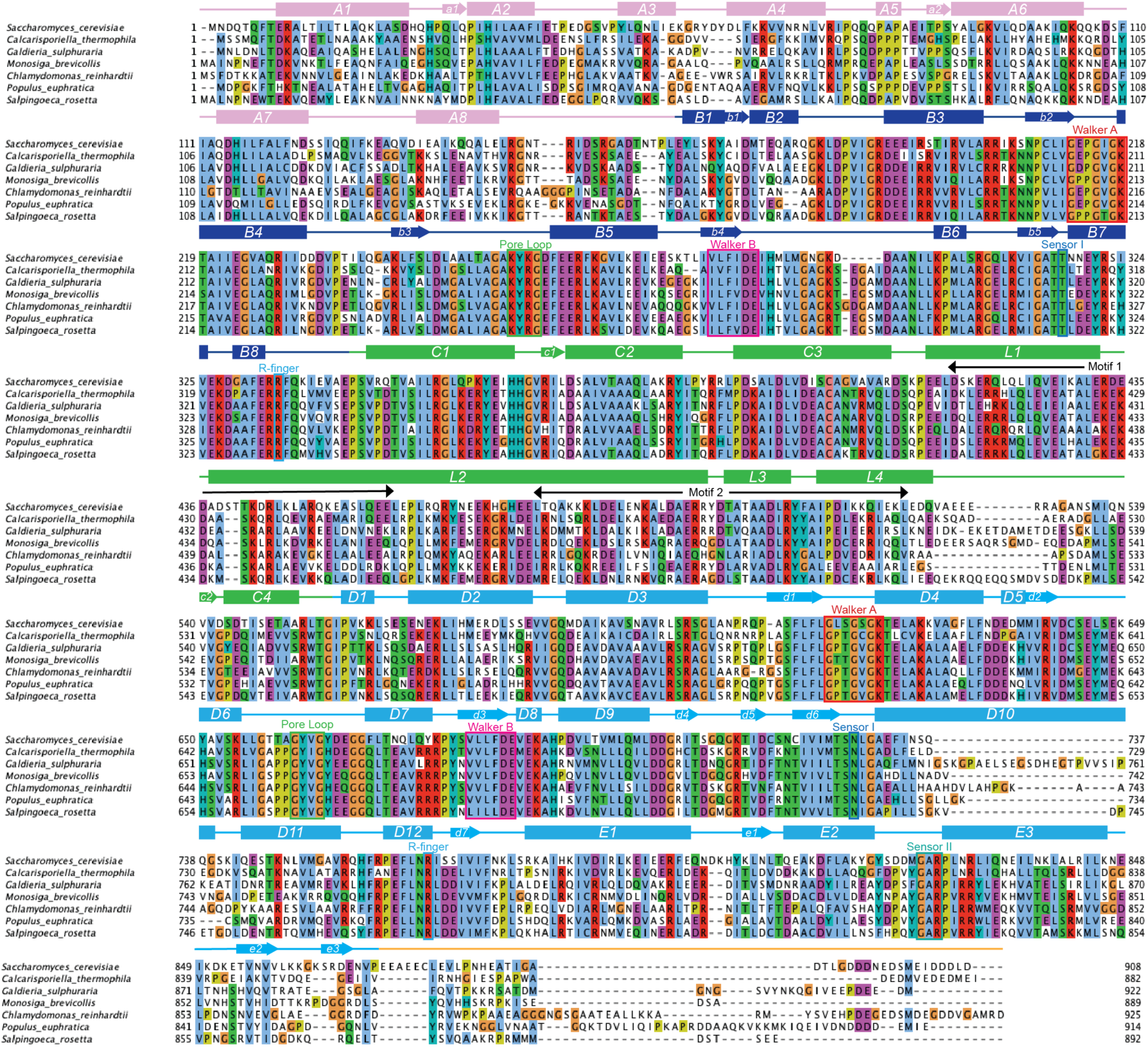
Alignment comparing ScHsp104 to Hsp104 homologs that rescue TDP-43 toxicity (related to Figure 1). Amino acid sequences of the indicated Hsp104 homologs were aligned using Clustal Omega, and the multiple sequence alignment was visualized with JalView. Positions in the alignment are colored by Clustal X convention, and structural elements of Hsp104 are indicated.

**Figure S5.**
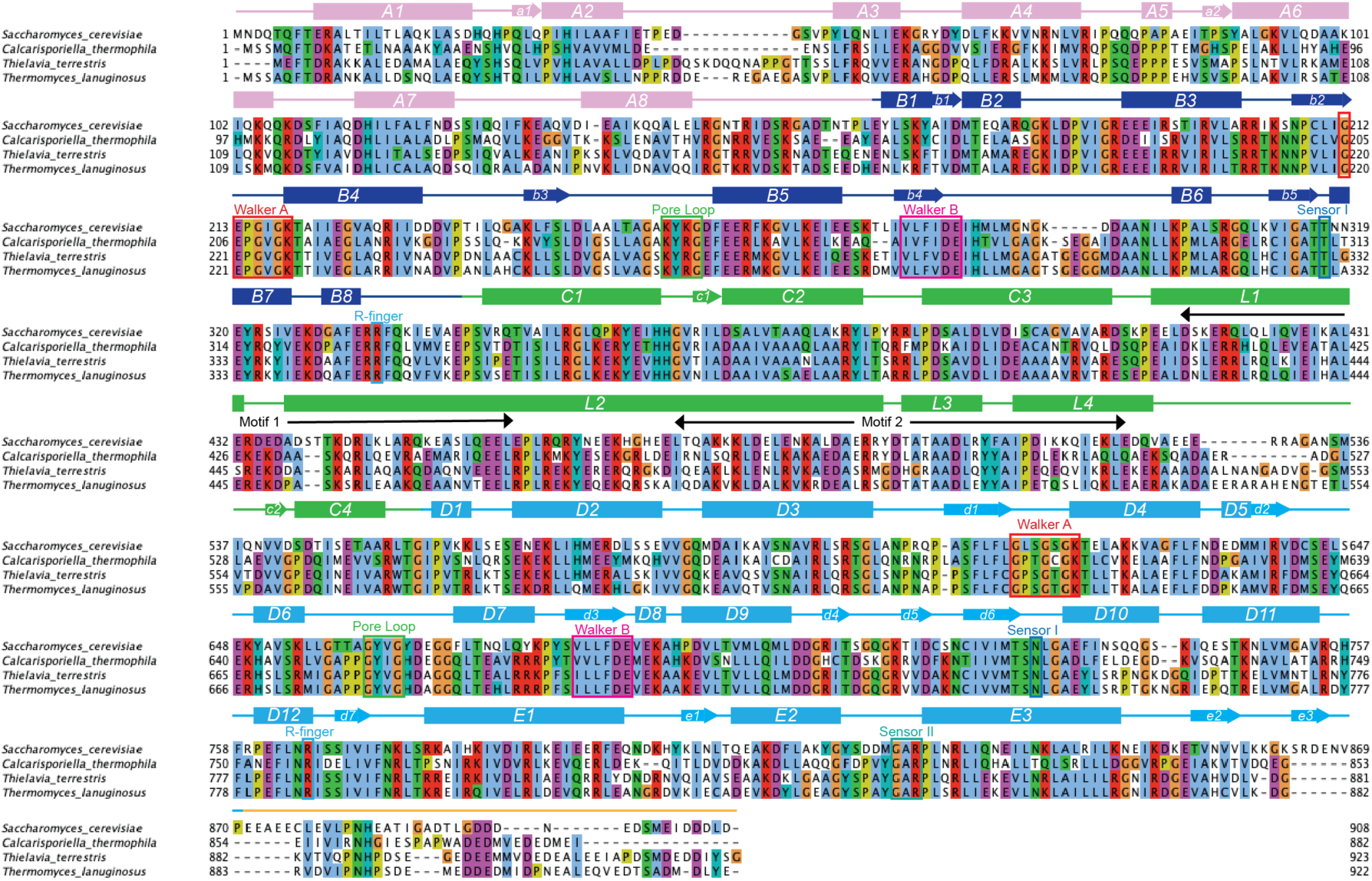
Alignment comparing ScHsp104 to Hsp104 homologs that rescue *α*Syn toxicity (related to Figure 2). Amino acid sequences of the indicated Hsp104 homologs were aligned using Clustal Omega, and the multiple sequence alignment was visualized with JalView. Amino acids are colored according to Clustal X convention, and structural elements of Hsp104 are indicated.

**Figure S6.**
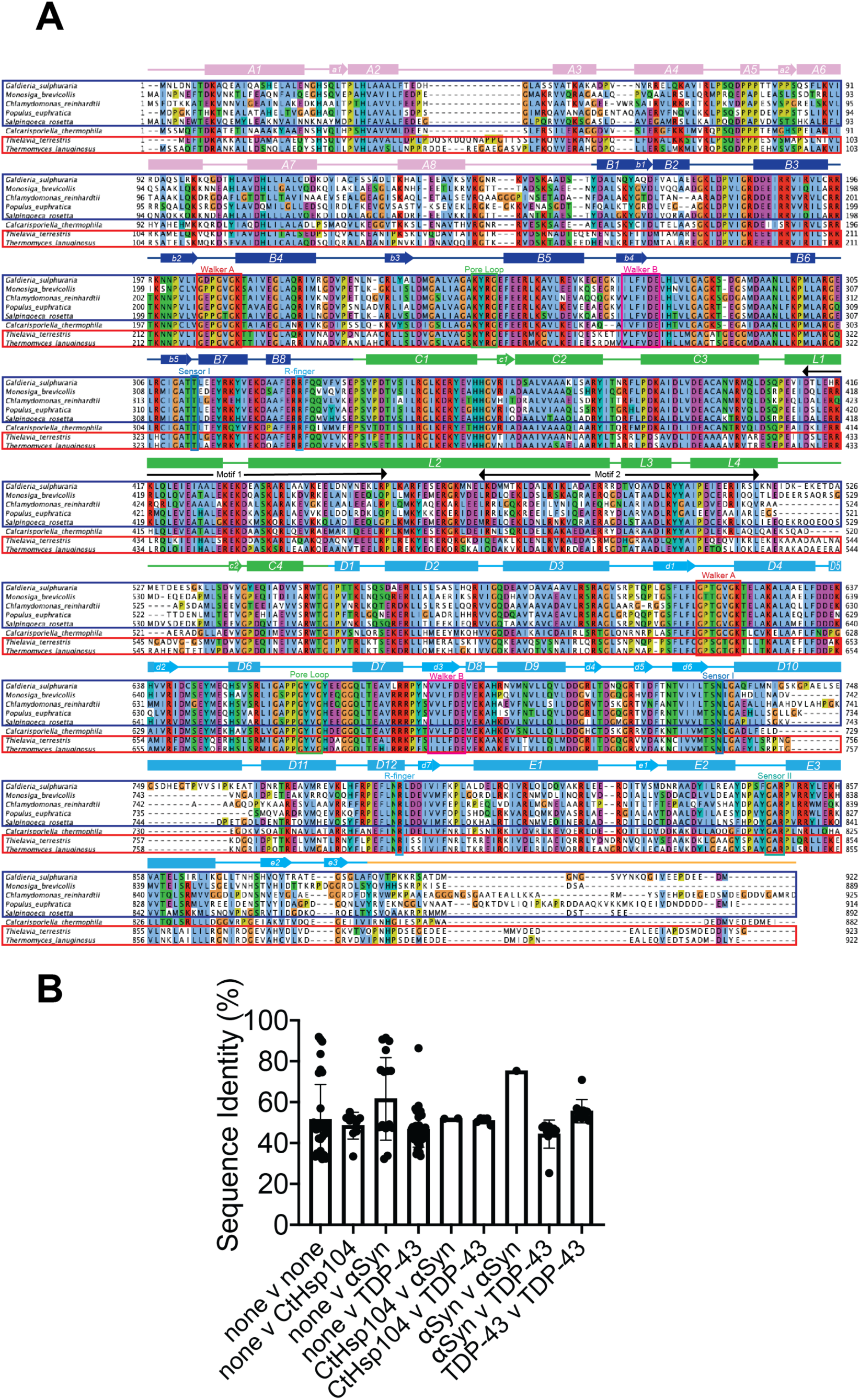
Alignment comparing Hsp104 homologs that rescue TDP-43 toxicity to those that rescue *α*Syn toxicity (related to Figures 1 and 2). **(A)** Amino acid sequences of the indicated Hsp104 homologs were aligned using Clustal Omega, and the multiple sequence alignment was visualized with JalView. Amino acids are colored according to Clustal X convention, and structural elements of Hsp104 are indicated. TDP-43-specific Hsp104 homologs are boxed in blue and alpha-synuclein-specific homologs are boxed in red. **(B)** Bar chart comparing average pairwise sequence identities between Hsp104 homologs with different toxicity suppression phenotypes.

**Figure S7.**
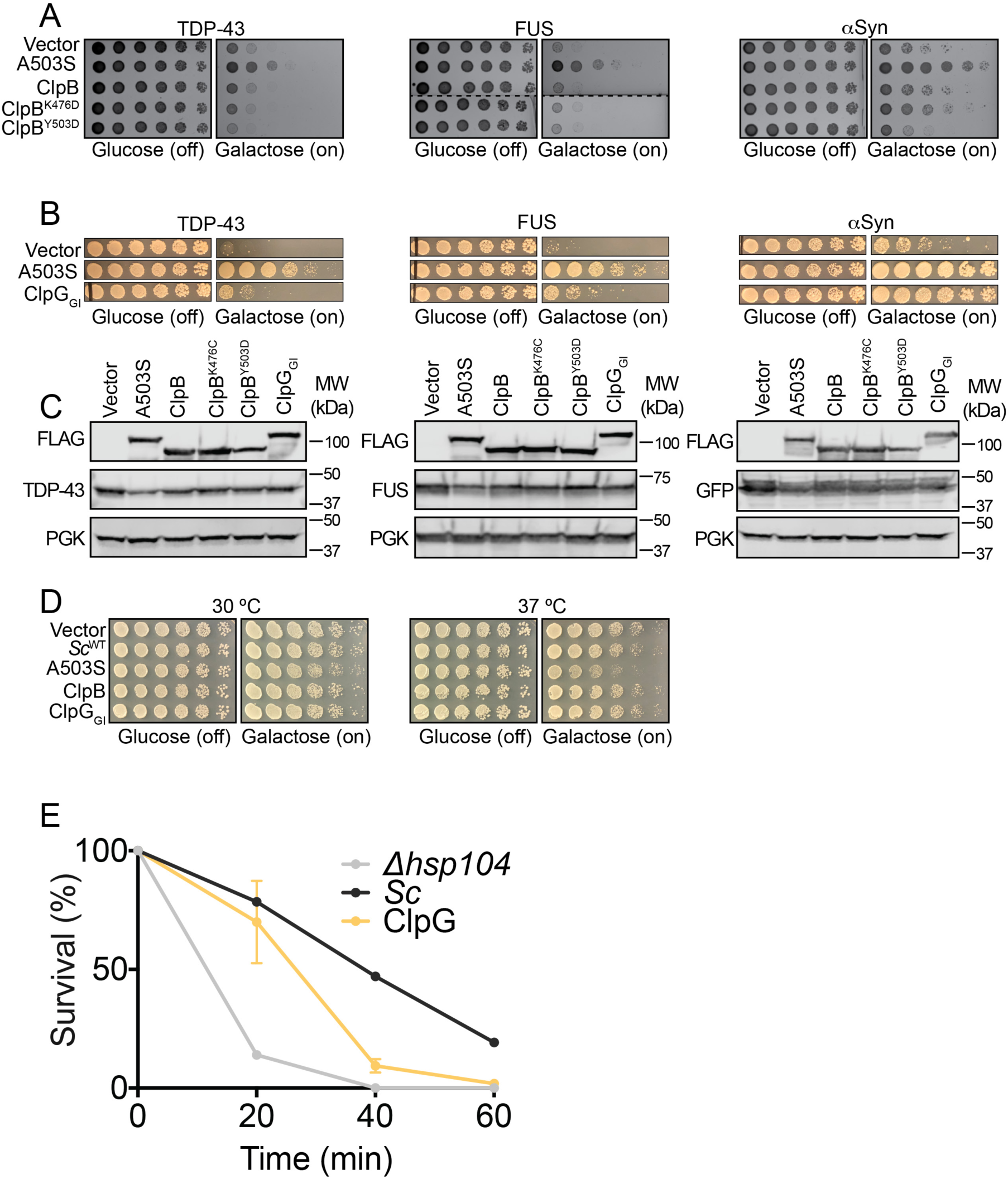
ClpB does not suppress TDP-43, FUS, or *α*Syn toxicity whereas ClpG_GI_ robustly suppresses *α*Syn toxicity and confers thermotolerance to *Δhsp104* yeast (related to Figures 1 and 2). **(A)** Spotting assay demonstrating that neither ClpB nor hyperactive ClpB variants suppress TDP-43, FUS, or *α*Syn toxicity. Dashed line indicates splicing of FUS plate. **(B)** Spotting assay demonstrating slight suppression of TDP-43 and FUS toxicity, and robust suppression of *α*Syn toxicity by ClpG_GI_. **(C)** Western blots show consistent expression of FLAG-tagged ClpB, ClpB variants, and ClpG in yeast. These Hsp104s also do not affect TDP-43, FUS, or *α*Syn levels in yeast. **(D)** Neither ClpB nor ClpG_GI_ cause temperature-dependent toxicity when expressed in yeast. **(E)** *Δhsp104* yeast carrying a plasmid encoding empty vector or the indicated Hsp104 homolog were pre-treated at 37 °C for 30 min, treated at 50 °C for 0-60 min, and plated. Surviving colonies were quantified after 2d recovery. Values represent means ±SEM (n=3 independent transformations).

**Figure S8.**
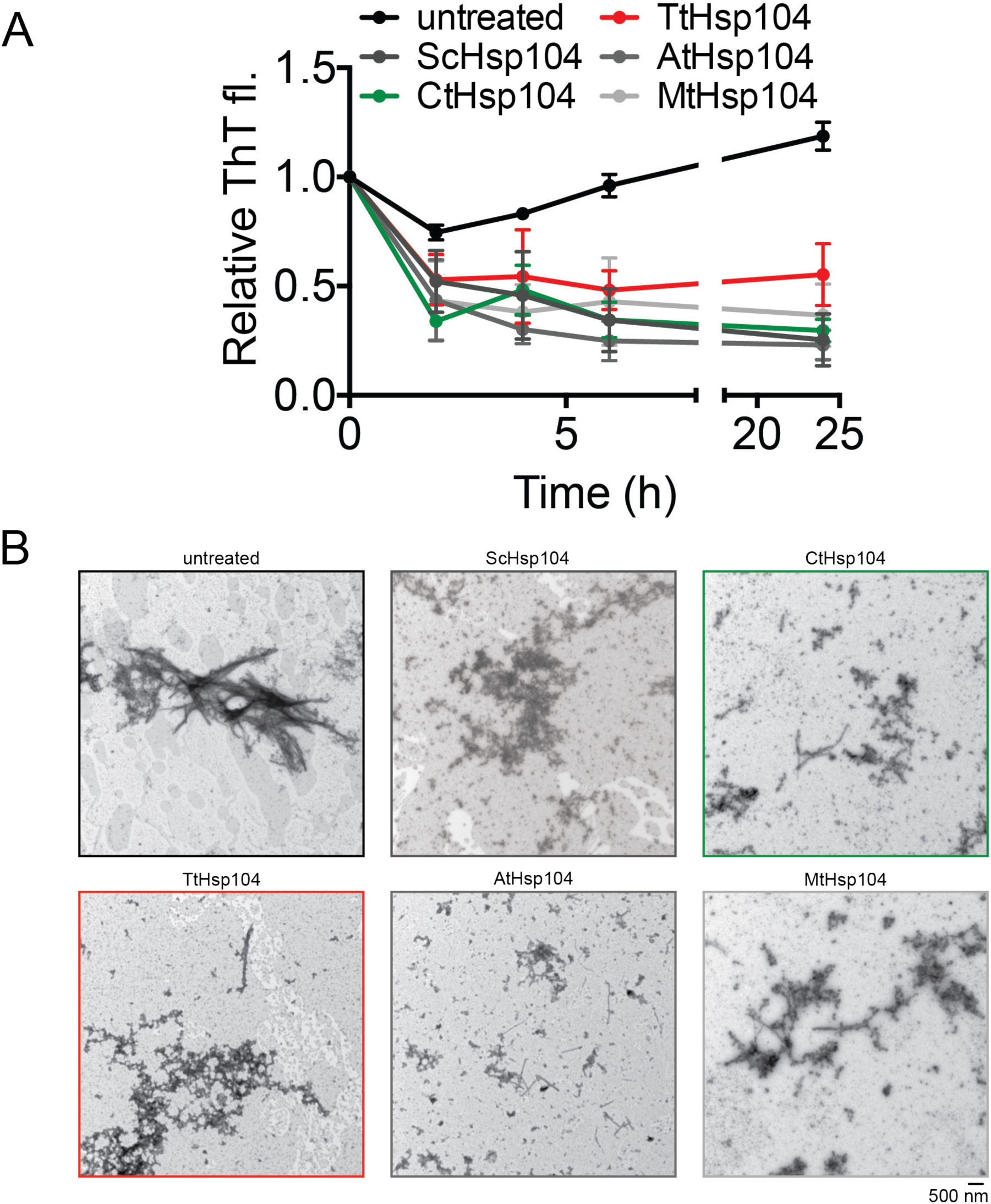
Hsp104 homologs remodel SEVI fibers (related to Figure 5). **(A)** SEVI fibrils (20 µM monomer) were incubated with buffer (untreated) or the indicated Hsp104 homolog (3 µM) for 0-24 h. Fibril integrity was assessed by ThT fluorescence. Values represent means ± SEM (n=3). **(B)** Representative EM images of SEVI fibrils incubated with buffer (untreated) or the indicated Hsp104 homolog (3 µM) for 3 h. Scale bar is indicated (bottom right of gallery).

**Figure S9.**
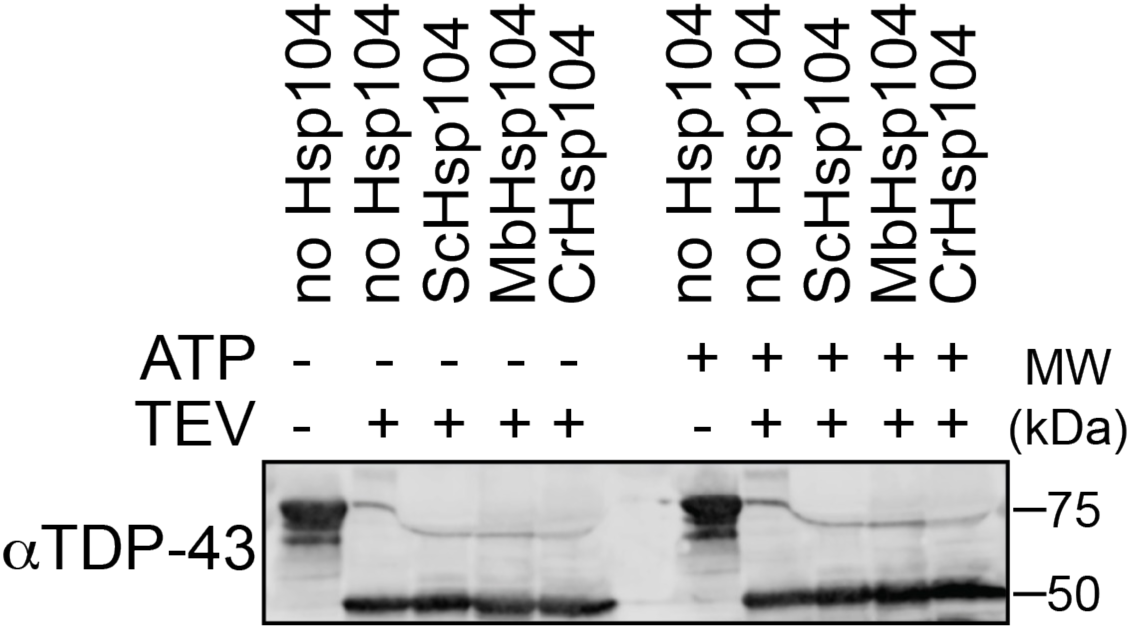
Hsp104 homologs do not affect MBP-TEV-TDP43 cleavage by TEV protease (related to Figure 5). Western blot of MBP-TEV-TDP43 incubated either alone, or in combination with TEV protease, Hsp104s, and ATP, as indicated above lanes. Molecular weight markers are indicated (right).

**Figure S10.**
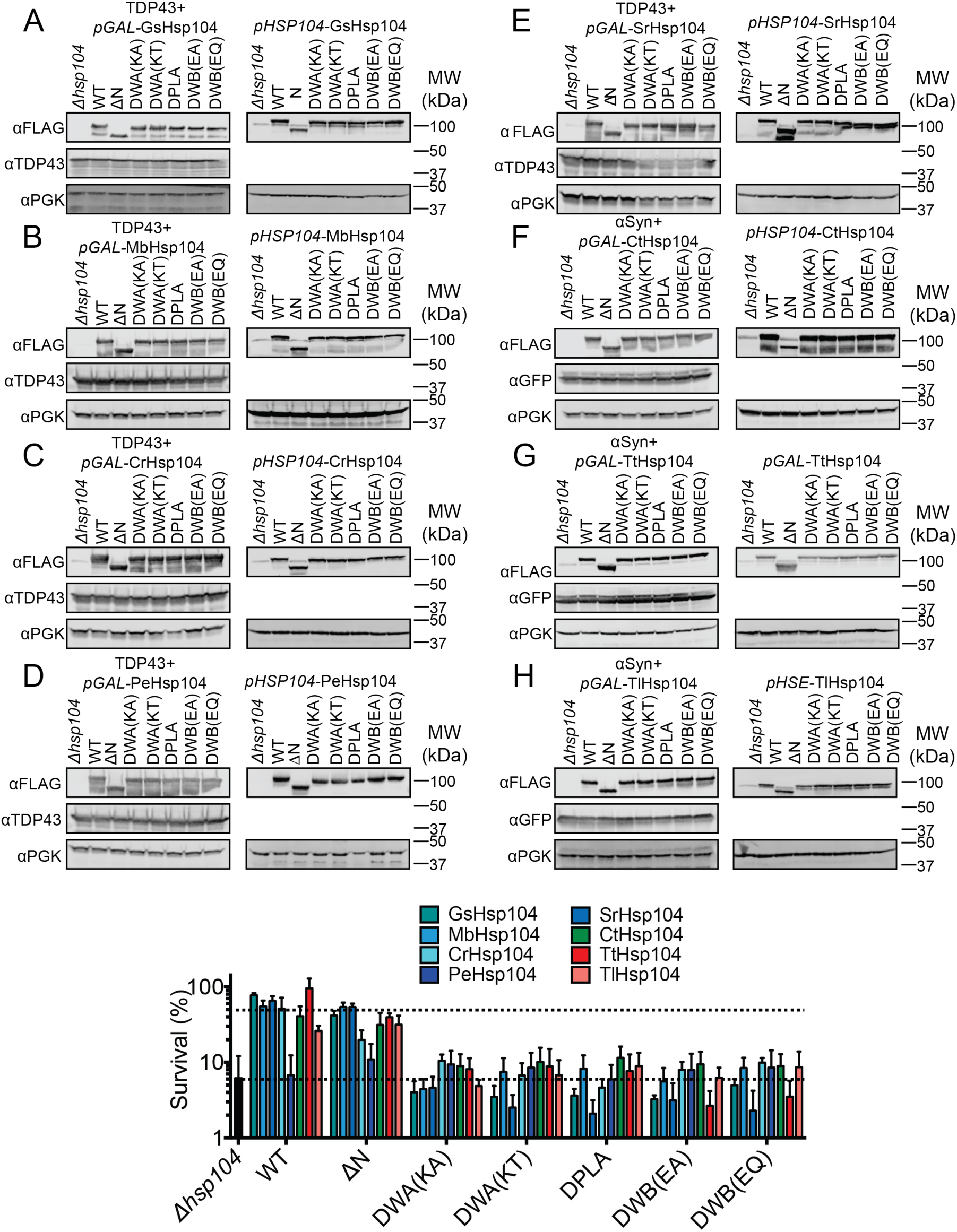
Thermotolerance of Hsp104 mutants are defective in thermotolerance and are consistently expressed (related to Figure 6). **(A-H)** Western blots show consistent expression of all homologs and mutants, both from p*GAL* and p*HSP104*. Molecular weight markers are indicated (right). **(I)** Survival of yeast expressing GsHsp104, MbHsp104, CrHsp104, PeHsp104, SrHsp104, CtHsp104, TtHsp104, TlHsp104, or the indicated chimera was assessed following 20 min heat shock at 50 °C. Values represent means ±SEM (n=3 independent transformations). Top dashed line indicates average survival of strains expressing wild-type Hsp104s, while bottom dashed line indicates average survival of *Δhsp104* cells.

**Figure S11.**
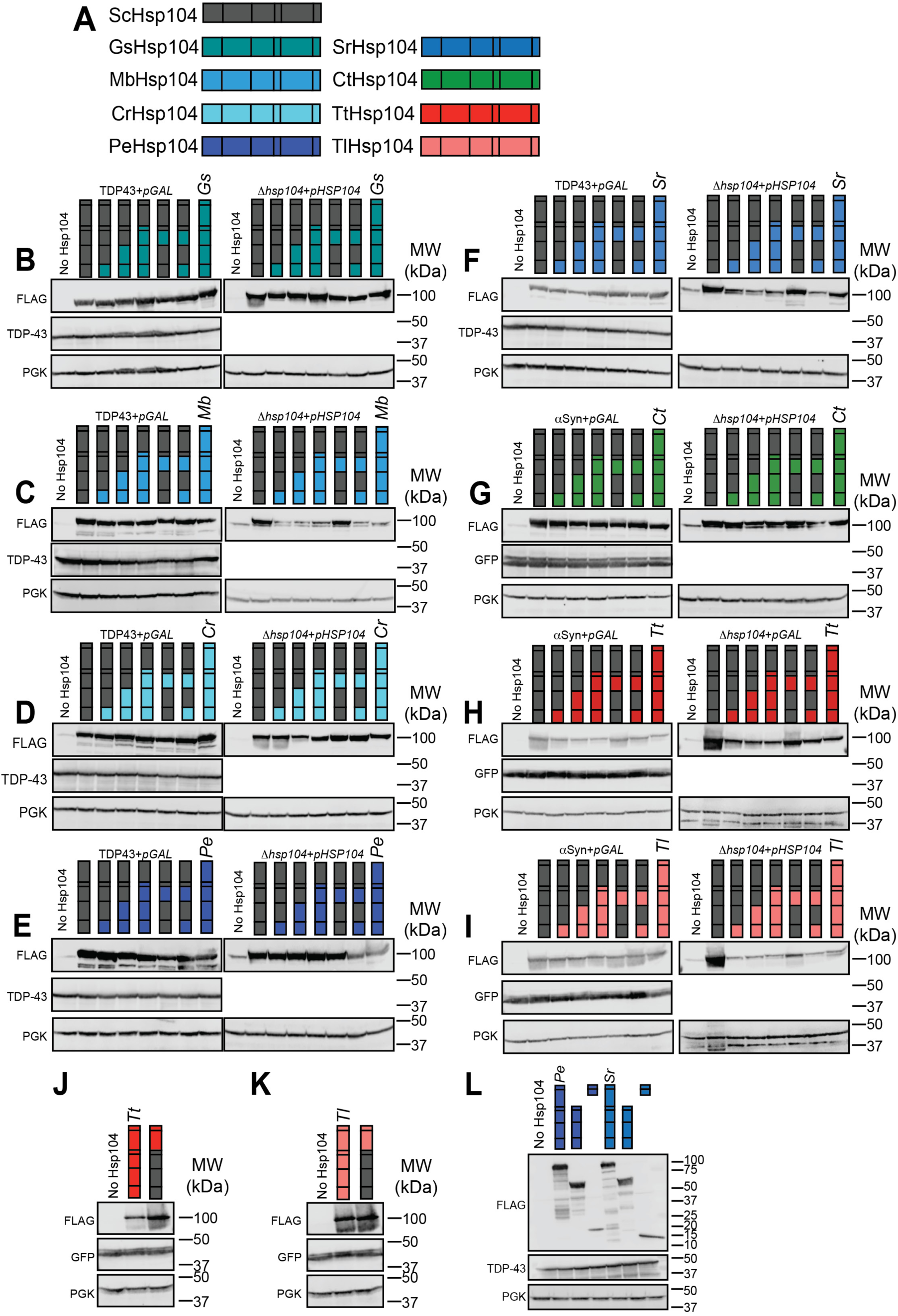
Chimeric Hsp104s and proteotoxic substrates are consistently expressed in yeast (related to Figure 7). **(A)** Color code of Hsp104 homologs, and schematic of Hsp104 domain organization. **(B-K)** Western blots show that chimeras between ScHsp104 and GsHsp104 (B), MbHsp104 (C), CrHsp104 (D), PeHsp104 (E), SrHsp104 (F), CtHsp104 (G), TtHsp104 (H,J) and TlHsp104 (I,K) are expressed consistently and at a level comparable to wild-type protein. TDP-43 (B-F) and *α*Syn (G-I) are expressed consistently in all strains. Molecular weight markers are indicated at right of each panel. **(L)** Western blot showing expression of PeHsp104, SrHsp104, or fragments derived from these homologs in TDP-43-expressing yeast strains. Molecular weight markers are indicated at the right of the panel.

**Figure S12.**
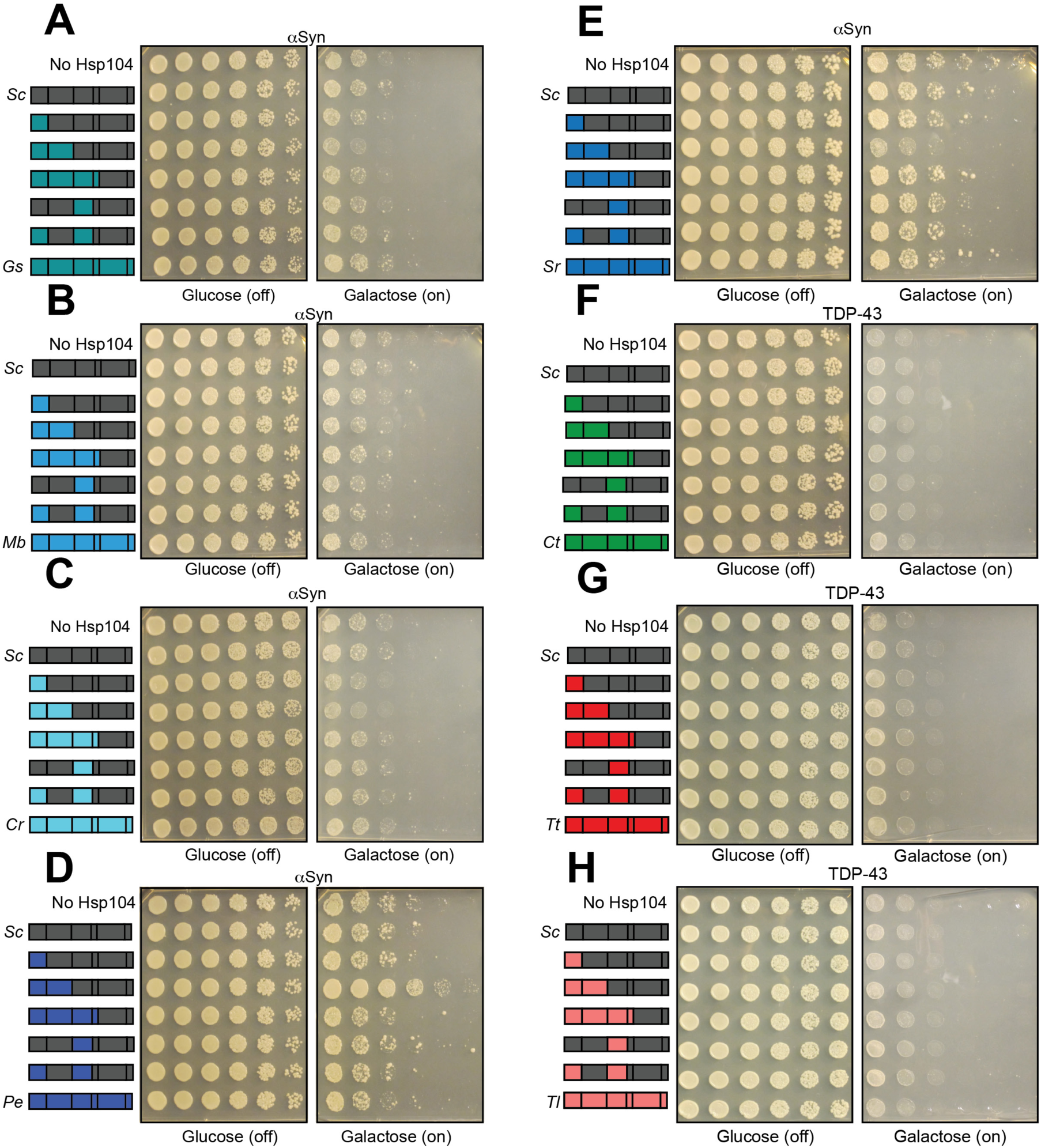
Characterization of Hsp104 chimera specificity (related to Figure 7). **(A)** Spotting assay of *Δhsp104* strains coexpressing *α*Syn and the indicated chimeras between ScHsp104 and GsHsp104 shows that none of these chimeras suppress *α*Syn toxicity. **(B)** Spotting assay of *Δhsp104* strains coexpressing *α*Syn and the indicated chimeras between ScHsp104 and MbHsp104 shows that none of these chimeras suppress *α*Syn toxicity. **(C)** Spotting assay of *Δhsp104* strains coexpressing *α*Syn and the indicated chimeras between ScHsp104 and CrHsp104 shows that none of these chimeras suppress *α*Syn toxicity. **(D)** Spotting assay of *Δhsp104* strains coexpressing *α*Syn and the indicated chimeras between ScHsp104 and PeHsp104 shows that a chimera possessing the NTD and NBD1 from PeHsp104 suppresses *α*Syn toxicity. **(E)** Spotting assay of *Δhsp104* strains coexpressing *α*Syn and the indicated chimeras between ScHsp104 and SrHsp104 shows that none of these chimeras suppress *α*Syn toxicity. **(F)** Spotting assay of *Δhsp104* strains coexpressing TDP-43 and the indicated chimeras between ScHsp104 and CtHsp104 shows that none of these chimeras suppress TDP-43 toxicity. **(G)** Spotting assay of *Δhsp104* strains coexpressing TDP-43 and the indicated chimeras between ScHsp104 and TtHsp104 shows that none of these chimeras suppress TDP-43 toxicity. **(H)** Spotting assay of *Δhsp104* strains coexpressing TDP-43 and the indicated chimeras between ScHsp104 and TlHsp104 shows that none of these chimeras suppress TDP-43 toxicity.

**Figure S13.**
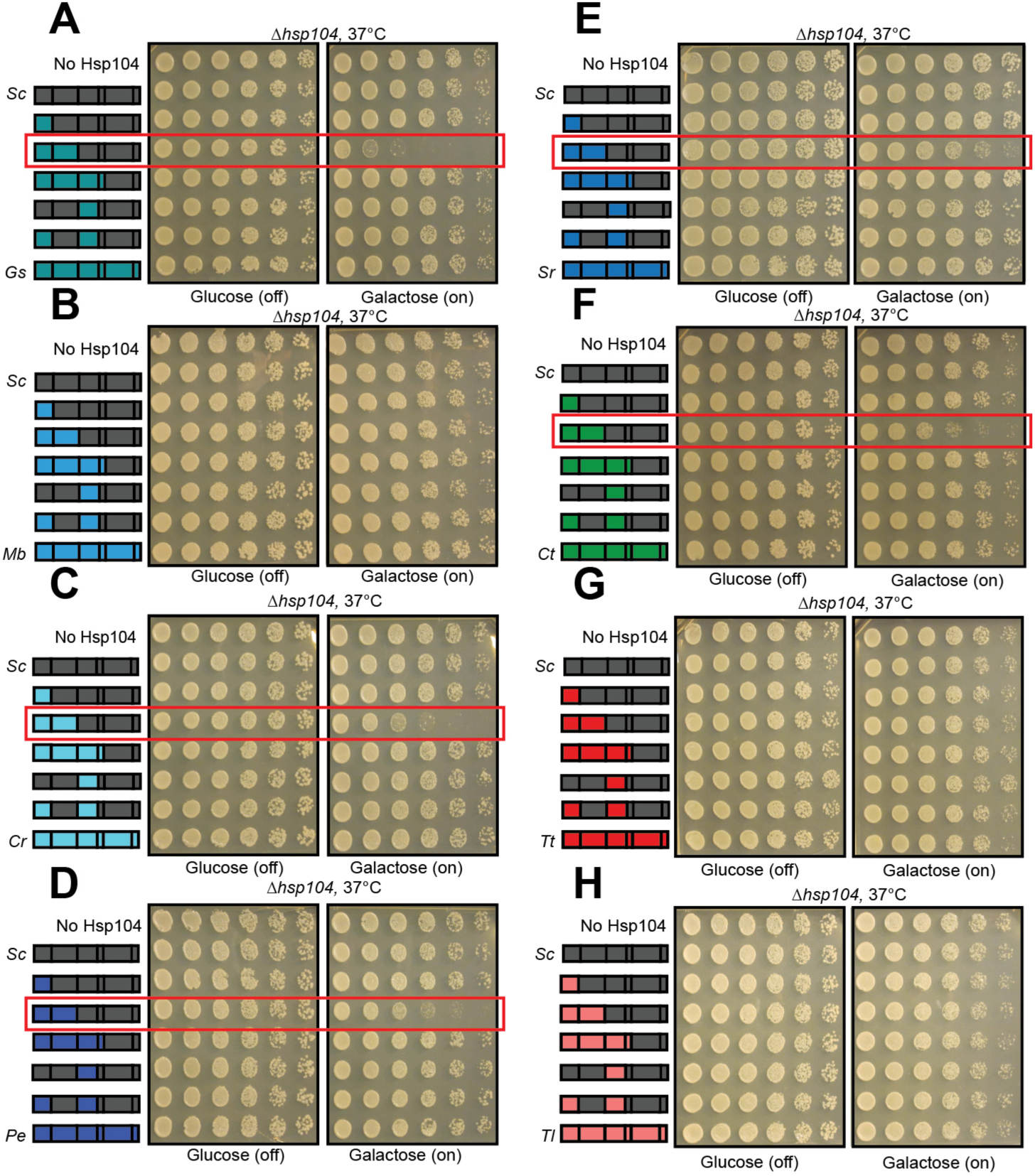
Characterization of chimera intrinsic toxicity (related to Figure 7). Strains expressing chimeras between ScHsp104 and GsHsp104 (A), MbHsp104 (B), CrHsp104 (C), PeHsp104 (D), SrHsp104 (E), CtHsp104 (F), TtHsp104 (G), and TlHsp104 (H) were spotted onto glucose and galactose media, and incubated at 37 °C for 2-3 days to observe growth. In several cases (panels A, C, D, E, and F) chimeras possessing non-cognate NTD:NBD1 units displayed toxicity.

**Figure S14.**
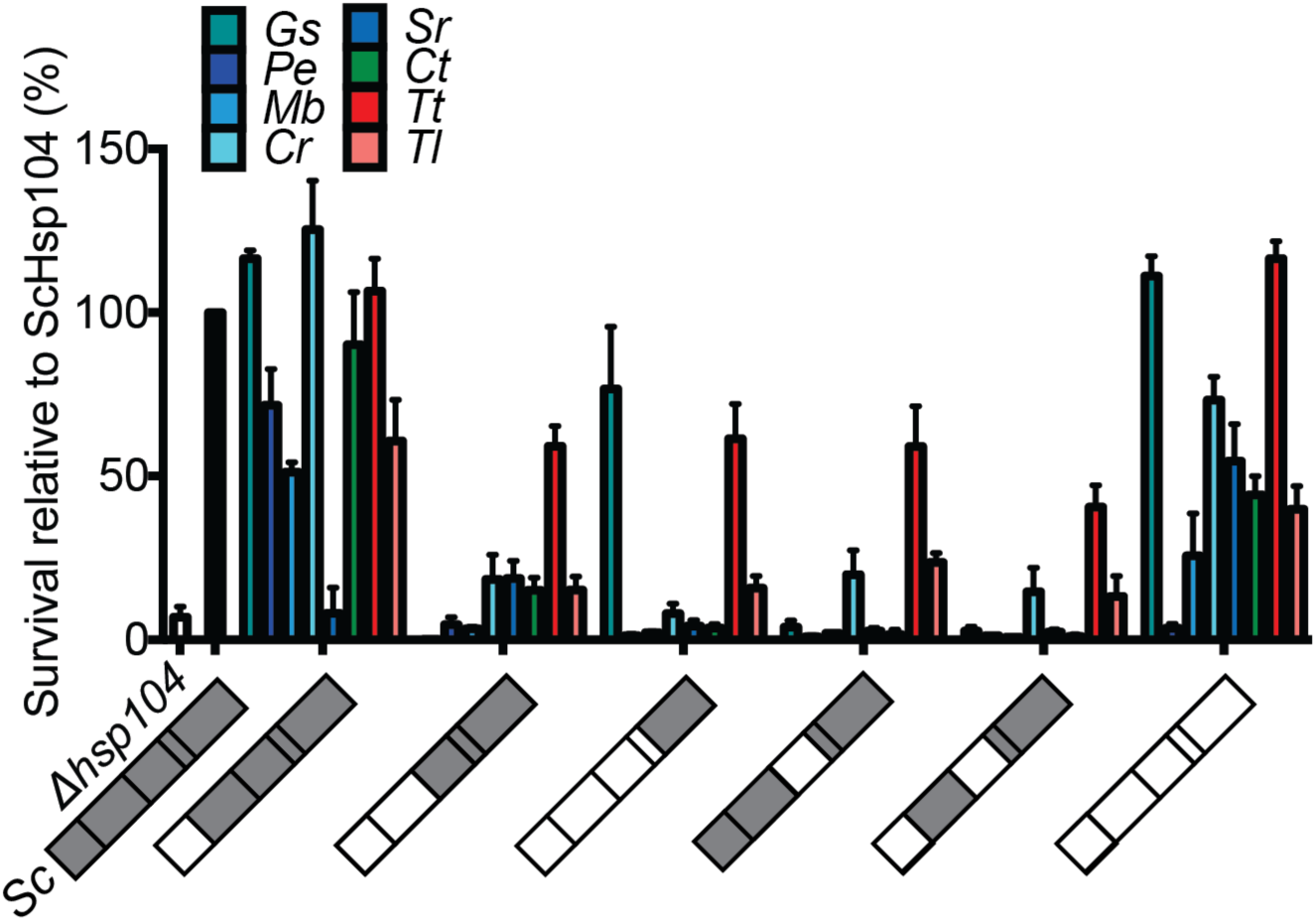
Thermotolerance activity of Hsp104 chimeras (related to Figure 7). Survival of yeast expressing GsHsp104, MbHsp104, CrHsp104, PeHsp104, SrHsp104, CtHsp104, TtHsp104, TlHsp104, or a chimera between one of these (as indicated by color in bar chart) and ScHsp104 was assessed following 20 min heat shock at 50 °C. Values are relative to wild-type ScHsp104 (single black bar at left) and represent means ±SEM (n=3 independent transformations). Illustrations on x-axis indicate chimera composition. Gray indicates portions of the chimera from ScHsp104 and white indicates variable regions from other homologs

**Table S1.**
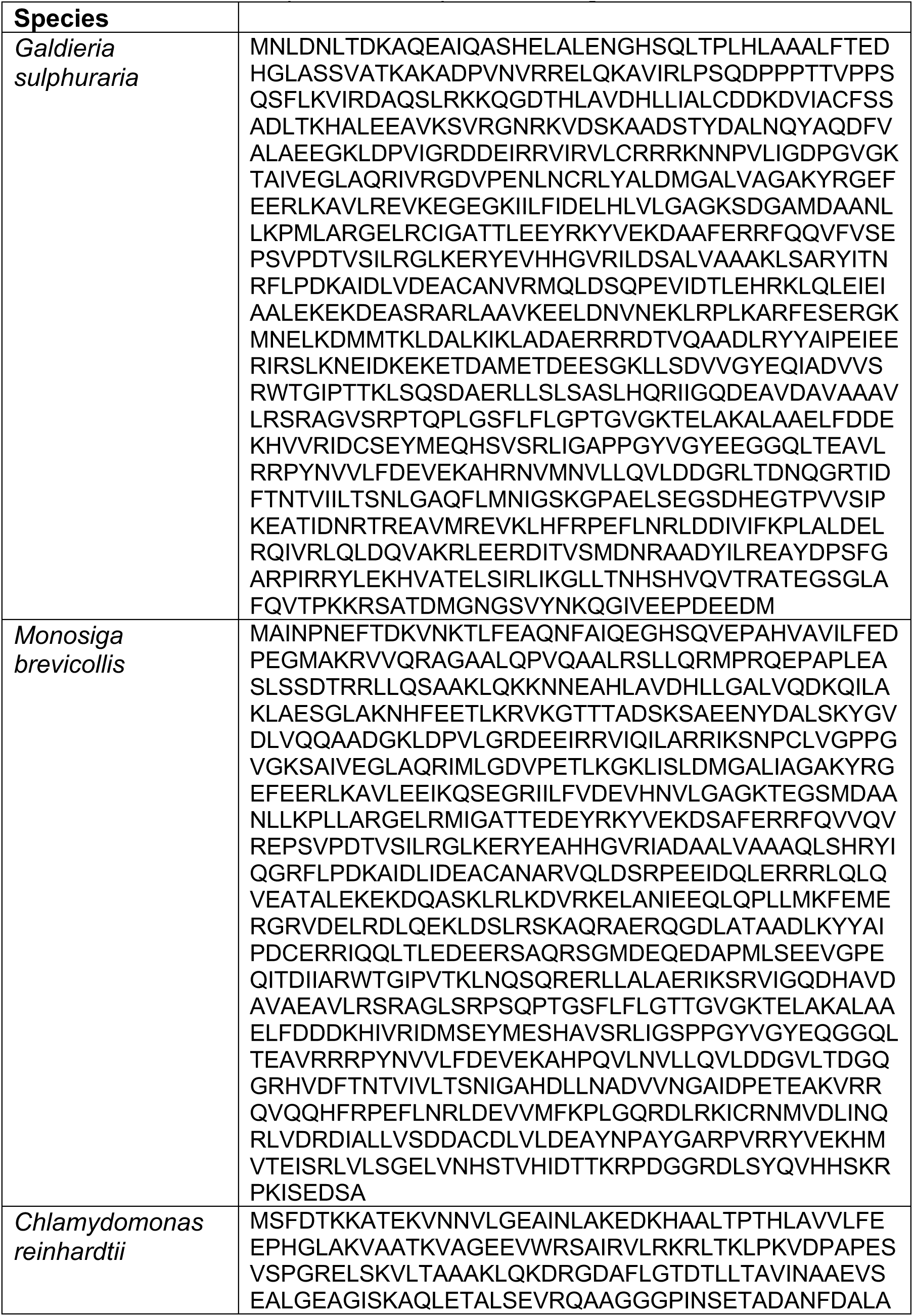
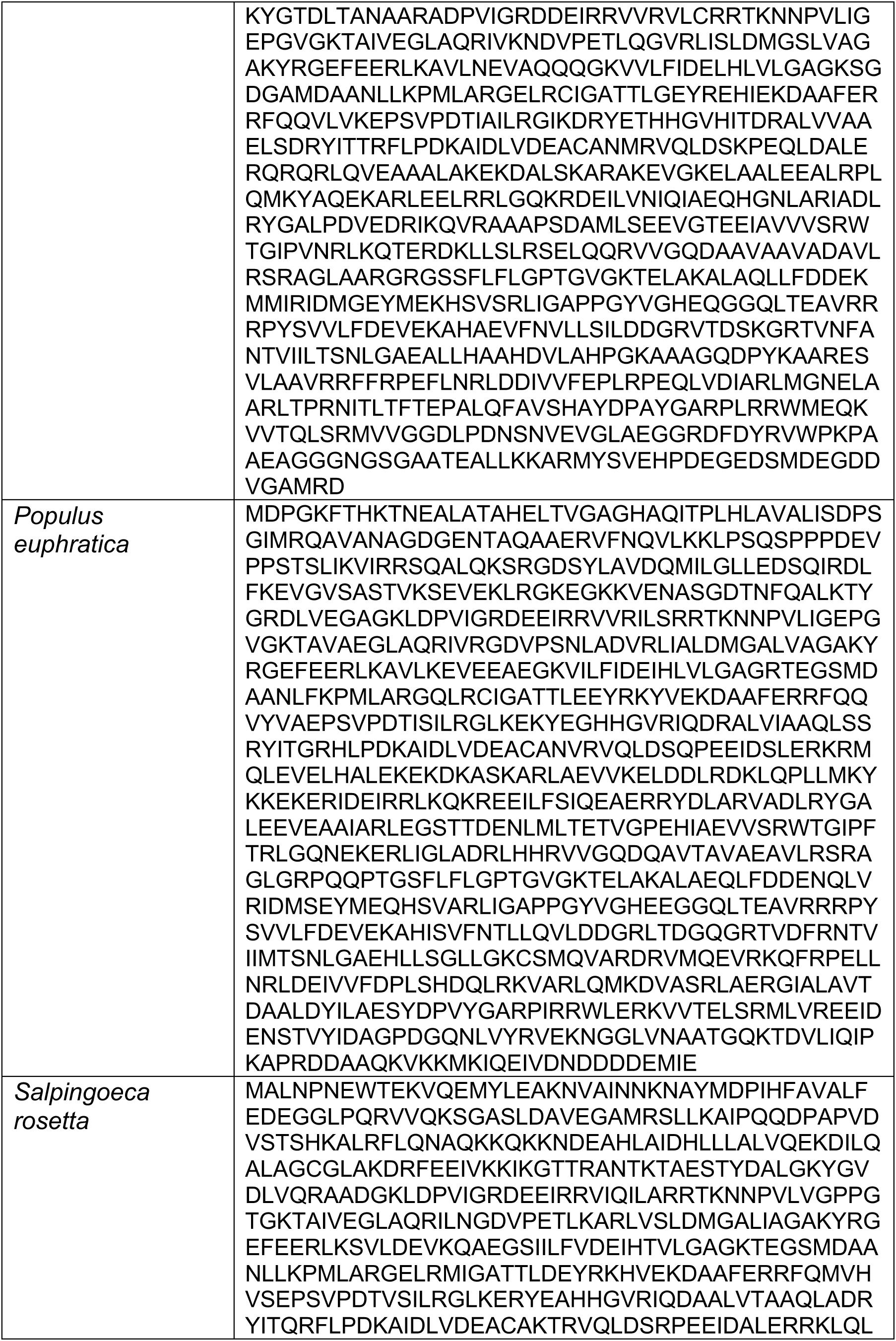
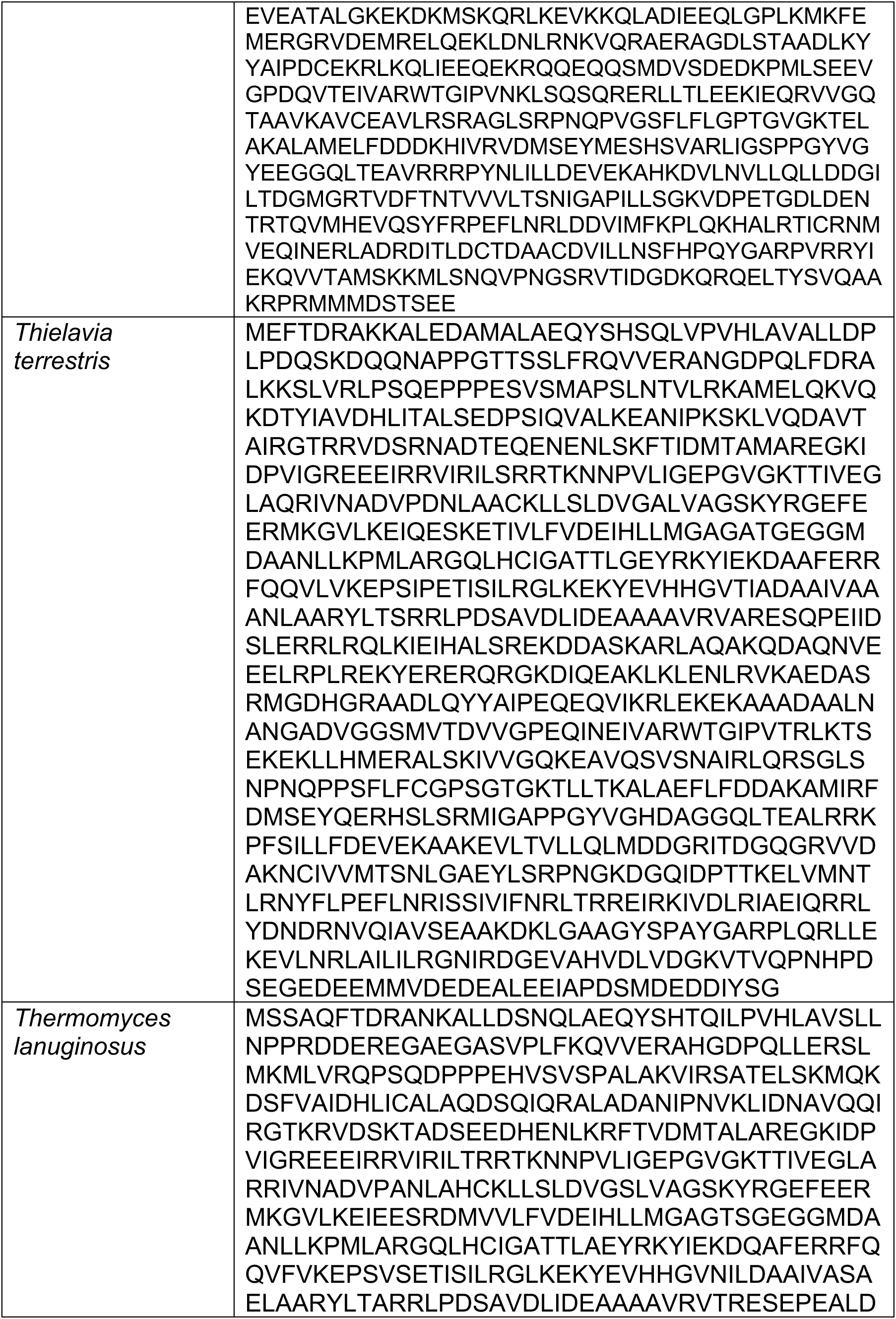
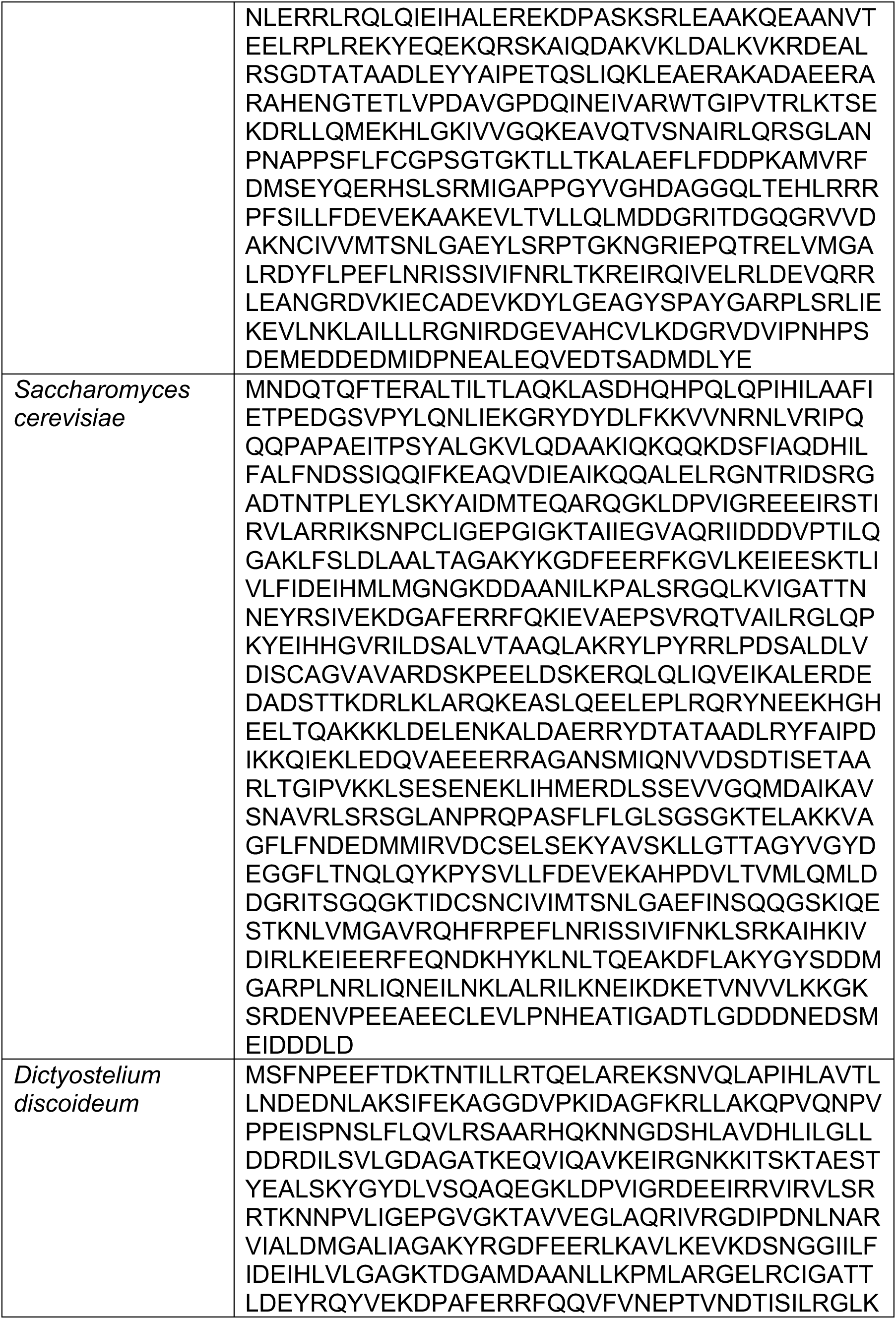
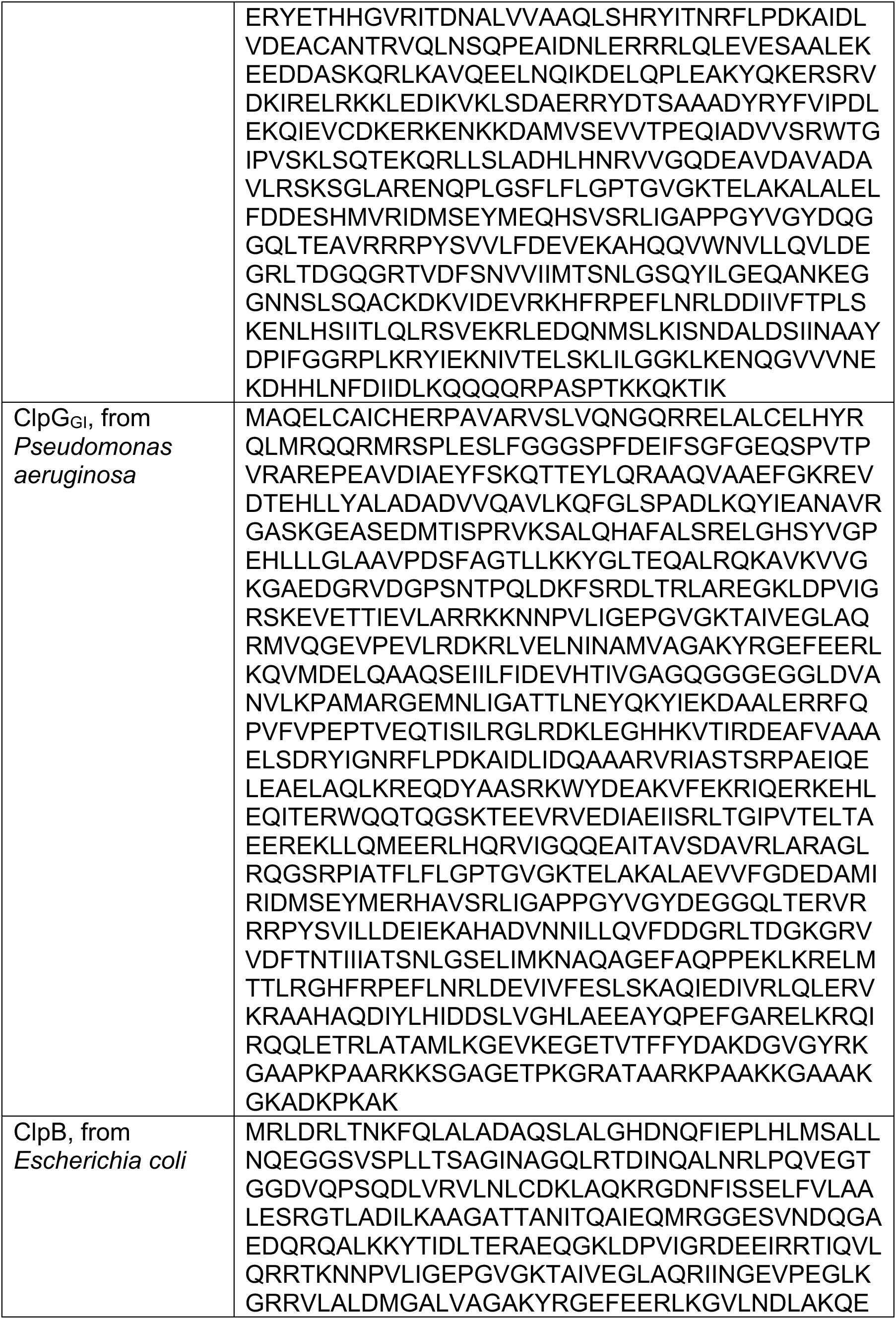
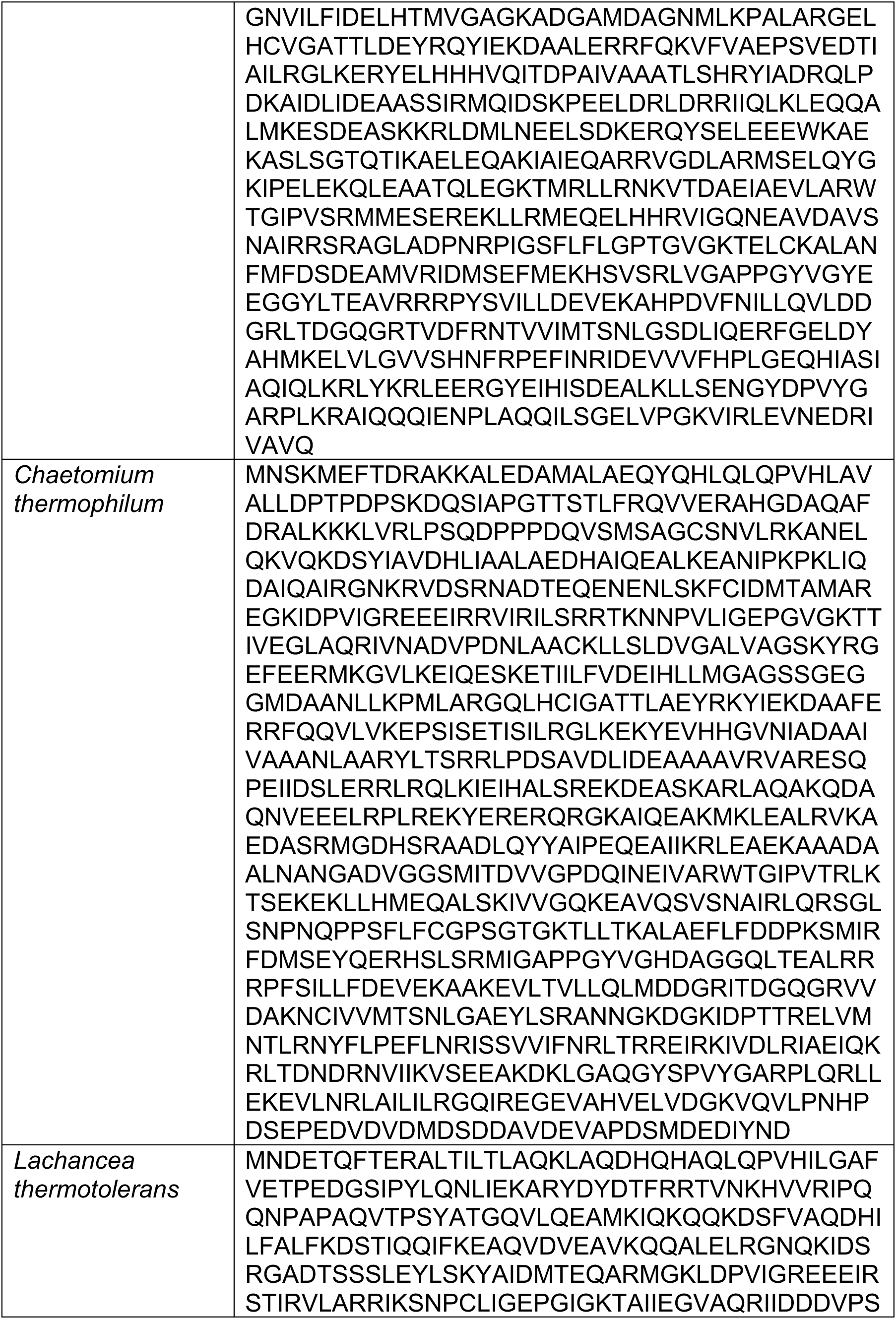
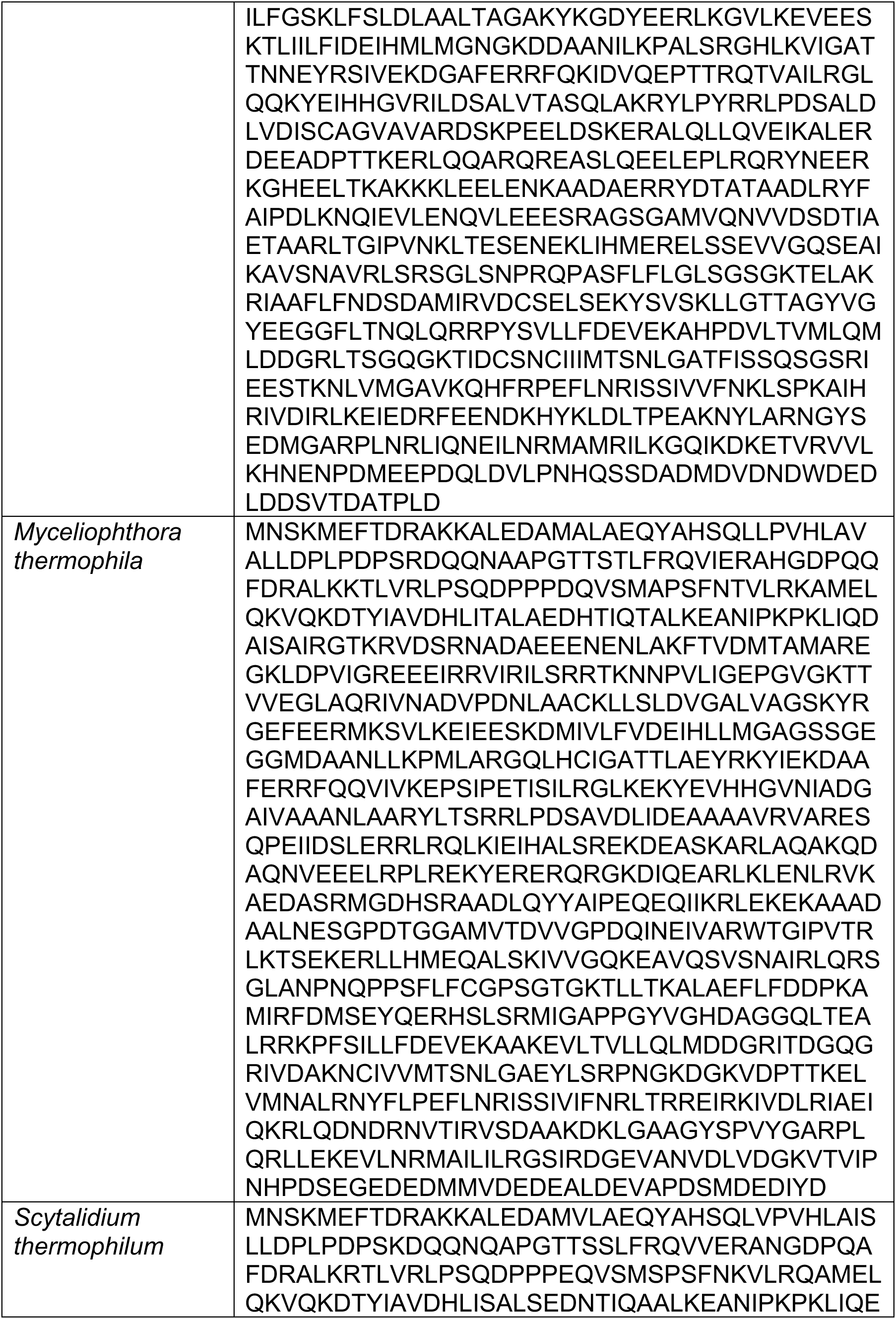
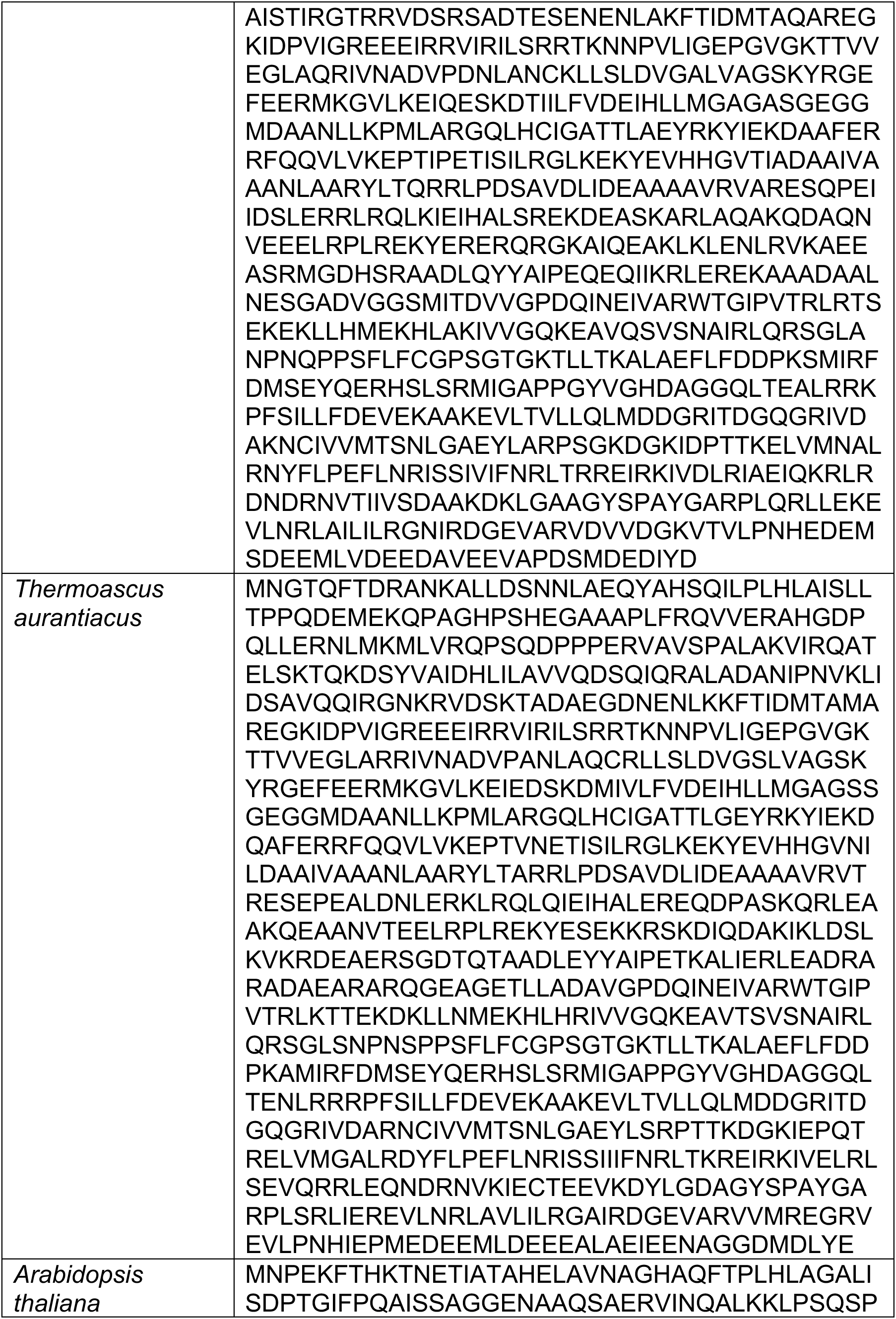
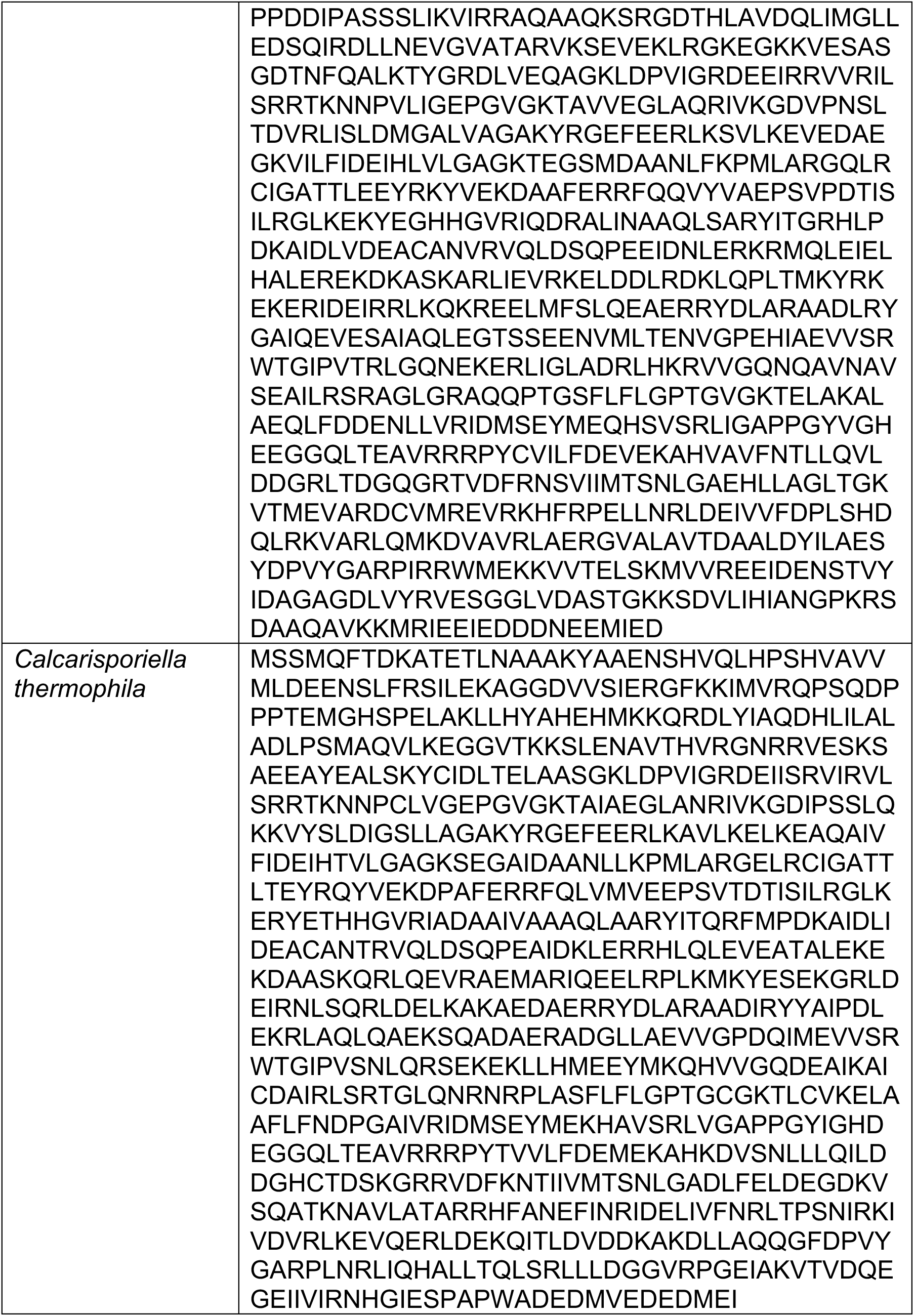
Amino acid sequences of Hsp104 homologs.

**Table S2.**
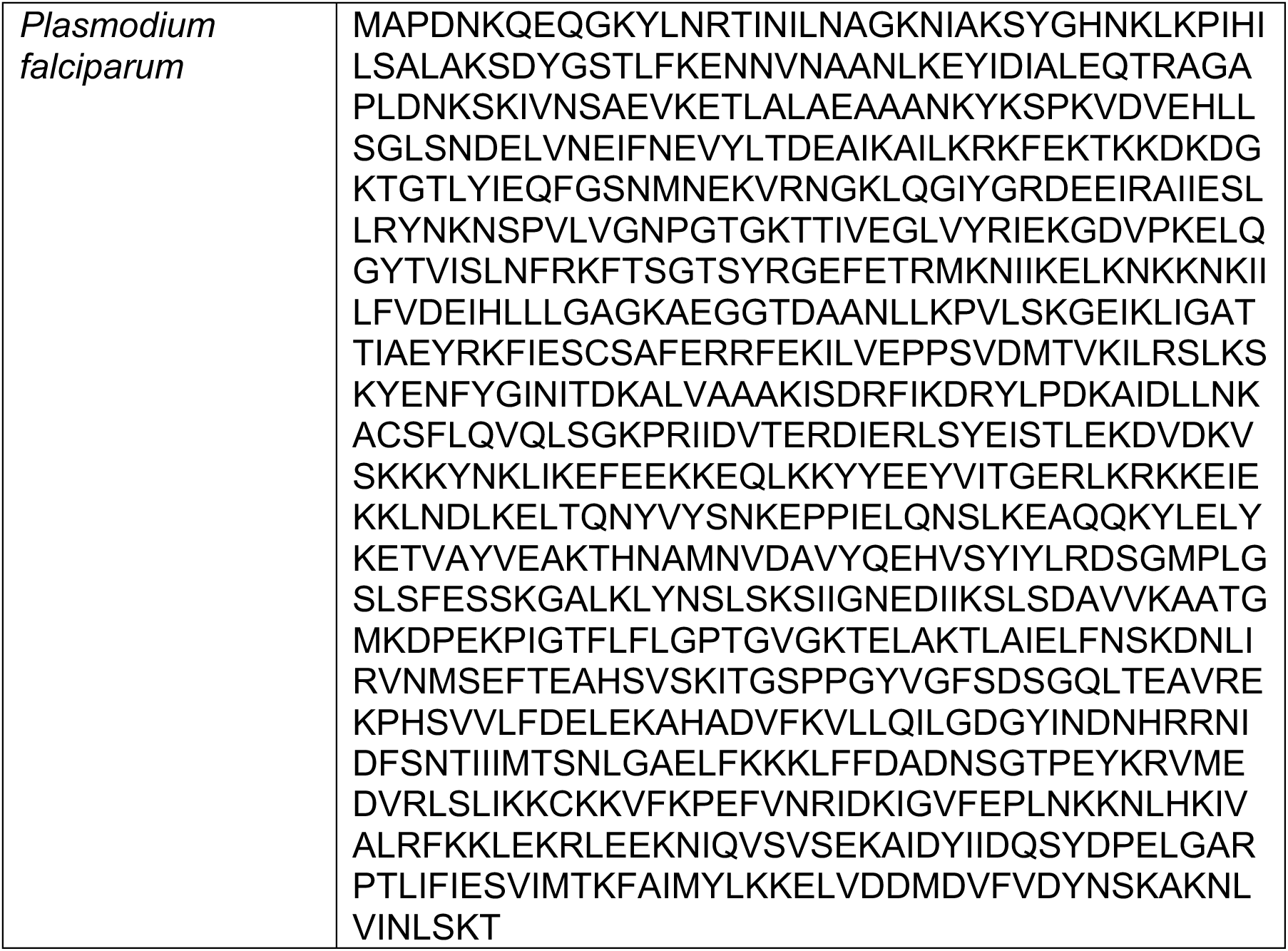
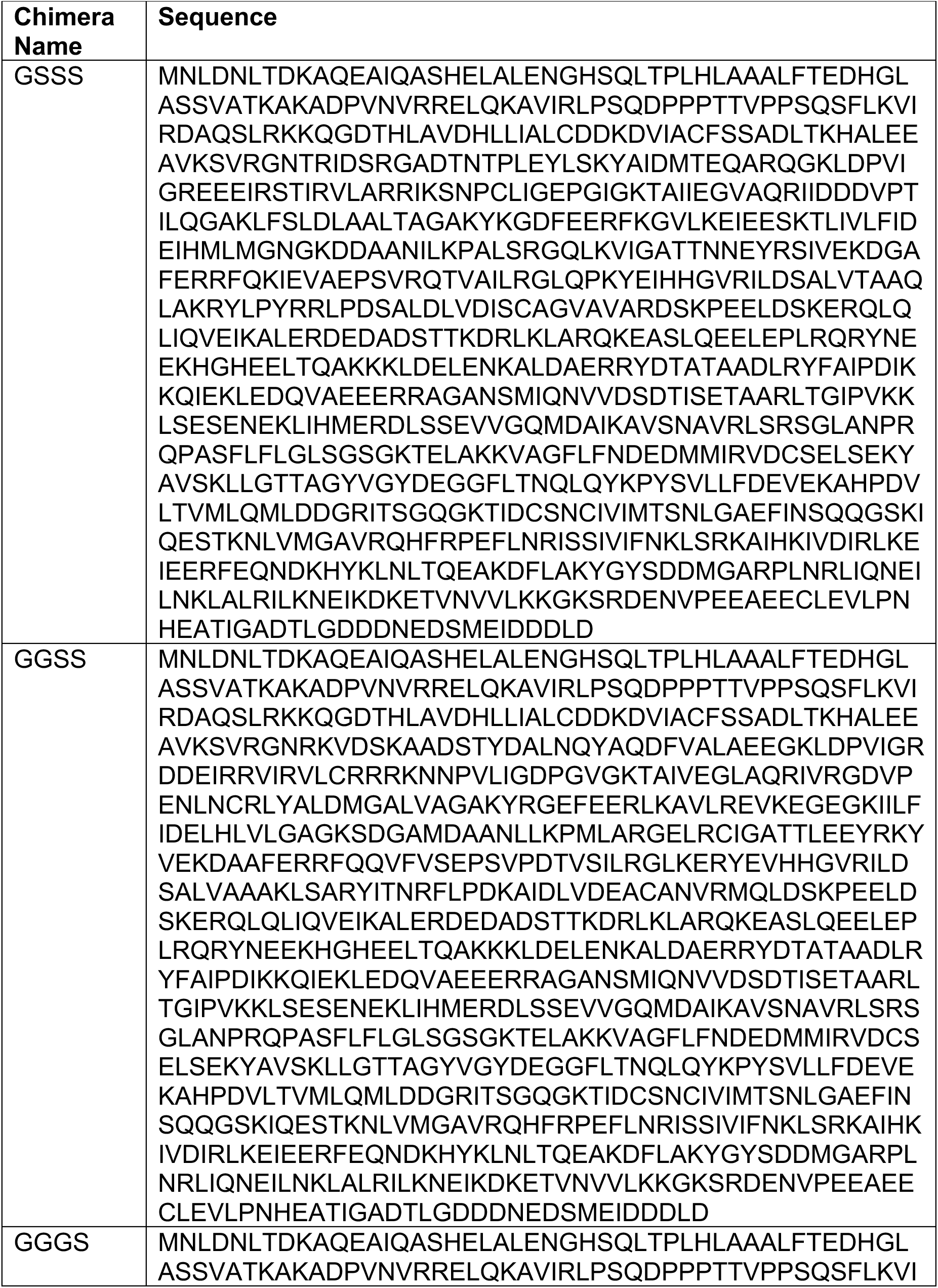
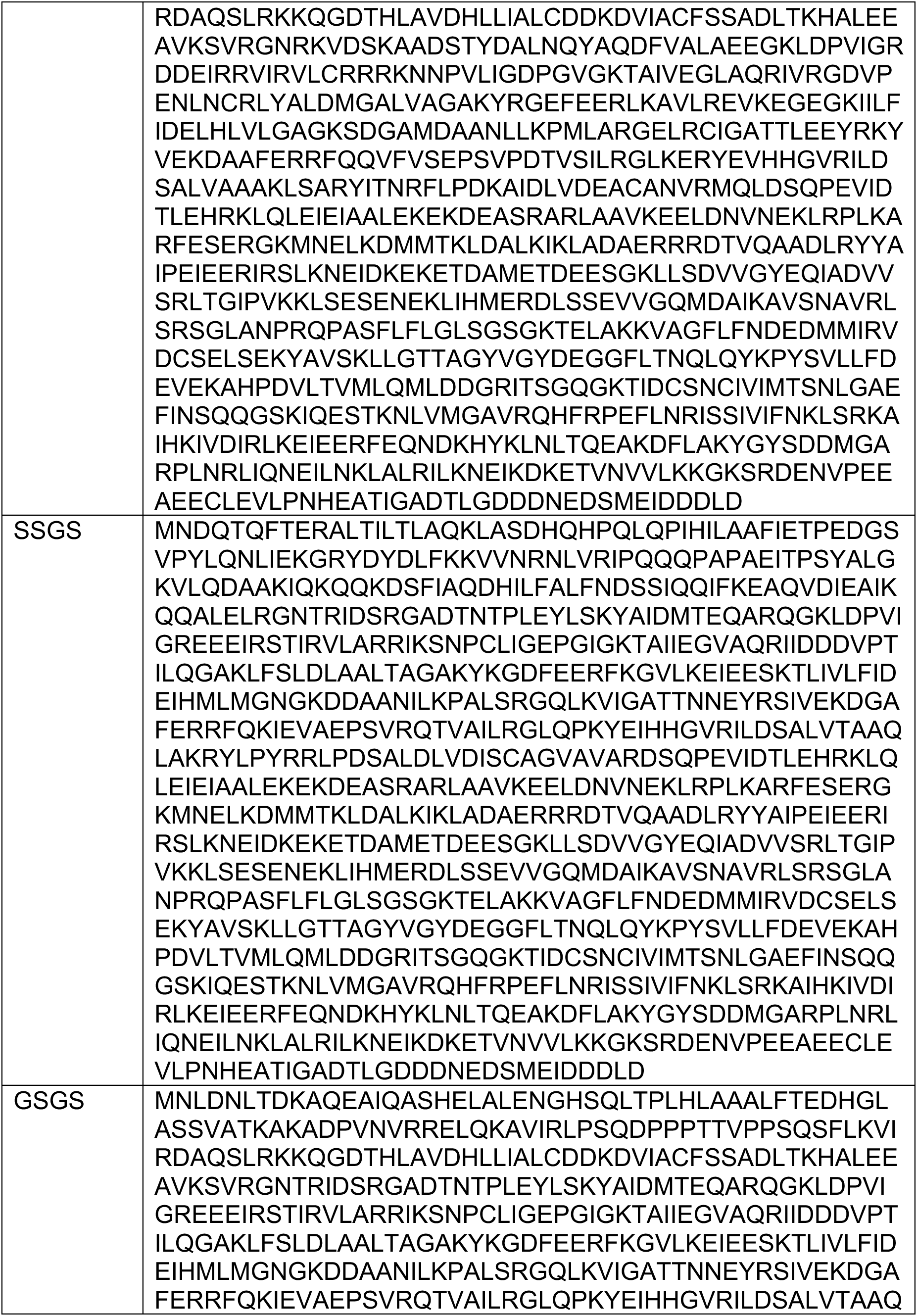
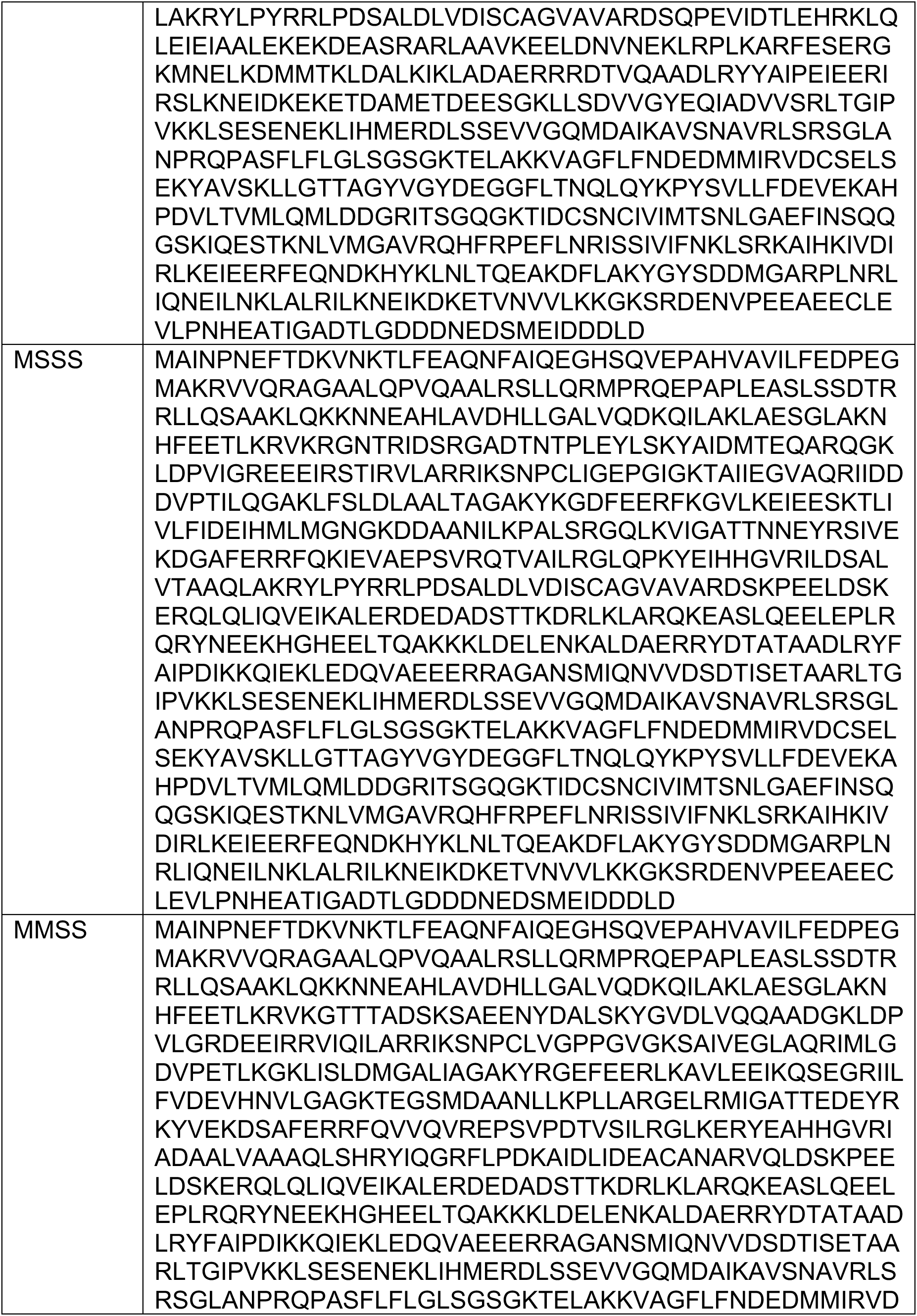
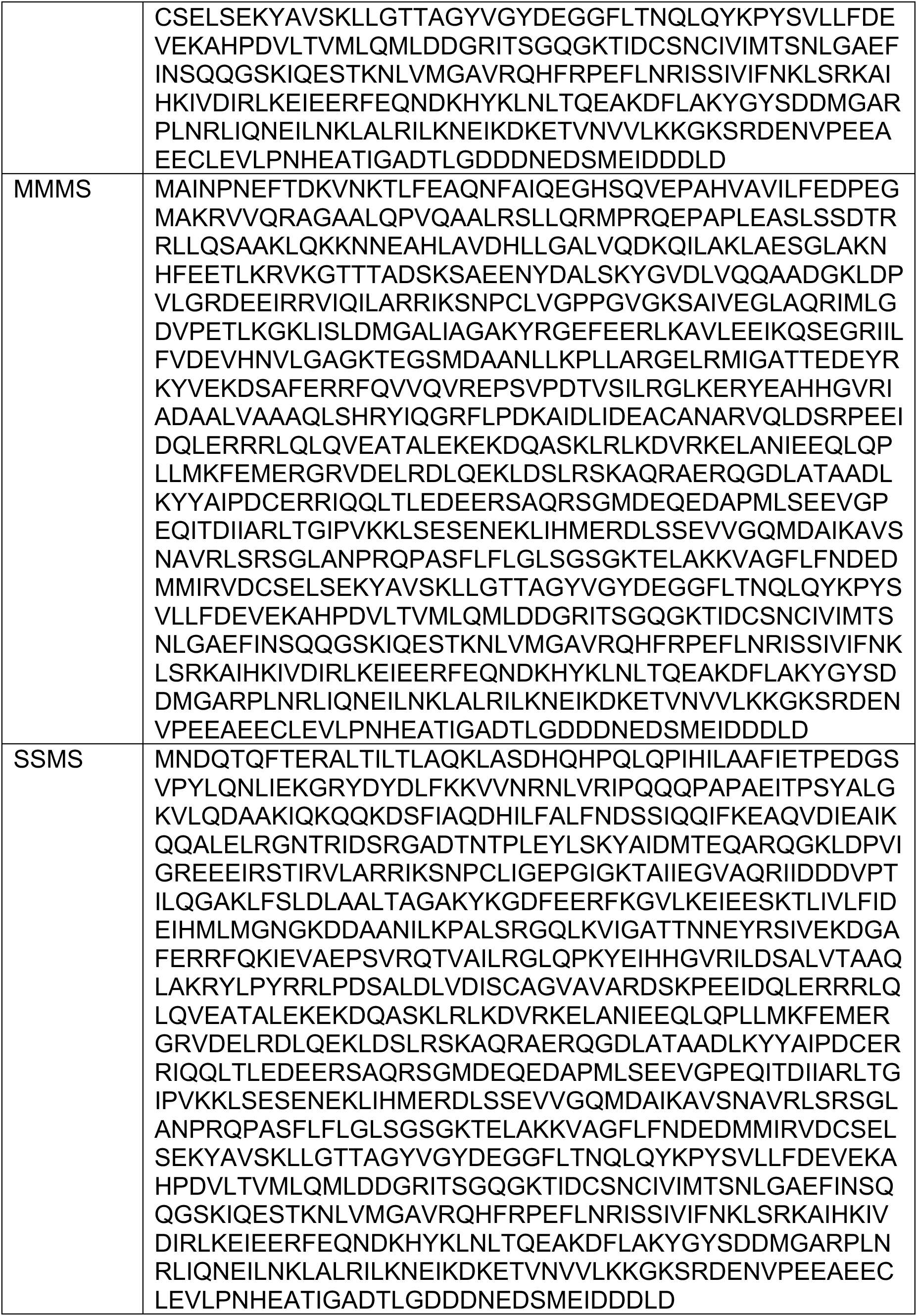
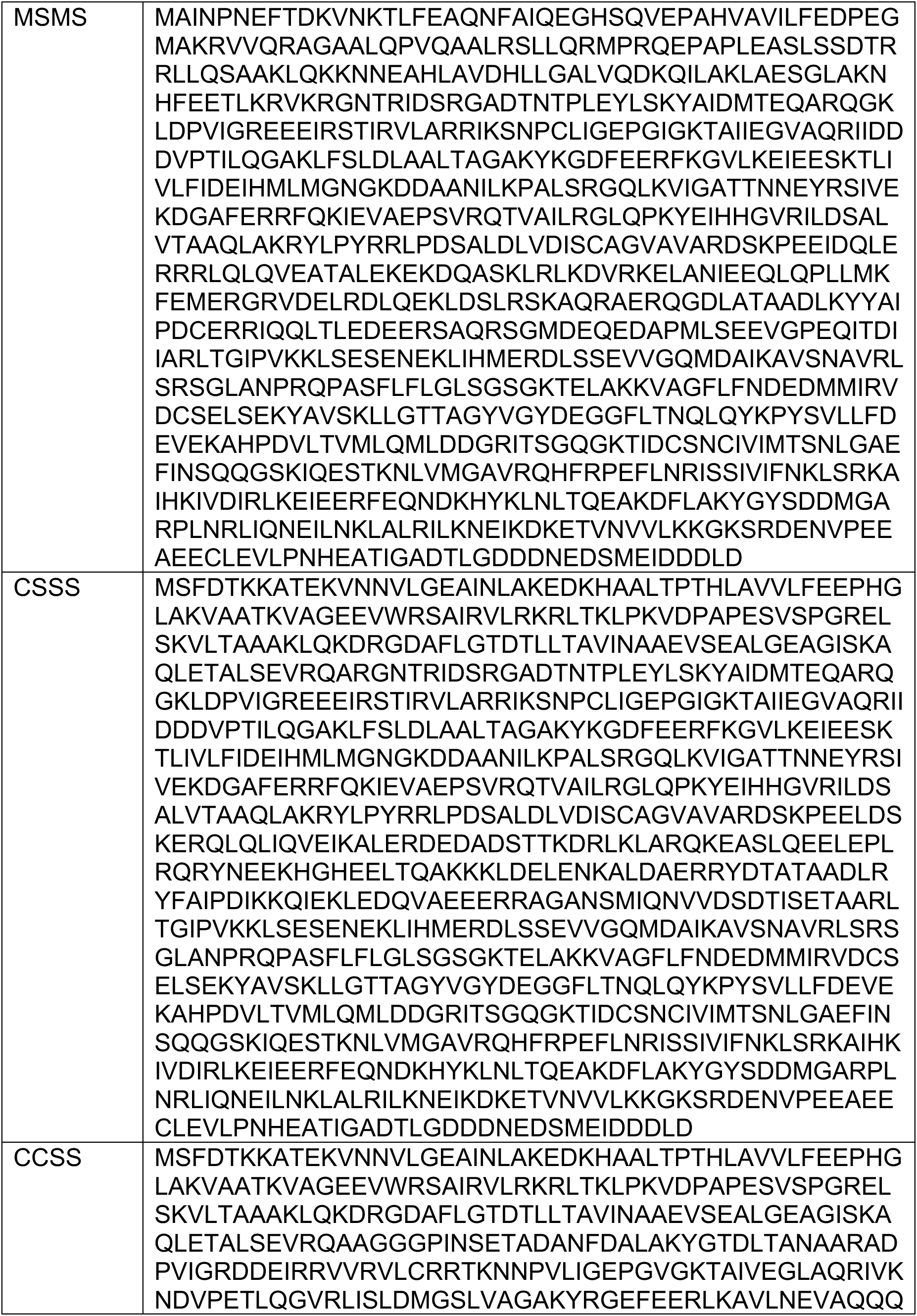
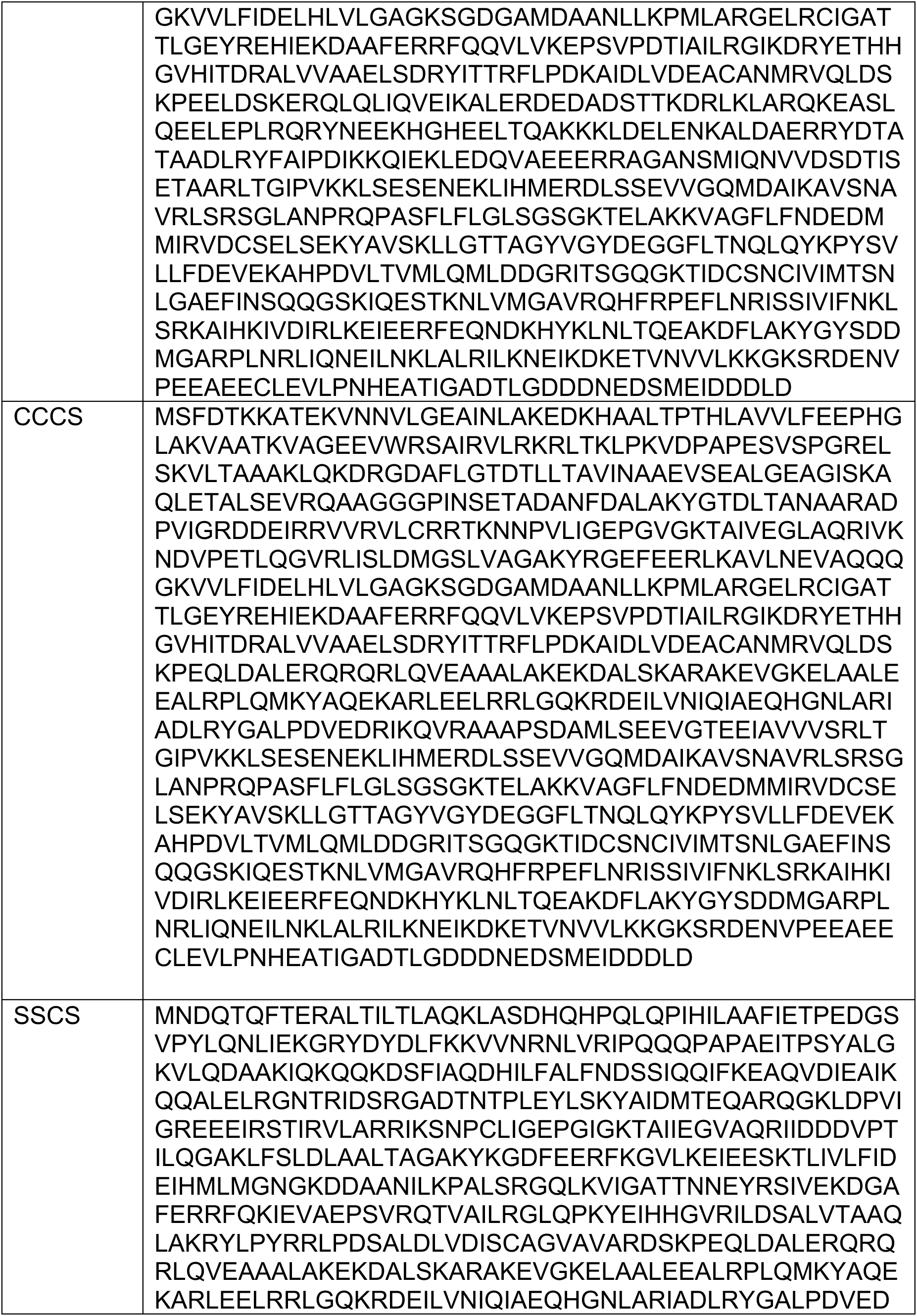
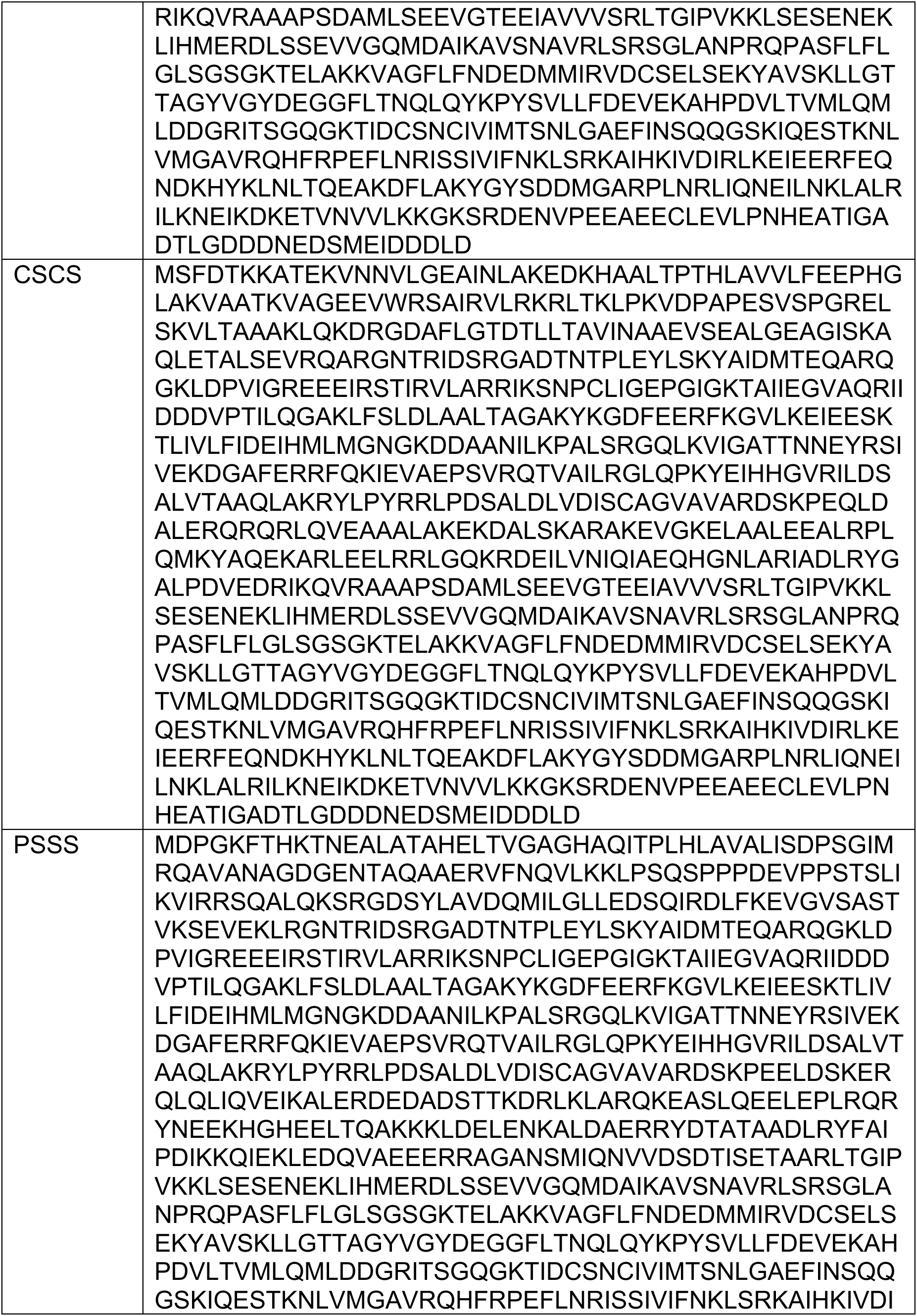
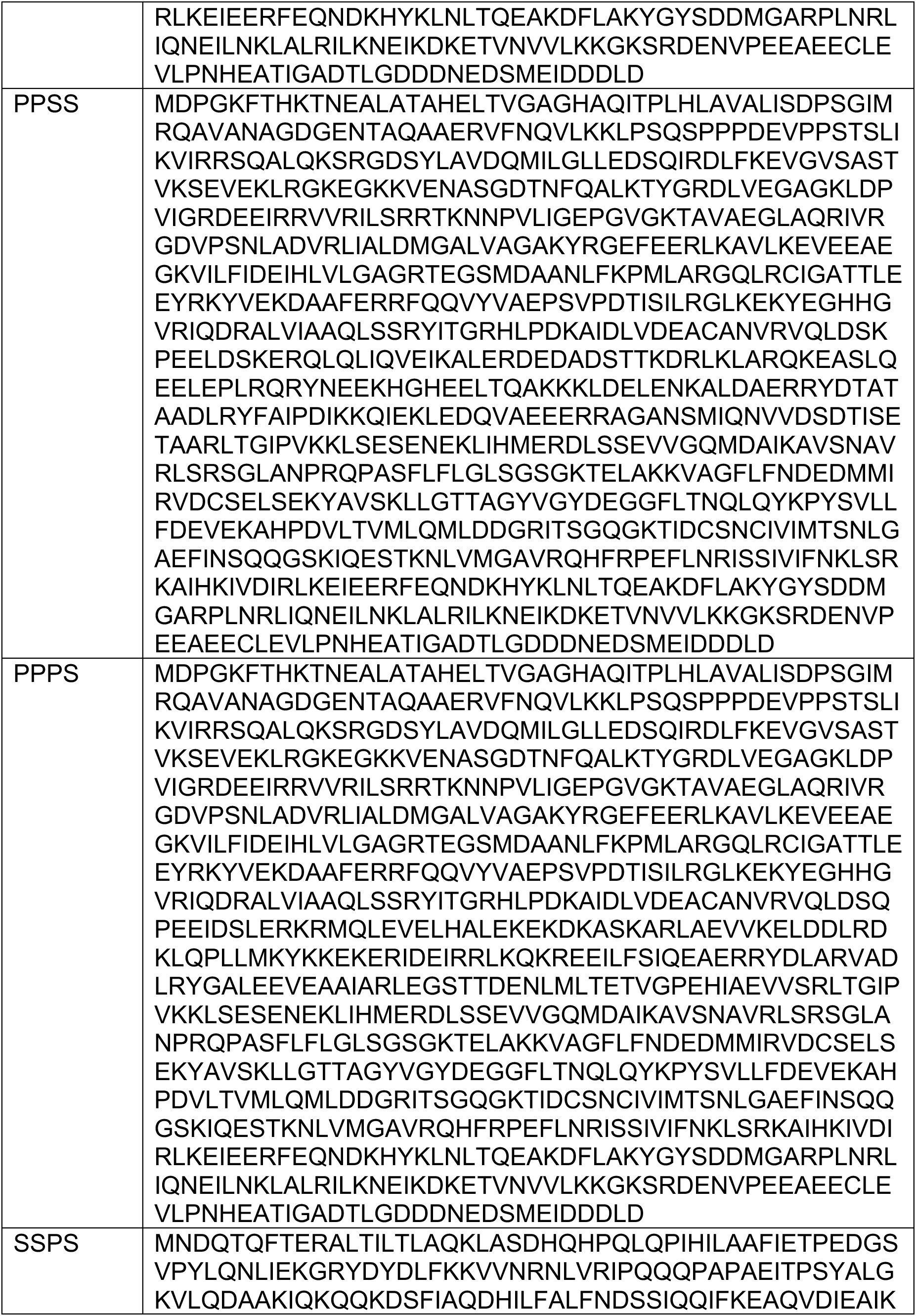
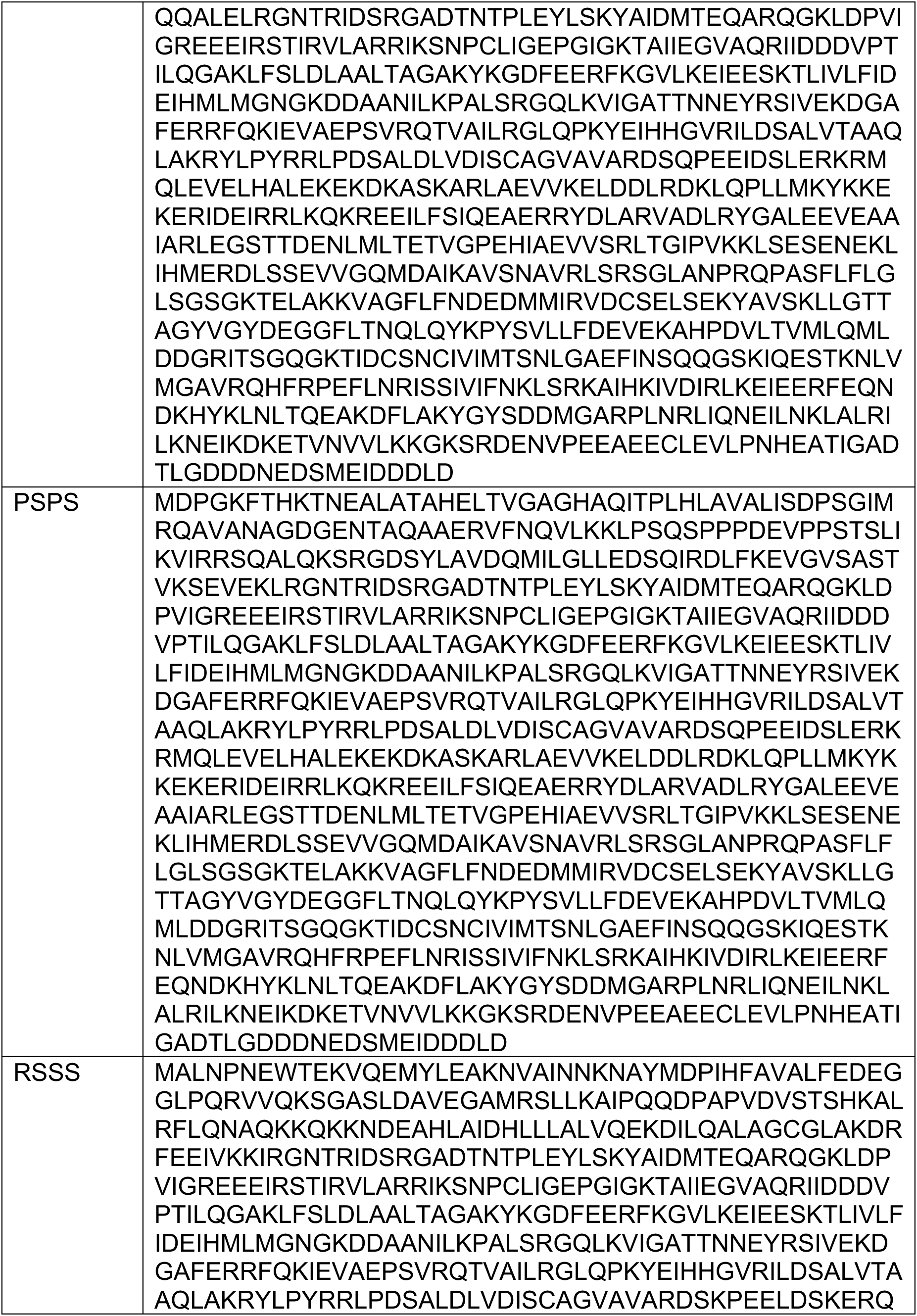
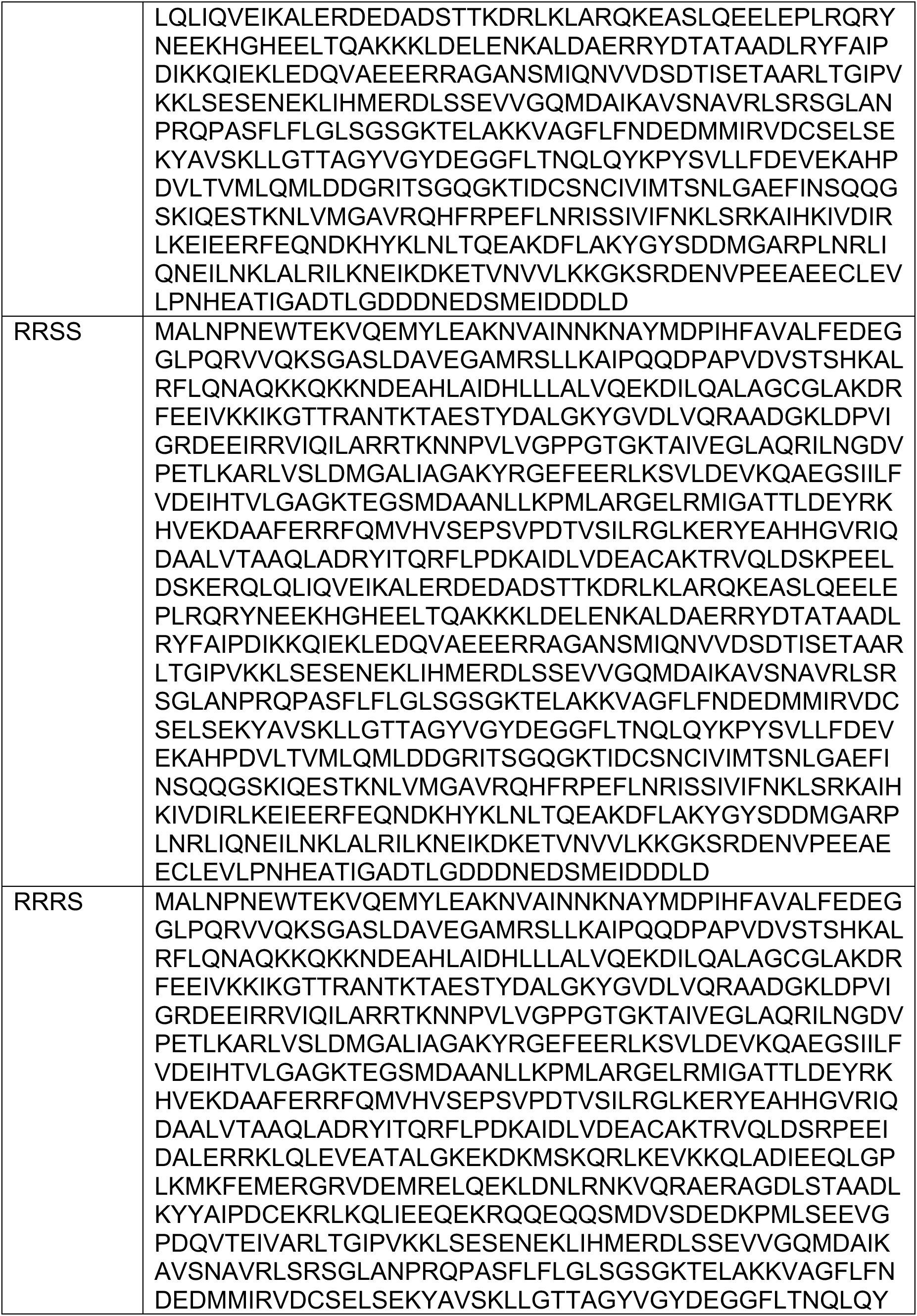
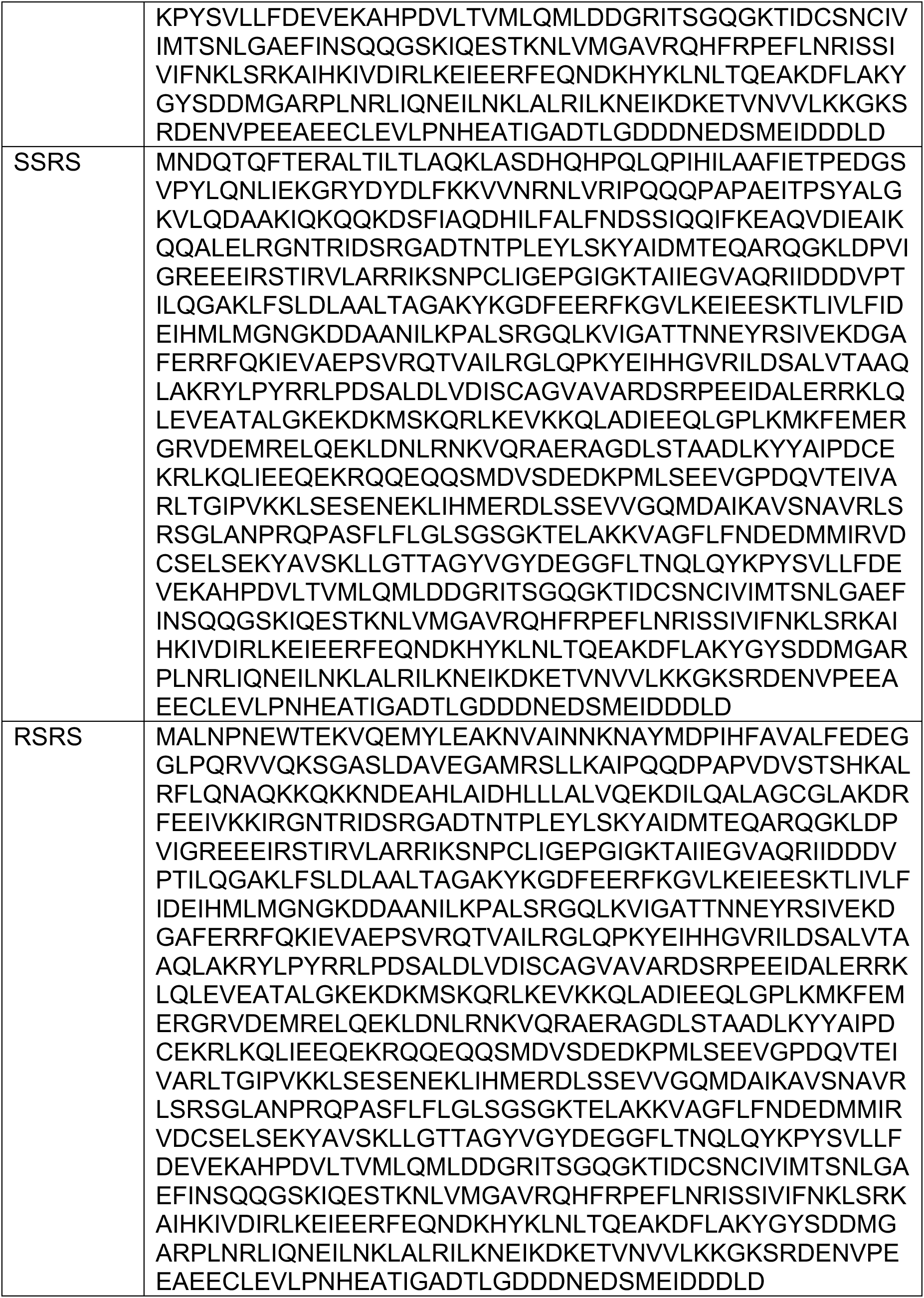
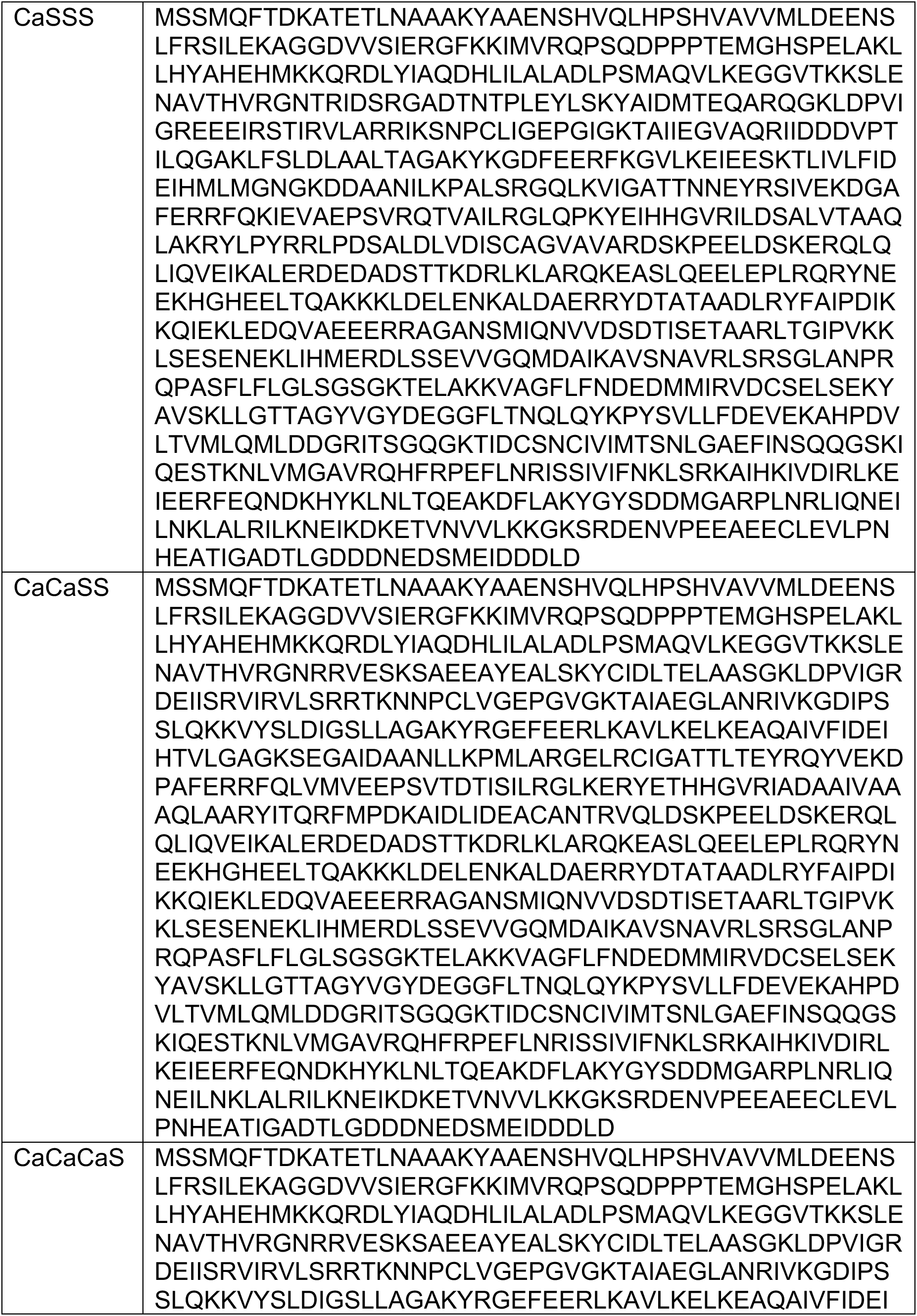
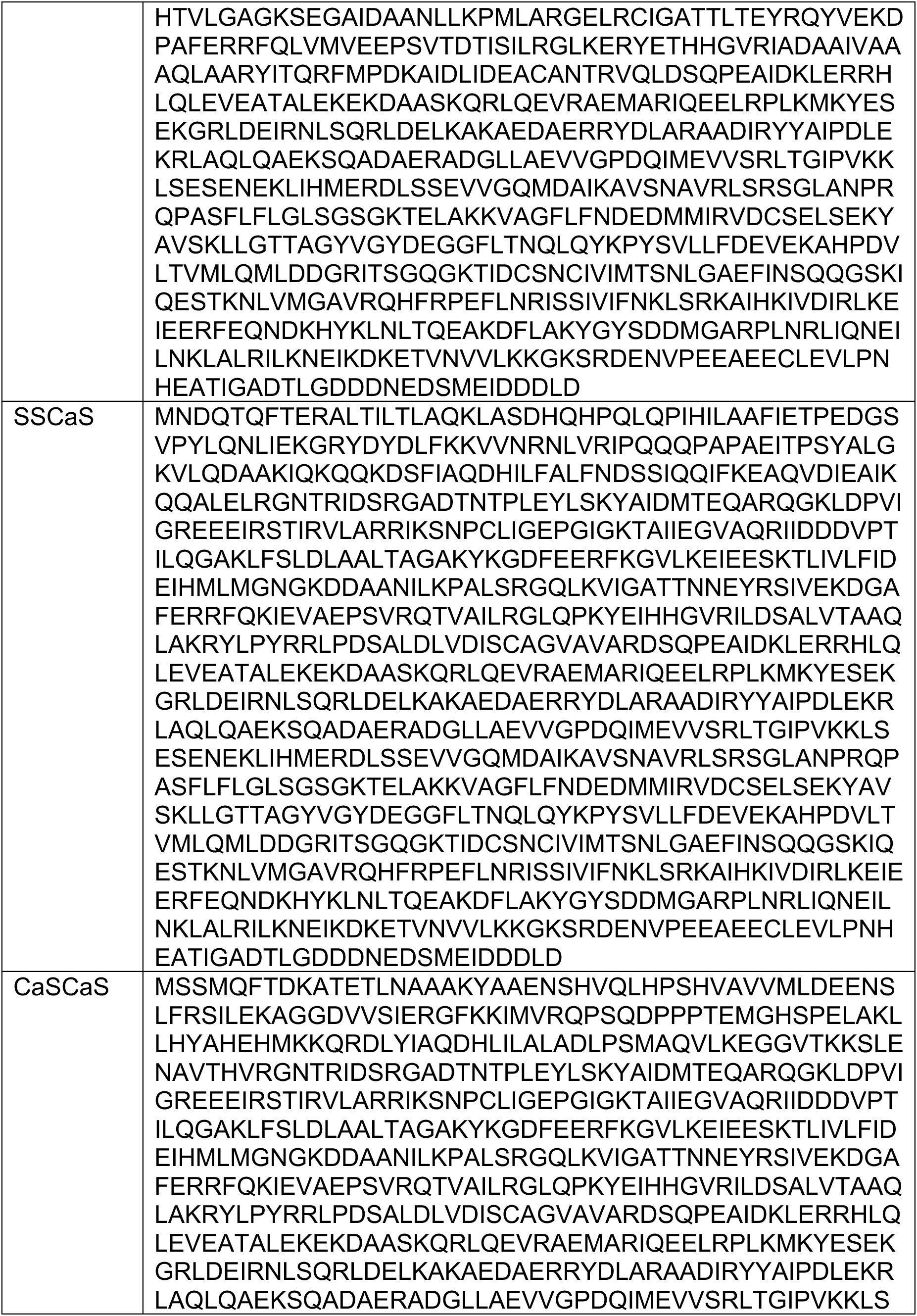
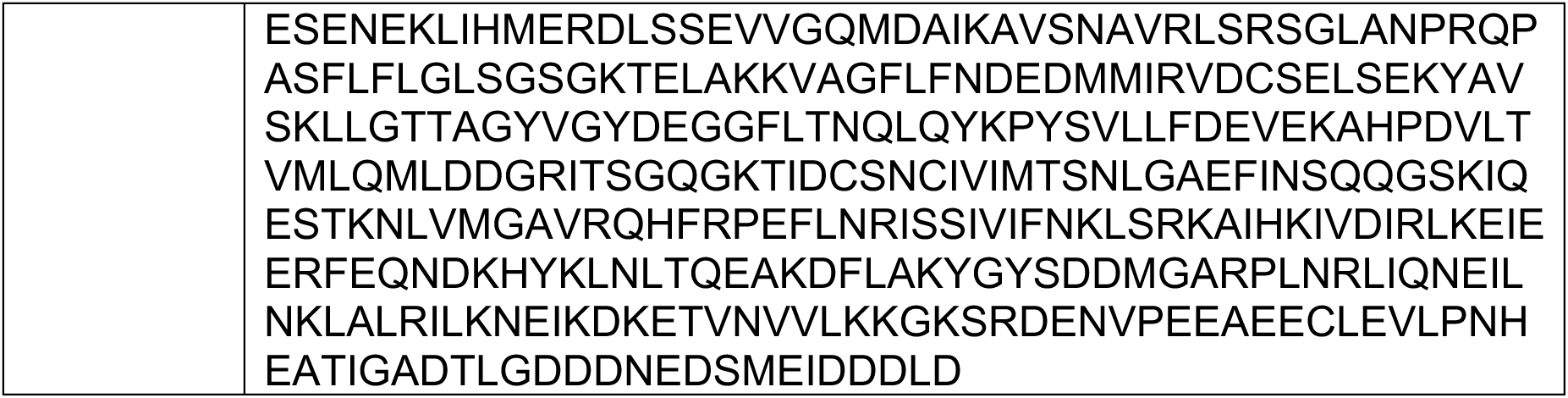
Amino acid sequences of Hsp104 chimeras.

**Table S3.** Pairwise sequence identity between Hsp104 homologs.

|  |  | Total<br>aa | Identical<br>aa | %<br>Identity | Similar<br>aa |
| --- | --- | --- | --- | --- | --- |
|  | *S.cerevisiae* | 908 |  |  |  |
| *S.cerevisiae* | *P.falciparum* | 881 | 319 | 34 | 316 |
| *S.cerevisiae* | *L.thermotolerans* | 907 | 761 | 84 | 105 |
| *S.cerevisiae* | *Ch.thermophilum* | 926 | 467 | 50 | 265 |
| *S.cerevisiae* | *S.thermophilum* | 924 | 463 | 49 | 265 |
| *S.cerevisiae* | *M.thermophila* | 925 | 459 | 49 | 274 |
| *S.cerevisiae* | *T.terrestris* | 923 | 463 | 49 | 262 |
| *S.cerevisiae* | *T.lanuginosus* | 922 | 454 | 48 | 294 |
| *S.cerevisiae* | *T.aurantiacus* | 926 | 458 | 49 | 291 |
| *S.cerevisiae* | *C.thermophila* | 882 | 418 | 46 | 292 |
| *S.cerevisiae* | *C.reinhardtii* | 925 | 383 | 40 | 298 |
| *S.cerevisiae* | *P.euphratica* | 914 | 391 | 42 | 298 |
| *S.cerevisiae* | *A.thaliana* | 911 | 394 | 42 | 294 |
| *S.cerevisiae* | *M.brevicollis* | 889 | 387 | 42 | 303 |
| *S.cerevisiae* | *S.rosetta* | 892 | 381 | 41 | 316 |
| *S.cerevisiae* | *G.sulphuraria* | 922 | 386 | 41 | 332 |
| *S.cerevisiae* | *D.discoideum* | 886 | 401 | 43 | 299 |
| *P.falciparum* | *L.thermotolerans* |  | 302 | 32 | 337 |
| *P.falciparum* | *Ch.thermophilum* |  | 317 | 33 | 331 |
| *P.falciparum* | *S.thermophilum* |  | 316 | 34 | 328 |
| *P.falciparum* | *M.thermophila* |  | 322 | 34 | 321 |
| *P.falciparum* | *T.terrestris* |  | 319 | 34 | 323 |
| *P.falciparum* | *T.lanuginosus* |  | 304 | 32 | 326 |
| *P.falciparum* | *T.aurantiacus* |  | 307 | 32 | 324 |
| *P.falciparum* | *C.thermophila* |  | 306 | 33 | 330 |
| *P.falciparum* | *C.reinhardtii* |  | 318 | 34 | 321 |
| *P.falciparum* | *P.euphratica* |  | 326 | 35 | 334 |
| *P.falciparum* | *A.thaliana* |  | 329 | 35 | 327 |
| *P.falciparum* | *M.brevicollis* |  | 321 | 35 | 332 |
| *P.falciparum* | *S.rosetta* |  | 316 | 35 | 337 |
| *P.falciparum* | *G.sulphuraria* |  | 341 | 36 | 324 |
| *P.falciparum* | *D.discoideum* |  | 328 | 36 | 343 |
| *L.thermotolerans* | *Ch.thermophilum* |  | 469 | 50 | 269 |
| *L.thermotolerans* | *S.thermophilum* |  | 459 | 49 | 279 |
| *L.thermotolerans* | *M.thermophila* |  | 465 | 50 | 273 |
| *L.thermotolerans* | *T.terrestris* |  | 459 | 49 | 272 |
| *L.thermotolerans* | *T.lanuginosus* |  | 448 | 48 | 300 |
| *L.thermotolerans* | *T.aurantiacus* |  | 450 | 48 | 298 |
| *L.thermotolerans* | *C.thermophila* |  | 411 | 45 | 298 |
| *L.thermotolerans* | *C.reinhardtii* |  | 367 | 39 | 315 |
| *L.thermotolerans* | *P.euphratica* |  | 390 | 42 | 310 |
| *L.thermotolerans* | *A.thaliana* |  | 391 | 42 | 302 |
| *L.thermotolerans* | *M.brevicollis* |  | 387 | 42 | 309 |
| *L.thermotolerans* | *S.rosetta* |  | 381 | 41 | 312 |
| *L.thermotolerans* | *G.sulphuraria* |  | 405 | 43 | 314 |
| *L.thermotolerans* | *D.discoideum* |  | 410 | 44 | 295 |
| *Ch.thermophilum* | *S.thermophilum* |  | 831 | 90 | 61 |
| *Ch.thermophilum* | *M.thermophila* |  | 841 | 91 | 54 |
| *Ch.thermophilum* | *T.terrestris* |  | 834 | 90 | 51 |
| *Ch.thermophilum* | *T.lanuginosus* |  | 697 | 75 | 147 |
| *Ch.thermophilum* | *T.aurantiacus* |  | 692 | 74 | 142 |
| *Ch.thermophilum* | *C.thermophila* |  | 486 | 52 | 258 |
| *Ch.thermophilum* | *C.reinhardtii* |  | 420 | 43 | 283 |
| *Ch.thermophilum* | *P.euphratica* |  | 452 | 48 | 271 |
| *Ch.thermophilum* | *A.thaliana* |  | 449 | 47 | 278 |
| *Ch.thermophilum* | *M.brevicollis* |  | 438 | 47 | 278 |
| *Ch.thermophilum* | *S.rosetta* |  | 430 | 46 | 280 |
| *Ch.thermophilum* | *G.sulphuraria* |  | 460 | 48 | 276 |
| *Ch.thermophilum* | *D.discoideum* |  | 446 | 47 | 278 |
| *S.thermophilum* | *M.thermophila* |  | 850 | 92 | 63 |
| *S.thermophilum* | *T.terrestris* |  | 840 | 91 | 60 |
| *S.thermophilum* | *T.lanuginosus* |  | 697 | 75 | 147 |
| *S.thermophilum* | *T.aurantiacus* |  | 703 | 76 | 132 |
| *S.thermophilum* | *C.thermophila* |  | 483 | 52 | 252 |
| *S.thermophilum* | *C.reinhardtii* |  | 424 | 44 | 281 |
| *S.thermophilum* | *P.euphratica* |  | 445 | 47 | 288 |
| *S.thermophilum* | *A.thaliana* |  | 458 | 48 | 273 |
| *S.thermophilum* | *M.brevicollis* |  | 440 | 48 | 276 |
| *S.thermophilum* | *S.rosetta* |  | 431 | 46 | 278 |
| *S.thermophilum* | *G.sulphuraria* |  | 463 | 49 | 278 |
| *S.thermophilum* | *D.discoideum* |  | 447 | 48 | 284 |
| *M.thermophila* | *T.terrestris* |  | 848 | 91 | 58 |
| *M.thermophila* | *T.lanuginosus* |  | 701 | 76 | 145 |
| *M.thermophila* | *T.aurantiacus* |  | 693 | 75 | 146 |
| *M.thermophila* | *C.thermophila* |  | 485 | 52 | 254 |
| *M.thermophila* | *C.reinhardtii* |  | 423 | 44 | 286 |
| *M.thermophila* | *P.euphratica* |  | 445 | 47 | 271 |
| *M.thermophila* | *A.thaliana* |  | 458 | 48 | 268 |
| *M.thermophila* | *M.brevicollis* |  | 439 | 47 | 280 |
| *M.thermophila* | *S.rosetta* |  | 432 | 46 | 275 |
| *M.thermophila* | *G.sulphuraria* |  | 457 | 48 | 282 |
| *M.thermophila* | *D.discoideum* |  | 451 | 48 | 281 |
| *T.terrestris* | *T.lanuginosus* |  | 698 | 75 | 143 |
| *T.terrestris* | *T.aurantiacus* |  | 695 | 75 | 141 |
| *T.terrestris* | *C.thermophila* |  | 483 | 52 | 251 |
| *T.terrestris* | *C.reinhardtii* |  | 427 | 45 | 280 |
| *T.terrestris* | *P.euphratica* |  | 447 | 47 | 277 |
| *T.terrestris* | *A.thaliana* |  | 454 | 48 | 274 |
| *T.terrestris* | *M.brevicollis* |  | 443 | 48 | 271 |
| *T.terrestris* | *S.rosetta* |  | 427 | 46 | 280 |
| *T.terrestris* | *G.sulphuraria* |  | 458 | 48 | 284 |
| *T.terrestris* | *D.discoideum* |  | 444 | 47 | 279 |
| *T.lanuginosus* | *T.aurantiacus* |  | 794 | 85 | 101 |
| *T.lanuginosus* | *C.thermophila* |  | 482 | 52 | 256 |
| *T.lanuginosus* | *C.reinhardtii* |  | 418 | 43 | 276 |
| *T.lanuginosus* | *P.euphratica* |  | 437 | 46 | 275 |
| *T.lanuginosus* | *A.thaliana* |  | 447 | 47 | 265 |
| *T.lanuginosus* | *M.brevicollis* |  | 437 | 47 | 277 |
| *T.lanuginosus* | *S.rosetta* |  | 420 | 25 | 290 |
| *T.lanuginosus* | *G.sulphuraria* |  | 460 | 49 | 281 |
| *T.lanuginosus* | *D.discoideum* |  | 444 | 48 | 292 |
| *T.aurantiacus* | *C.thermophila* |  | 475 | 51 | 261 |
| *T.aurantiacus* | *C.reinhardtii* |  | 424 | 44 | 271 |
| *T.aurantiacus* | *P.euphratica* |  | 444 | 47 | 270 |
| *T.aurantiacus* | *A.thaliana* |  | 446 | 47 | 267 |
| *T.aurantiacus* | *M.brevicollis* |  | 419 | 45 | 293 |
| *T.aurantiacus* | *S.rosetta* |  | 417 | 45 | 293 |
| *T.aurantiacus* | *G.sulphuraria* |  | 456 | 48 | 281 |
| *T.aurantiacus* | *D.discoideum* |  | 439 | 47 | 300 |
| *C.thermophila* | *C.reinhardtii* |  | 454 | 49 | 257 |
| *C.thermophila* | *P.euphratica* |  | 463 | 50 | 257 |
| *C.thermophila* | *A.thaliana* |  | 464 | 50 | 253 |
| *C.thermophila* | *M.brevicollis* |  | 468 | 52 | 258 |
| *C.thermophila* | *S.rosetta* |  | 467 | 52 | 259 |
| *C.thermophila* | *G.sulphuraria* |  | 479 | 52 | 264 |
| *C.thermophila* | *D.discoideum* |  | 494 | 55 | 263 |
| *C.reinhardtii* | *P.euphratica* |  | 526 | 56 | 236 |
| *C.reinhardtii* | *A.thaliana* |  | 523 | 56 | 242 |
| *C.reinhardtii* | *M.brevicollis* |  | 487 | 52 | 248 |
| *C.reinhardtii* | *S.rosetta* |  | 480 | 51 | 245 |
| *C.reinhardtii* | *G.sulphuraria* |  | 503 | 53 | 236 |
| *C.reinhardtii* | *D.discoideum* |  | 491 | 53 | 248 |
| *P.euphratica* | *A.thaliana* |  | 791 | 86 | 97 |
| *P.euphratica* | *M.brevicollis* |  | 500 | 54 | 252 |
| *P.euphratica* | *S.rosetta* |  | 487 | 52 | 261 |
| *P.euphratica* | *G.sulphuraria* |  | 526 | 55 | 242 |
| *P.euphratica* | *D.discoideum* |  | 525 | 57 | 235 |
| *A.thaliana* | *M.brevicollis* |  | 507 | 54 | 236 |
| *A.thaliana* | *S.rosetta* |  | 500 | 54 | 255 |
| *A.thaliana* | *G.sulphuraria* |  | 527 | 56 | 242 |
| *A.thaliana* | *D.discoideum* |  | 527 | 57 | 231 |
| *M.brevicollis* | *S.rosetta* |  | 634 | 71 | 176 |
| *M.brevicollis* | *G.sulphuraria* |  | 528 | 57 | 225 |
| *M.brevicollis* | *D.discoideum* |  | 512 | 57 | 255 |
| *S.rosetta* | *G.sulphuraria* |  | 517 | 56 | 240 |
| *S.rosetta* | *D.discoideum* |  | 499 | 56 | 282 |
| *G.sulphuraria* | *D.discoideum* |  | 538 | 58 | 240 |

